# Investigating Alternative Models of Acute HIV Infection

**DOI:** 10.1101/2023.06.04.543605

**Authors:** Ellie Mainou, Ruy Ribeiro, Jessica M Conway

## Abstract

Understanding the dynamics to acute HIV infection may provide insights into the mechanisms of early viral control with potential implications for vaccine design. The standard viral dynamics model explains HIV viral dynamics during acute infection reasonably well. However, the model makes simplifying assumptions, neglecting some aspects of HIV. For example, in the standard model, target cells are infected by a single HIV virion. Yet, cellular multiplicity of infection (MOI) may have considerable effects in pathogenesis and viral evolution. Further, when using the standard model, we take constant infected cell death rates, simplifying the dynamic immune responses. Here, we use four models—1) the standard viral dynamics model, 2) an alternate model incorporating cellular MOI, 3) a model assuming density-dependent death rate of infected cells and 4) a model combining (2) and (3)—to investigate acute infection dynamics in 43 people tested very early after HIV exposure. We find that all models explain the data, but different models describe differing features of the dynamics more accurately. For example, while the standard viral dynamics model may be the most parsimonious model, viral peaks are better explained by a model allowing for cellular MOI. These results suggest that heterogeneity in within-host viral dynamics cannot be captured by a single model. Thus depending on the aspect of interest, a corresponding model should be employed.

## 1 Introduction

HIV remains a global health challenge, with 38.4 million people living with HIV in 2021 [1]. To control HIV incidence, a successful vaccine is needed. However, candidate vaccines tested so far have failed or provide modest efficacy [2]. Elucidating the key events that occur during acute infection may facilitate the development of effective vaccines. Here, we aim to better characterize the dynamics of early HIV-1 infection by testing four different mechanistic hypotheses; specifically, we aim to investigate the effects of cellular coinfection and of the adaptive immune responses on acute HIV infection dynamics.

Acute HIV infection covers a period of 4–5 weeks in which the virus disseminates from the initial site of infection into various tissues and organs [3]. Primary infection kinetics are characterized by exponential increase in the number of virus particles in peripheral blood, reaching a peak, followed by a decline to steady state level, which is referred to as the viral setpoint [4–6]. The viral peak coincides with the first appearance of an adaptive immune response [7]. The decline in plasma viremia is attributed to adaptive immune responses [7], and/or to target cell limitation [8, 9] (Figure 1).

**Figure 1:**
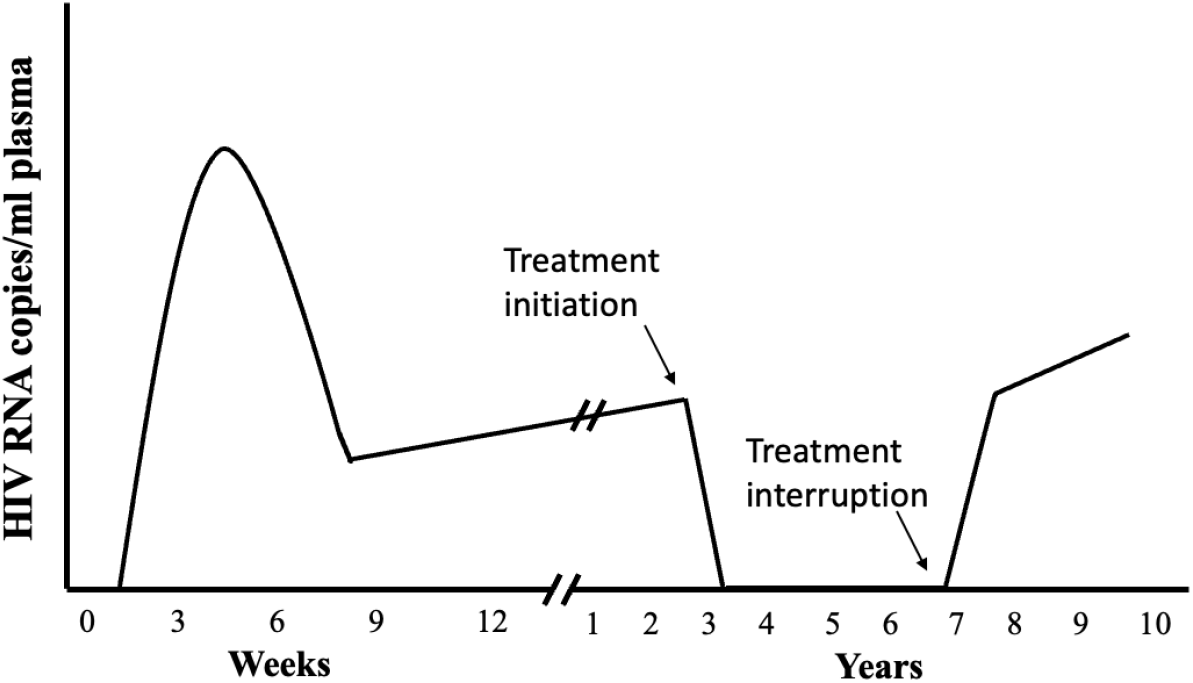
HIV within-host dynamics.Within weeks from seroconversion, HIV viral load increases rapidly, reaches a peak and decreases to an almost stable viral load. Treatment is effective in decreasing viral load to undetectable levels. Treatment interruption leads to rebound viremia to pre-therapy levels.

Mathematical models have helped to better understand the processes driving within-host dynamics of viral infections.In the case of HIV, acute infection has been modeled with a standard model of viral dynamics initially developed to explore viral decay during treatment [10]. The model consists of uninfected target cells (CD4+ T cells), infected cells and free plasma virus. A central assumption of the model is that target cells are either uninfected or infected with a single virion. Statistical fitting of this model to viral load measurements has yielded estimates for the within-host basic reproductive number *R*_0_ [11], as well as estimates for key parameters [9, 10, 12–16]. Overall, the standard model has been able to capture viral loads in people with HIV early in infection and provide insight into viral kinetics [8–10, 13]. However, fits to viral load measurements underestimates peak viremia and produce oscillatory dynamics during the setpoint [17]. In addition, certain important assumptions are made. First, the model assumed a constant, log-linear decay of the infected cell population. While this assumption may be valid for the early stages of the infection, it might not be realistic for longer periods of months or years. As the infection is established, the increased presence of infected cells may stimulate immune responses that could lead to an increased killing of infected cells. Holte et al. (2006) incorporated this feature to the standard model. Fits to this model are able to explain HIV infection dynamics [18]. Reeves et al. (2021) fit an ensemble of models to the to a rich data set of acute infection (RV217 data set) [19] using a population non-linear mixed-effects framework and found that the model developed by Holte et al. provides the best fit at the population level [20]. It should be noted that that this model does not incorporate explicitly adaptive immune responses.

A potentially important infection mechanism not included in the standard model is cellular multiplicity of infection, which may have considerable effects in pathogenesis and evolution. Cellular coinfection can allow for the generation of recombinant viruses. For example, Levy et al. (2004) found that a single round of replication in T lymphocytes in culture generated an average of nine recombination events per virus [21]. Genetic recombination may assist HIV-1 diversification and escape from both antiviral therapies and host immune responses [21]. The effects of cellular MOI have been poorly explored in mathematical models of viral infectious diseases [22]. Dixit & Perelson (2004 and 2005) developed a model for HIV cellular coinfection [17, 23]. A central assumption of this model is that viral production remains constant across infected cells, regardless of multiplicity of infection. Under this scenario, predictions of this model are identical to the standard viral dynamics model. However, if multiply infected cells are characterized by an increased burst size, predictions are altered. In this case, the establishment of infection may not be dependent on the basic reproduction number only, but also on the initial virus load. In addition, the viral population may not grow exponentially at a constant rate, but in an increasing rate as viral load increases [24]. Whether burst size is constant across different classes of infected cells remains unknown [21]. Wodarz & Levy (2011) and Guo et al. (2020) built upon to Dixit & Perelson (2005) and developed a system of ODEs, where each class of infected cells is represented by an equation. These models were used to make theoretical predictions and were not fit to data, to our knowledge. HIV-1 models that account for multiplicity of infection remain high-dimensional and difficult to be validated with observed viral load datasets (see Discussion section).

To account for cellular coinfection, without resorting to a high-dimensional system, Koelle et al. (2019) developed a new class of low-dimensional within-host models whose structure flexibly allows for cellular coinfection. Their model is based on the structure of epidemiological macroparasite models [25–27], where target cells are analogous to hosts and virions are analogous to macroparasites. In this model, target cells, either uninfected or (singly or multiply) infected are represented by a single equation, dramatically reducing the dimensionality [22]. The model was fit to an influenza viral load data set and was found to perform better than standard influenza viral dynamics model. In addition, it was able to capture the peak viremia, which the standard viral dynamics model predictions with influenza data underestimates [22].

Here, we aim to better characterize the dynamics of early HIV-1 infection by testing different mechanistic hypotheses expressed as simple models, to investigate the effect of adaptive immune responses and cellular coinfection in acute HIV infection dynamics. We are interested in using simple models. We use the standard viral dynamics model, the Density-Dependent Cell Death model developed by Holte et al. (2006) [18] and we adapt the MOI model [22] to the dynamics of HIV-1. Finally, we develop a new model that incorporates both multiplicity of infection and the

Density-Dependent Cell Death of infected cells. We fit these four models to viral load data of 43 participants in a prospective study of acute HIV infection [19]. We fit each study participant viral loads individually. Because we focus on early infection dynamics, data points for each study participant are limited. Therefore, fitting complex models to a few data would be extremely difficult, but these simple models remain appropriate. For this same reason, we do not test models that incorporate adaptive immune responses explicitly; such models would add a number of parameters and our dataset would not be sufficient for model fitting.

In the following, we perform model selection to assess the overall performance of the models, and we also test their ability to predict certain viral quantitative measures, namely the growth and decay rates, the viral setpoint, the peak magnitude, the peak timing and a joint measurement that accounts simultaneously for both the magnitude and the timing of peak viremia. We find that overall, the Density-Dependent Cell Death model describes acute HIV infection dynamics best. However, different models explain differing quantitative features of the dynamics more accurately, as we will demonstrate. We finish by discussing the effect of adaptive immune responses on determining the level of the viral setpoint.

## 2 Methods

### 2.1 The Data

Viral load measurements were obtained from the RV217 study [19]. Participants were recruited in four locations (Uganda, Kenya, Tanzania and Thailand) between June 2009 and June 2015. Study participants underwent twice-weekly small-volume blood collection by fingerstick and large-volume blood collections every 6 months. Once tests for HIV-1 RNA were reactive, large-volume blood samples were collected twice weekly for 4 weeks. Participants with confirmed HIV-1 infection were enrolled in the long-term follow-up phase. Analysis is restricted to 43 study participants in whom at least two large-volume blood sample showed detectable HIV-1 RNA and a nonreactive enzyme immunoassay, who had at least one study visit before viral RNA detection and who had quantitative HIV-1 RNA data. We include study participants for whom at least two viral load measurements were taken before inferred peak viremia. Day 0 is defined as the day of the first positive nucleic acid test. We aim to characterize dynamics of acute infection and thus restrict viral load measurements to the second viral setpoint measurement for each study participant.

#### 2.1.1 Data Analysis

We characterize viral quantitative measures for each study participant’s viral load curve. These measures include 1) the growth rate, 2) the decay rate, 3) the viral setpoint, 4) the peak viremia, 5) the timing of viral peak, and 6) a joint measurement of the the peak magnitude and timing.

We calculate the growth and decay rates and setpoint for each study participant in the following way: we categorize data points as belonging to the growth phase, decay phase or setpoint. We then fit linear regressions to the growth and decay data points. The slopes of these regressions are the growth and decay rates. For setpoint, we fit a horizontal line (Figure 2 for an illustration).

**Figure 2:**
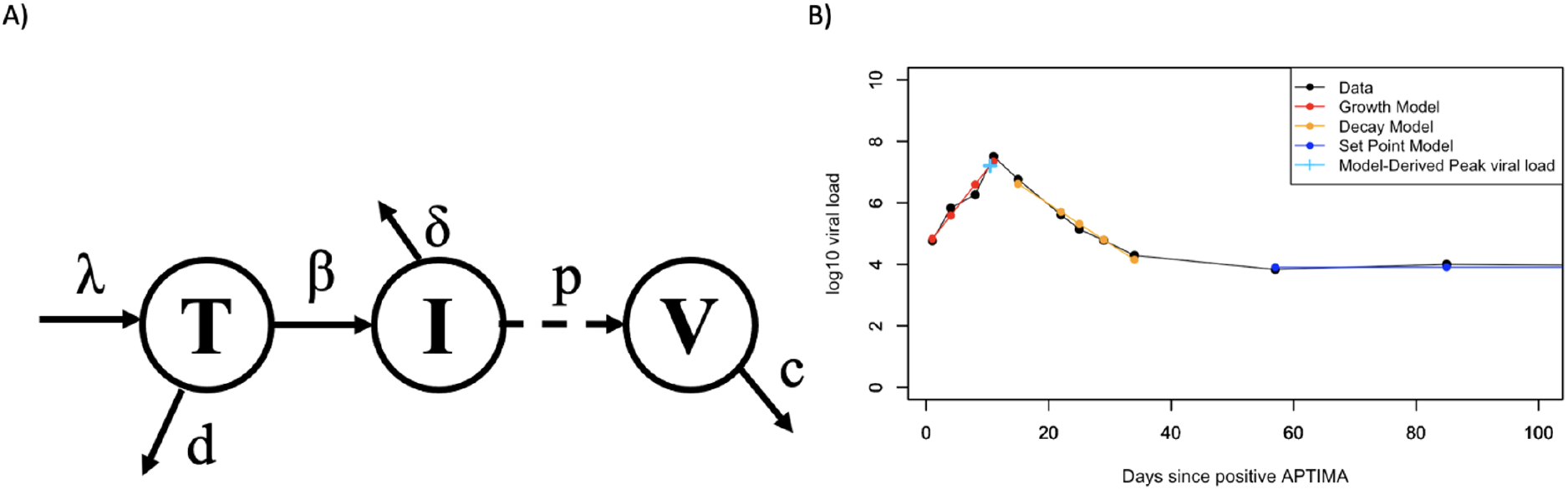
A) Figure schematic of the standard model. Target cells, *T* are sourced at rate *λ* and die at rate *d*. Virus *V* infects target cells at rate *β*. Infected cells, *I* die at rate *δ* and produce virions at rate *p*. Free virus gets cleared at rate *c*. B) Example of how quantitative measures are estimated for each study participants. Black points represent viral load measurements for study participant 10066. Red, orange and blue points represent the predicted growth, decay and setpoint from the linear regression model. Light blue cross represents the intersection of the growth and decay lines.

Peak magnitude is estimated as the maximum value between the observed maximum viral load and the intersection of the growth and decay lines. Peak timing is calculated as the time when the peak magnitude occurs. Finally, to account for both magnitude and timing of the peak we derive the a joint measurement that incorporates both measures:

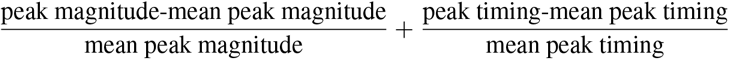

Estimated quantitative measures are reported in Supplementary Table S1.

### 2.2 The Models

To investigate infection dynamics among study participants, we use four models:

#### 1. Standard model

The standard viral dynamics model of acute infection was developed to provide the first estimates of fundamental within-host parameters, such as the viral clearance rate and the infected cell lifespan [13] and to explore viral decay during treatment [10]. The standard model is a target cell-limited model, i.e. HIV infection is limited by the availability of target cells, consisting of three compartments–uninfected cells, infected cells and free virus. Target cells, *T*, are produced at rate *λ*, die at rate *d* and become infected at rate *k*. Infected cells, *I*, die at a constant rate *δ* and produce HIV-1 virions at rate *p*. Free virus, *V*, is cleared from the blood at rate *c* (Figure 2). This model can be summarized in the following system of differential equations:

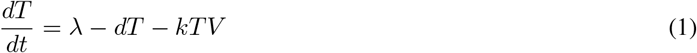

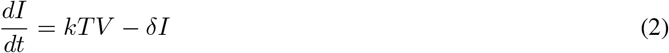

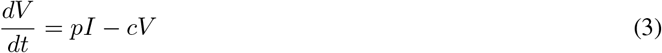

This target-cell limited model has been shown to capture the dynamics of early infection, however it produces oscillatory dynamics and tends to underestimate the magnitude of peak viremia [9].

#### 2. Density-dependent death of infected cells

An alternative model proposed by Holte et al. [18] was developed to explain the biphasic decay of HIV virus after treatment initiation. This model allows for nonlinear log decay of infected cell populations, by incorporating adaptive immune responses. The model equations are identical to those of the standard viral dynamics model, with the exception of *δI*, which now becomes *δI*^*γ*^. *γ* represents density-dependence in the death of infected cells. Note that for *γ* = 1, this model reduces to the standard viral dynamics model.

#### 3. MOI model

Both of the aforementioned models do not make specific assumptions about multiple cellular infections. In the case of HIV, however, there is evidence for cellular coinfection, both *in vivo* and *in vitro* [21, 28–30]. Previous models developed to allow for cellular coinfection are high-dimensional [31] and therefore complicated to ft to available data convincingly. To allow for cellular multiplicity of infection, without resorting to a high-dimensional system, Koelle et al. developed a within-host model that is a close analogy to the population level macroparasite models [22]. They used it to investigate within-host influenza dynamics and found an improved characterization of overall dynamics, as well as of peak viremia, motivating us to test this simple model as well. This is again a target cell-limited model and takes on the following general form:

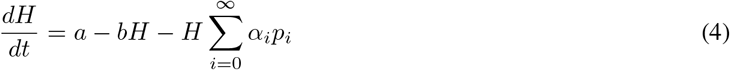

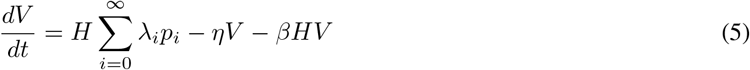

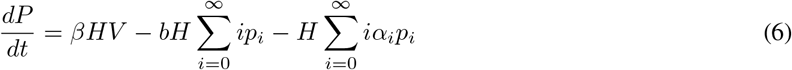

*H* represents the total number of target cells. This includes both uninfected and infected cells with a variable number of virions. Both unifected and infected cells are targets to further infection. Target cells are produced at a constant rate *a* and die at rate *b*. 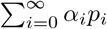 is the infection-induced mortality of target cells. *α*_*i*_ is the death rate of target cells infected with *i* virions and *p*_*i*_ is the proportion of target cells that are infected with cellular MOI of *i*. The variable *V* is the concentration of free virus. 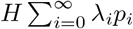 represents the viral production, where *λ*_*i*_ is the viral production rate of cells infected with MOI *i*. Free virus is cleared at rate *η* and is lost due to entry into target cells at rate *β. P* represents the amount of internalized virus, across all target cells. *βHV* captures the increase in internalized virus due to cell entry of free virus. 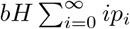 represents the death of internalized virus due to the background mortality of target cells and 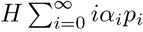 represents the deaths of internalized virus due to the infection-induced mortality of target cells. In the absence of data supporting a more complex formulation, we make the simple assumption that viral production rates and infection-induced mortality rates scale linearly with MOI (*λ*_*i*_ = *i · λ, α*_*i*_ = *i · α*), and that MOI follows a binomial distribution (Koelle et al. 2019), the system becomes the following:

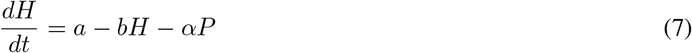

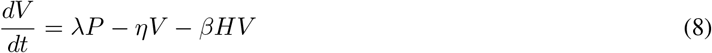

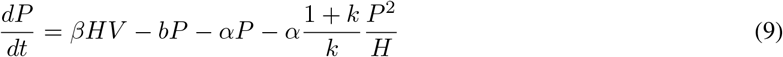

where *k* is the dispersion parameter of the negative binomial distribution (Koelle et al 2019).

#### 4. Density-Dependent Death of infected cells & MOI

By comparing a density-dependent infected cell death model to an MOI model, we will evaluate which mechanism dominates in early infection and therefore explains the data better. But both may be equally important. To account for both cellular coinfection and density-dependence of infected cell death, in a relatively simple model, we use the MOI model as a basis and incorporate density-dependence with rate *γ*. We assume that the sheer number of infected cells enhances the death rate. That number is reflected in the term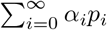 in the target cell equation.Similarly, internalized virus P is removed at a density-dependent way from the term 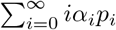. These terms become 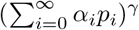 and 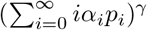 respectively.

Following the same simplifying assumptions as in the *MOI model*, the system of differential equations becomes the following:

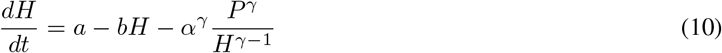

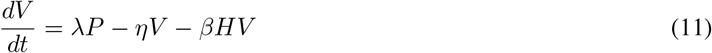

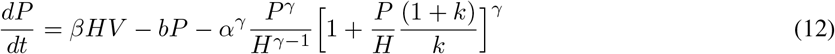

We restrict ourselves to these four simple models. We avoid models incorporating explicit adaptive immune responses due to additional parameters, which would render model fitting to our sparse data remarkably challenging. We do expect, though, adaptive immunity to play a significant role in the later phase of acute infection dynamics, namely the viral setpoint [7, 9, 18].

### 2.3 Model Fitting

We aim to capture the differences in viral load dynamics among study participants, we fit each model to the viral load data of each study participant separately. We report the mean, median and standard deviation of estimated parameter values of each model (Table 1), as well as the estimated values for each study participant (Supplementary Table 2-5). We fix the background target cell death rate *d* = 0.01 per day [9]. We assume that prior to infection, target cells are at equilibrium and therefore, we set *λ* = *d· T*_0_, and fix the equilibrium target cell count *T*_0_ = 10^6^ cells/mL, assuming that 1 in 1000 cells are available as targets for infection [9, 32]. We fit the mass action infectivity (*β*), the infected cell death rate (*δ*), the viral production rate (*p*), the viral clearance rate (*c*), the dispersion parameter of the binomial distribution (*k*) for the MOI models, the intensity of density-dependence (*γ*), as well as *t*_0_, which is the start of exponential viral growth. In order to estimate parameters, we use the *optim* package in *R* by performing general-purpose optimization [33].

**Table 1:** Parameter values: mean values and standard deviation for parameters of each model.

|  | **log<sub>10</sub> viral production (virions/cell day)** | **infected cell death rate (day<sup>-1</sup>)** | **viral clearance rate (day<sup>-1</sup>)** | **mass action infectivity (mL/ virion day)** | **t<sub>0</sub> (days)** | **γ** | **Dispersion parameter** |
| --- | --- | --- | --- | --- | --- | --- | --- |
| Standard | 2.07(0.71) | 0.81(1.22) | 9.98(6.94) | $6.77 \times 10^{-7} (1.72 \times 10^{-6})$ | -8.41(6.77) | | |
| Density-Dependent Cell Death | 3.04(0.73) | 2.08(1.39) | 13.1(9.37) | $1.42 \times 10^{-7} (3.5 \times 10^{-7})$ | -6.13(5.12) | 1.04(1.03) | |
| MOI | 2.29(0.81) | 1.04(0.99) | 11(10.02) | $6.8 \times 10^{-7} (1.9 \times 10^{-6})$ | -11.3(8.1) | | 3.01(3.89) |
| Density-Dependent Cell Death & MOI | 2.7(1.14) | 2.3(3.13) | 10.74(3) | $1.69 \times 10^{-6} (6.8 \times 10^{-6})$ | -9.03(4.61) | 1.02(1.02) | 9.75(16.9) |

### 2.4 Model Comparison

To determine which model explains the data set best for each participant, we use the Akaike Information Criterion (AIC), the Bayesian Information Criterion (BIC) and the corrected Akaike Information Criterion (AICc). BIC penalizes more heavily for more parameters [34], and AICc corrects for small sample sizes [35].

In addition to assessing the overall performance of the model, we are interested in evaluating their ability to predict certain quantitative measures, namely: 1) viral growth rate, 2) viral peak magnitude, 3) viral peak time, 4) a joint peak measurement that combines both magnitude and time, 5) viral decay rate and 6) setpoint. Growth and decay rates are calculated as the tangents of the growth part and decay part of the fitted viral load curve, as described above (Figure 2B). Setpoint is the steady-state viremia. Peak magnitude is calculated as the maximum viral load and peak timing is the time when the maximum viral load occurs. The joint measurement for the peak is sum of the difference between the magnitude and the timing weighted by the respective means, as described above, 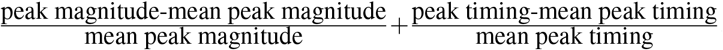. To compare the quantitative measures among models, we fit a linear regressions to the data-derived and the model-derived quantitative measures. The model that explains a quantitative measure best is the one for which the regression coefficient is closest to 1 and the intercept is closest to 0.

## 3 Results

### The Density-Dependent Cell Death model outperforms by AIC, BIC

We report the fitted parameter values for each study participant and each model (Supplementary Tables S2-S5) and the fitted curves to the viral load measurements of each study participants (Figure 3). We observe that all models perform comparably well. Table 1 contains the mean values of the estimated parameters for each model. The mean values of the infected cell death rate and viral clearance rate for all models appears to be higher compared to reported values for acute infection [9]. Mean values for viral production rate and mass action infectivity are comparable to reported values [9, 36]. Finally burst sizes, *p/δ*, are higher for the Density-Dependent Cell Death model model (Figure 4), more consistent with the values provided for Simian Immunodeficiency Virus (SIV) [15]. We also note that the Density-Dependent Cell Death & MOI model is more difficult to fit.

**Figure 3:**
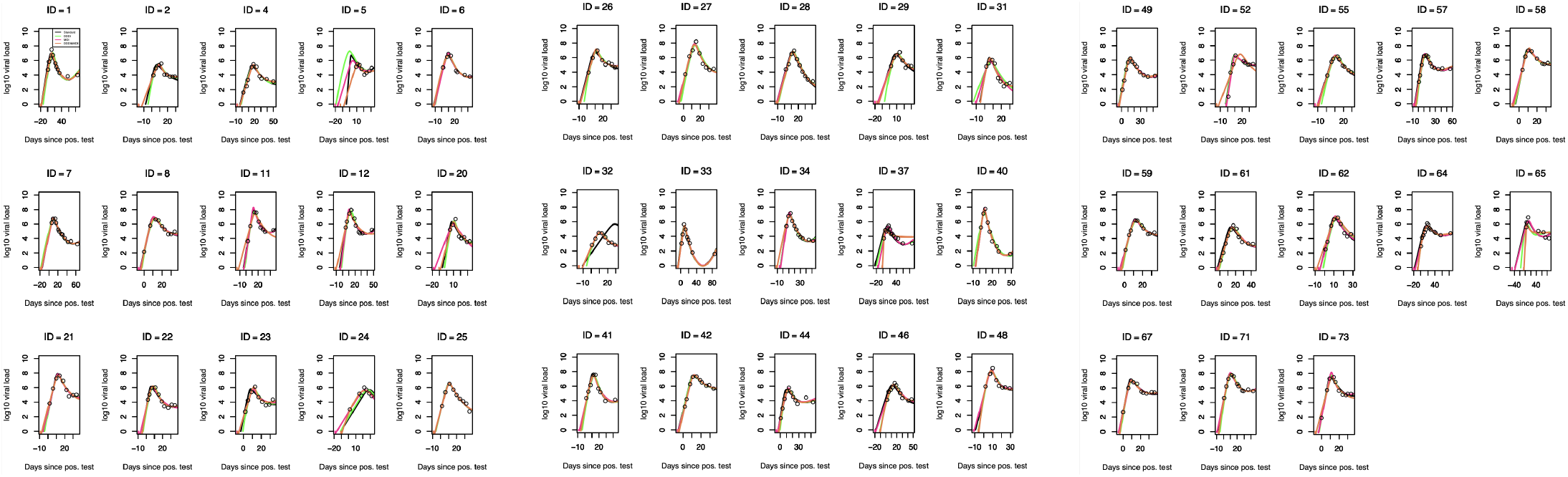
Viral load measurement for each study participant (black points) and fitted curves for the Standard model (black line), Density depedent cell death model (green line), MOI model (pink line) and Density depedent cell death model & MOI model (orange line). Curves overlap, so they are not all visible.

**Figure 4:**
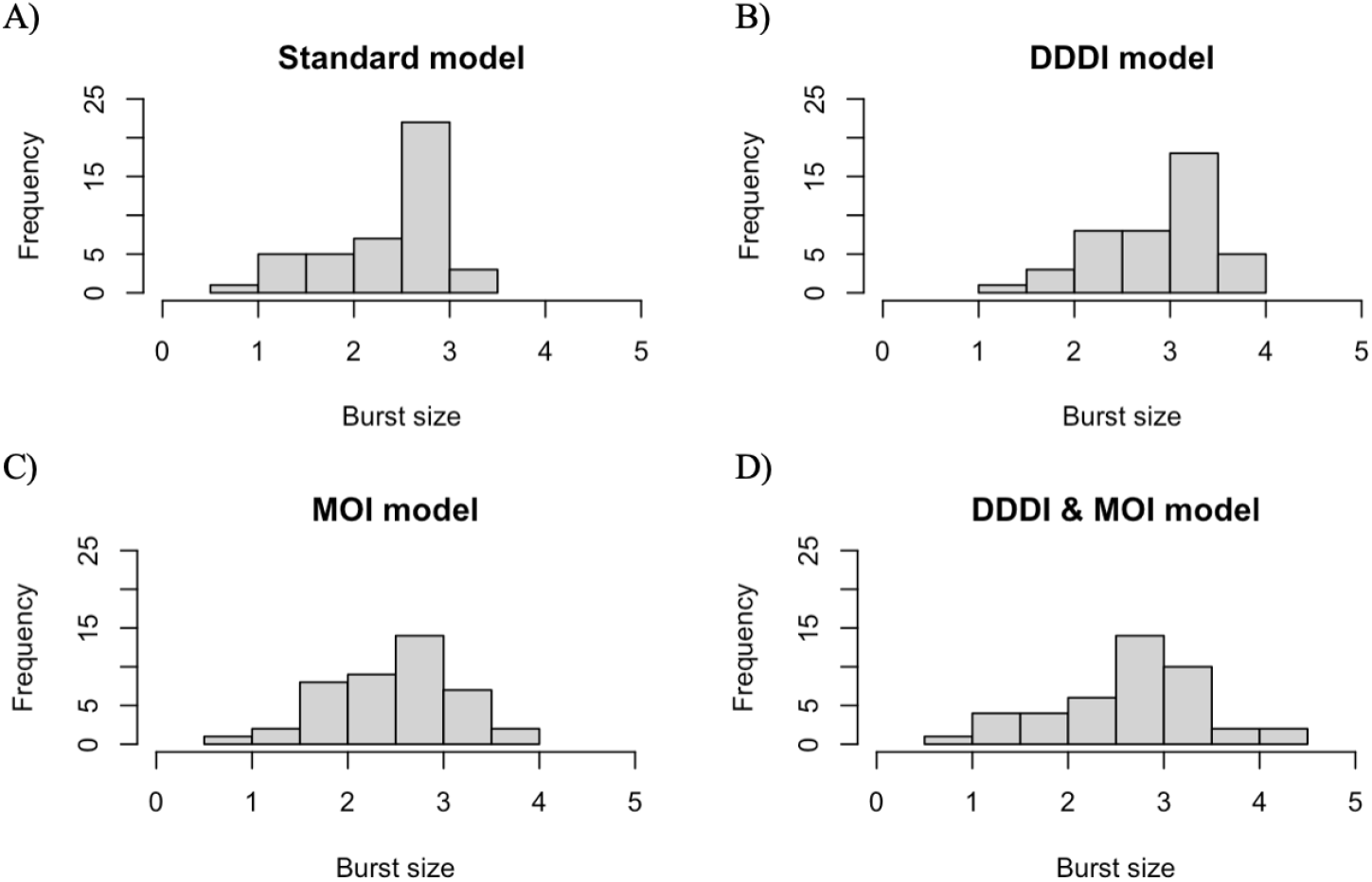
Distribution of the estimated viral burst sizes for the Standard model (A), Density-dependent cell death model (B), MOI model (C) and Density-dependent cell death & MOI model (D).

We first compare the performance of each model by computing the Akaike Information Criterion (AIC), the Bayesian Infromation Criterion (BIC) and the corrected Akaike Information Criterion (AICc). Table 2 reports the number of study participants for which a particular model is selected by the each estimator. The Density-Dependent Cell Death model is selected by AIC and BIC for the majority of study participants (24 out of 43 study participants). AICc selects overwhelmingly for the Standard model (39 out of 43), suggesting that the standard model performs better for a small number of data points. Using AIC, BIC and AICc, we are interested in determining if other models perform comparably well to the best model for each study participant. In Table 2 we also report for how many study participants a model fit yield the lowest or comparable to the lowest AIC, BIC and AICc (i.e. a difference of *<* 4 [37]). We find that the Density-Dependent Cell Death model still outperforms by AIC and BIC, whereas the Standard model outperforms by AICc. However, for a considerable number of study participants, the standard model provides comparable fits to the density-dependent model (using AIC - 29 participants, or BIC - 32 participants). The MOI and Density-Dependent & MOI models consistently underperform by AIC, BIC and AICc.

**Table 2:** AIC, BIC and AICc. Each row of the table represents for how many study participants the AIC/BIC/AICc is the lowest or comparable to the lowest.

|  | **AIC** | **AIC not different** | **BIC** | **BIC not different** | **AICc** | **AICc not different** |
| --- | --- | --- | --- | --- | --- | --- |
| Standard | 17 | 29 | 17 | 32 | 40 | 40 |
| Density-Dependent Cell Death | 24 | 36 | 24 | 35 | 1 | 5 |
| MOI | 0 | 4 | 0 | 4 | 0 | 0 |
| Density-Dependent Cell Death & MOI | 2 | 7 | 2 | 6 | 2 | 2 |

The AIC, BIC and AICc provide an estimate for the goodness of fit for each model. We are interested in determining how well these models can predict specific viral quantitative measurements namely, the growth rate, the peak, the peak timing, the decay rate and the setpoint. To determine how well each model explains a quantitative measurement, we fit a linear regression between the data-derived and the model-derived values of that quantitative measurement. The model that describe the quantitative measurement the best is the one for which the regression coefficient is closest to 1, the intercept is closest to 0 and has the highest *R*^2^. Tables 3-8 contain the regression fits for each model. Figure 5 show the data-derived and model derived *quantitative measurements* and the respective regression lines.

**Table 3:** Growth rate comparison: we report the coefficient, intercept (related p-values in parentheses) and *R*^2^ of the linear regression fitted to the data-derived and model-derived growth rate for each model tested.

|  | **Coefficient** | **Intercept** | **Adjusted $R^2$** |
| --- | --- | --- | --- |
| Standard | 0.29 ($3.57 \times 10^{-8}$) | 0.2 (0.0003) | 0.14 |
| Density-Dependent Cell Death | 0.71 (0.014) | 0.03 (0.59) | 0.47 |
| MOI | 0.69 (0.006) | 0.06 (0.18) | 0.49 |
| Density-Dependent Cell Death & MOI | 0.27 ($5 \times 10^{-6}$) | 0.22 (0.0008) | 0.06 |

**Figure 5:**
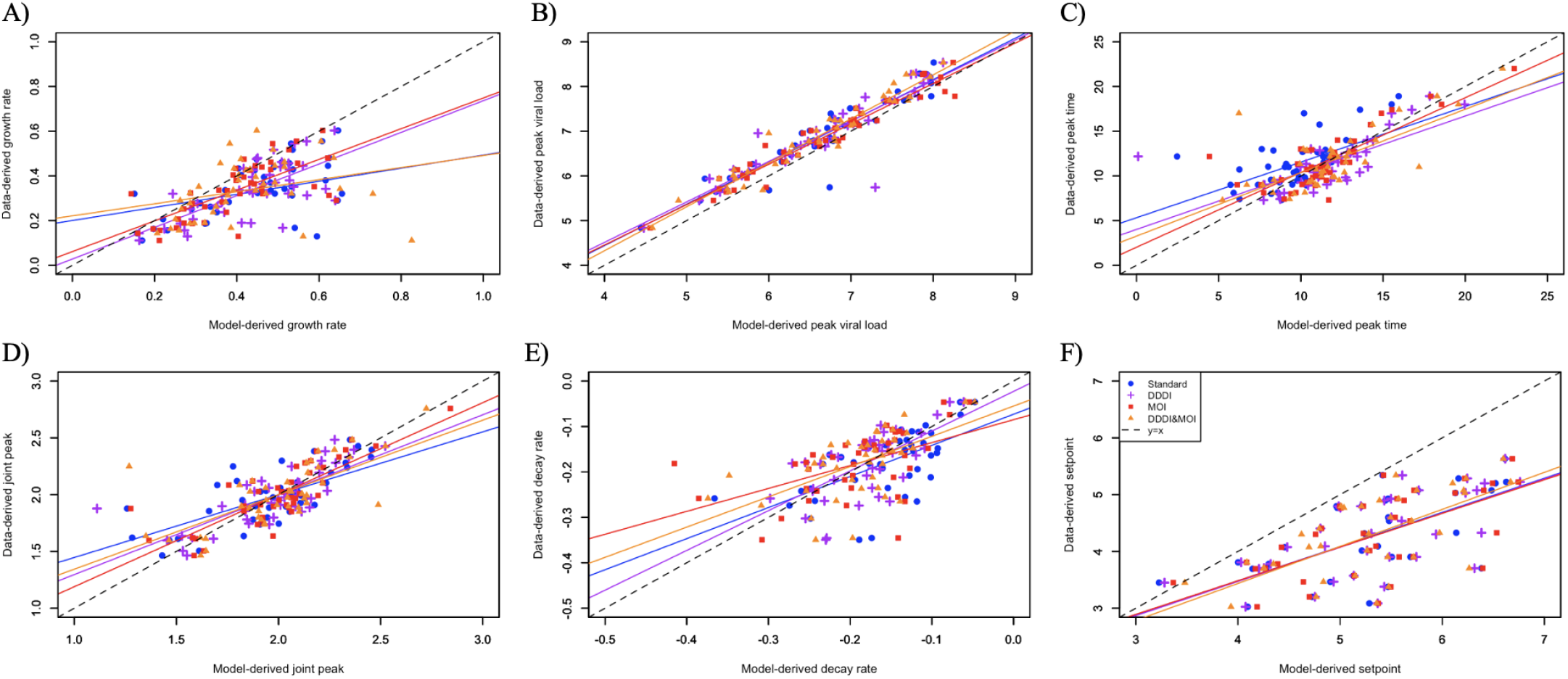
Data-derived and model derived measurements (solid points) for each study participant and model, along with the derived regression line (solid line) and the y=x line (dashed line) for the growth rate (A), peak magnitude (B), decay rate (C), peak time (D), joint peak measurement(E) and setpoint (F).

### All models overestimate the growth rate

We find that the growth rate is best explained by the Density-Dependent Infected Cell Death model. Notably, the Standard model and the Density-Dependent Infected Cell Death & MOI model explain the growth rate particularly poorly (Table 3). All models tend to predict fast growth, much faster than the data suggests. We hypothesize that this may attributable to a central assumption of all models: mass action infectivity of target cells. The models we explore do not incorporate the effects of spatial structure in the viral population that may be important especially in early dynamics before the infection becomes systemic. For HIV there is evidence for a spatial structure coming from genetic compartmentalization of the virus [31, 38, 39].

### Models incorporating cellular coinfection explain best all measurements of the peak

All models are successful in explaining peak magnitude (Table 4); the best model is the Density-Dependent Cell Death & MOI one. Regarding the timing of the peak, we find that the MOI model explains it better (Table 5). Similarly, our joint peak measurement, which incorporates both the magnitude and the timing of the peak, is better explained by the MOI model (Table 6). These results suggest that burst-size heterogeneity is important to explain the viral peak. We hypothesize that this is due to the assumption that cellular coinfection increases viral output compared to cell infection with a single virion. The ability to infect a cell multiple times reduces target cell limitation, allowing for higher peak viremia.

**Table 4:** Peak magnitude comparison: we report the coefficient, intercept (related p-values in parentheses) and *R*^2^ of the linear regression fitted to the data-derived and model-derived peak magnitude for each model tested.

|  | **Coefficient** | **Intercept** | **Adjusted $R^2$** |
| --- | --- | --- | --- |
| Standard | 0.93 (0.15) | 0.73 (0.03) | 0.89 |
| Density-Dependent Cell Death | 0.91 (0.12) | 0.9 (0.03) | 0.85 |
| MOI | 0.9 (0.01) | 0.84 (0.0013) | 0.94 |
| Density-Dependent Cell Death & MOI | 0.98 (0.69) | 0.39 (0.16) | 0.93 |

**Table 5:**
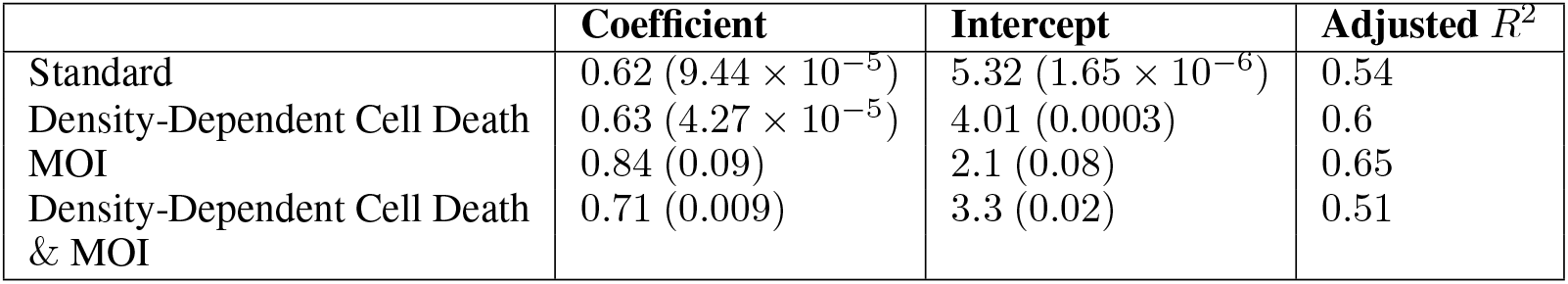
Peak timing comparison: we report the coefficient, intercept (related p-values in parentheses) and *R*^2^ of the linear regression fitted to the data-derived and model-derived peak timing for each model tested.

|  | **Coefficient** | **Intercept** | **Adjusted $R^2$** |
| --- | --- | --- | --- |
| Standard | 0.62 ($9.44 \times 10^{-5}$) | 5.32 ($1.65 \times 10^{-6}$) | 0.54 |
| Density-Dependent Cell Death | 0.63 ($4.27 \times 10^{-5}$) | 4.01 (0.0003) | 0.6 |
| MOI | 0.84 (0.09) | 2.1 (0.08) | 0.65 |
| Density-Dependent Cell Death & MOI | 0.71 (0.009) | 3.3 (0.02) | 0.51 |

**Table 6:**
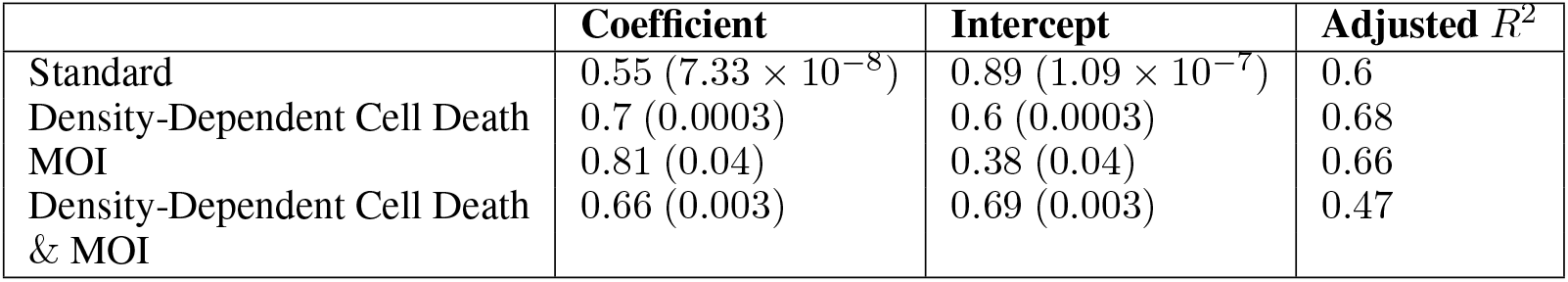
Joint peak measurement comparison: we report the coefficient, intercept (related p-values in parentheses) and *R*^2^ of the linear regression fitted to the data-derived and model-derived joint peak measurement for each model tested.

|  | **Coefficient** | **Intercept** | **Adjusted $R^2$** |
| --- | --- | --- | --- |
| Standard | 0.55 ($7.33 \times 10^{-8}$) | 0.89 ($1.09 \times 10^{-7}$) | 0.6 |
| Density-Dependent Cell Death | 0.7 (0.0003) | 0.6 (0.0003) | 0.68 |
| MOI | 0.81 (0.04) | 0.38 (0.04) | 0.66 |
| Density-Dependent Cell Death & MOI | 0.66 (0.003) | 0.69 (0.003) | 0.47 |

### Immune responses explain viral decay

the decay rate is better explained by the Density-Dependent Cell Death model (Table 7), suggesting that immune responses are increasingly important in explaining viral dynamics. It should be noted that the Standard model is nested within the Density-Dependent Cell Death model. The presence of the *γ* parameter in the Density-Dependent Cell Death model, which is a proxy the adaptive immune response, allows for more flexibility to capture both fast and slow decays.

**Table 7:**
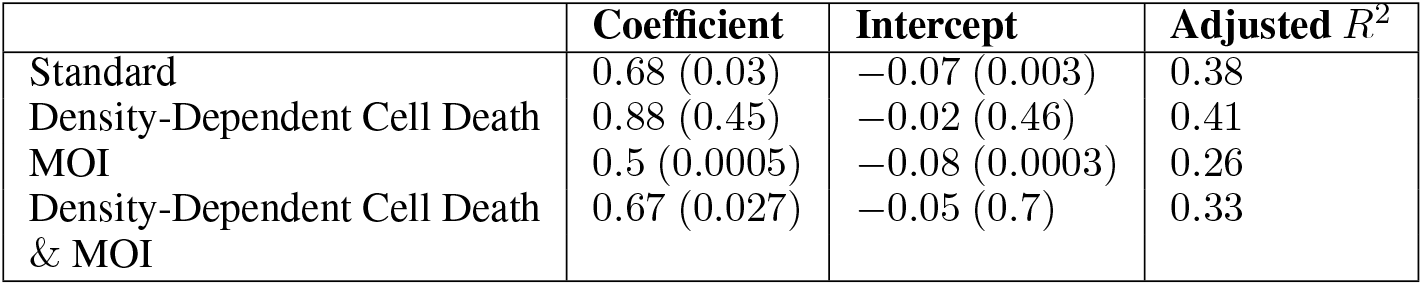
Decay rate comparison: we report the coefficient, intercept (related p-values in parentheses) and *R*^2^ of the linear regression fitted to the data-derived and model-derived decay rate for each model tested.

### All models overestimate the setpoint

Even though the setpoint is better explained by the Density-Dependent Cell Death & MOI model (Table 8), it should be noted that none of the models is particularly successful in explaining it. All models overestimate viral setpoint, which suggests that a more robust adaptive immune response not accounted for in our modeling is present at that phase.

**Table 8:** Setpoint comparison: we report the coefficient, intercept (related p-values in parentheses) and *R*^2^ of the linear regression fitted to the data-derived and model-derived setpoint for each model tested.

|  | **Coefficient** | **Intercept** | **Adjusted $R^2$** |
| --- | --- | --- | --- |
| Standard | 0.6 (0.0023) | 1.07 (0.11) | 0.4 |
| Density-Dependent Cell Death | 0.6 (0.001) | 1.6 (0.09) | 0.43 |
| MOI | 0.59 (0.001) | 1.11 (0.08) | 0.41 |
| Density-Dependent Cell Death & MOI | 0.65 (0.008) | 0.83 (0.21) | 0.43 |

## 4 Discussion

The Density-dependent cell death model provides the best goodness of fit by AIC and BIC, which is consistent with the results of Reeves et al. [20]. Reeves et al. fitted several compartment models to the the RV217 dataset in a nonlinear mixed-effects framework and determined that the model explaining the data best is the Density-dependent cell death one [20]. For their fitting, they fix the viral clearance rate to 23 *day*^−1^. We estimated a mean value of 13.1 *day*^−1^, which is lower compared to their fixed value. We also find mass action infectivity to be approximately three orders of magnitude lower compared to their estimates. In constrast, we find the infected cell death rate to be an order of magnitude higher compared to Reeves et al’s estimates. Surprisingly, the simple target cell-limited model also provides reasonable fits, and with fewer parameters is often the best model by AICc.

Peak viremia, both its timing and magnitude, is best explained by models that incorporate cellular coinfection. This is analogous to Koelle et al. [22]. In their study, they validated their MOI model using within-host data from ponies experimentally infected with influenza virus A subtype H3N8. In addition to finding by model selection that the MOI model performs better than the standard model, it also captureas the peak more effectively than the standard model, which tends to underestimate it, both for influenza and HIV. It appears allowing for multiple cellular infections in the modeling framework and permitting increased viral production rates for multiply infected cells, increases plasma viremia closer to observed peak levels. Regarding the magnitude of the peak, we find that the combination of MOI and Density-Dependent Cell Death is most successful at capturing it. This finding suggests that immune responses start to appear and play a role in viral dynamics. The decay rate is also best explained by the Density-Dependent Cell Death model, indicating that the effect of adaptive immune responses is exacerbated. Our results support that adaptive immune responses contribute to the initial decline in viral load, but do not exclude the effect of target cell limitation, as suggested by Philips (1996) [8, 9]. Indeed there have been observations that suggest that the immune systems controls viremia during primary infection: there is an negative association between an increase in CTL precursor frequency and a decline in viremia [7]. In addition, there has been documented an inverse correlation between the setpoint viral load and the number of effector CTLs [9, 40].

We found that the model that describes the setpoint best is the Density-dependent cell death model & MOI, indicating that both cellular coinfection and immune responses may be important in explaining viral dynamics. Nevertheless, no model, including the Density-dependent cell death model & MOI model one is particularly successful in explaining the setpoint, as is demonstrated by both the regression coefficient and the adjusted *R*^2^. This suggests that the these models are not enough to explain setpoint. Adaptive immune responses are already present and control the virus. Instead a model that explicitly incorporates adaptive immune responses is expected to explain setpoint dynamics more effectively.

Cellular coinfection in people with HIV has been experimentally demonstrated [21, 28–30]. The coinfection of a cell with multiple virions can lead to the formation of new recombinant viruses [21], which in turn can affect host adaptation [31], as well as the emergence and spread of drug resistance [41–44]. The MOI model developed by Koelle et al. [22] is not the first attempt to model cellular coinfection. A class of ODE models was developed by Wodarz & Levy [31], Guo et al. [45] and Dixit & Perelson [17, 23]. These models are high-dimensional and, to our knowledge, have mainly been used to derive qualitative conclusions about viral dynamics. In the MOI model, we make certain assumptions, namely that both the viral production rate, *λ*_*i*_, and the infected cell death rate, *α*_*i*_, scale linearly with MOI, *i* (*λ*_*i*_ = *λ· i, α*_*i*_ = *α· i*. This assumption is consistent with Koelle et al. (2009) [22]. Regardless, there are different burst size distributions that could be explored by the model. Indeed, an increased burst size can alter basic infection dynamics: under this scenario, virus population does not follow straight exponential growth, but the rate of exponential growth can increase over time, as viremia increases [24], with effects on host adaptation. Despite these theoretical predictions, there are no empirical experiments on HIV that quantify these relationships.

AIC, BIC and AICc are valuable in evaluating the overall goodness of fit and to compare models of different number of parameters. However, as our results suggest, these estimators do not provide a complete picture of a model’s performance. Even though a particular model best explains overall dynamics, different models explain different specific aspects of the dynamics. For example, we find that overall, HIV-1 acute infection dynamics are best described by a Density-Dependent Infected Cell Death model [18], yet peak viremia is best explained by the MOI model [22]. This suggests that if we are interested in the viral peak, instead of the Density Dependent Infected Cell Death, we should use the MOI model. In spite of these, we find that all models provide insight into the viral kinetics and within-host dynamics of acute HIV infection.

## 5 Acknowledgements

J.M.C. acknowledges the support of the National Science Foundation (grant no. DMS-1714654) and National Institutes of Health (grant nos. R21-AI143443-01A1 and R01-OD011095). E.M. acknowledges the support of the National Science Foundation (grant no. DMS-1714654). R.M.R. acknowledges NIH grants R01-AI152703 and R01-AI028433.

The Data used in this study derived from a study funded with Federal funds from Cooperative Agreement W81XWH-18-2-0040 issued by The U.S. Army Medical Research Acquisition Activity.

We thank the RV 217 study volunteers and study teams in Uganda, Kenya and Thailand for their contribution.

## 6 Disclaimer

The content is solely the responsibility of the authors and does not necessarily represent the position or policy of the US Army, the US Department of Defense, the US Department of Health and Human Services, or the US government. No official endorsement should be inferred.

## Notes

### Competing Interest Statement

The authors have declared no competing interest.

