## Supplementary Information for "Investigating Alternative Models of Acute HIV Infection"

**Alternative models of Acute HIV Infection—Supplementary Material**

Table S1: Growth, decay, peak and setpoint linear model parameter estimation and Standard-model derived $R_0$ for each study participant. We derive $R_0 = (1+r/d)e^{r\tau}$, where r=growth rate, d=decay rate and $\tau$= 1 day is the eclipse phase. $R_0$ is reported for the reader's comparison to [1]. We also report the least squares estimate, which is the objective function used in *optim* for the goodness of fit for each linear model. "NA" signifies that the measurement was not estimated.

| ID | Growth rate | Intercept growth model | Least squares for growth model | Decay rate | Intercept decay model | Least squares for decay model | Setpoint | Least squares for setpoint model | Peak magnitude | Peak timing | $R_0$ |
|---|---|---|---|---|---|---|---|---|---|---|---|
| 1 | 0.25 | 4.58 | 0.2016 | -0.13 | 8.56 | 0.0819 | 3.9 | 0.0153 | 7.51 | 11 | 3.79 |
| 2 | 0.32 | 3.31 | 0 | -0.14 | 6.72 | 0.5309 | 3.69 | 0.1233 | 5.71 | 7.41 | 4.65 |
| 3 | NA | NA | NA | -0.06 | 6.29 | 0.4739 | 3.6 | 0.2921 | NA | NA | NA |
| 4 | 0.28 | 0.7 | 0.1383 | -0.21 | 9.95 | 0.0507 | 3.81 | 0.4583 | 5.94 | 18.91 | 3.04 |
| 5 | 0.19 | 3.43 | 0 | -0.19 | 8.08 | 8E-04 | 4.3 | 1.0839 | 5.74 | 12.17 | 2.41 |
| 6 | 0.24 | 4.57 | 0.0022 | -0.24 | 9.93 | 0.0409 | 4.9 | 0.1729 | 7.27 | 11.34 | 2.55 |
| 7 | 0.19 | 5.05 | 0 | -0.11 | 7.78 | 0.5335 | 3.38 | 0.2315 | 6.78 | 9.28 | 3.31 |
| 8 | 0.52 | 2.16 | 0 | -0.15 | 8.74 | 0.0529 | 3.9 | 0.8169 | 7.25 | 9.83 | 7.39 |
| 9 | NA | NA | NA | -0.1 | 6.5 | 0.0029 | 2.64 | 0.1699 | NA | NA | NA |
| 10 | NA | NA | NA | -0.23 | 7.82 | 0.1995 | 4.29 | 0.2713 | NA | NA | NA |
| 11 | 0.35 | 2.86 | 0.2604 | -0.18 | 10.34 | 0.4038 | NA | NA | 7.78 | 14.01 | 4.16 |
| 12 | 0.31 | 4.03 | 0.0151 | -0.24 | 11.13 | 0.2202 | 5.2 | 0.513 | 8.07 | 12.92 | 3.17 |
| 13 | NA | NA | NA | NA | NA | NA | NA | NA | NA | NA | NA |
| 14 | NA | NA | NA | -0.07 | 6.41 | 2.6683 | 4.55 | 0.0174 | NA | NA | NA |
| 15 | NA | NA | NA | -0.07 | 5.95 | 3.7063 | 2.28 | 0.4413 | NA | NA | NA |
| 16 | NA | NA | NA | -0.33 | 10.9 | 0.0648 | 3.33 | 2.5103 | NA | NA | NA |
| 17 | NA | NA | NA | -0.08 | 8.44 | 0.1341 | 3.53 | 0.1992 | NA | NA | NA |
| 18 | NA | NA | NA | -0.14 | 6.78 | 0.7755 | 2.51 | 0.0469 | NA | NA | NA |
| 19 | 0.62 | 2.71 | 0 | -0.3 | 11.45 | 0.0033 | NA | NA | 8.61 | 9.55 | 5.7 |
| 20 | 0.17 | 4.38 | 0 | -0.35 | 10.99 | 0.3772 | 3.46 | 0.4849 | 6.68 | 13 | 1.76 |
| 21 | 0.43 | 3.07 | 0.0748 | -0.26 | 11.46 | 0.0652 | 5.08 | 0.8903 | 8.27 | 12.12 | 4.03 |
| 22 | 0.32 | 3.72 | 0.0021 | -0.18 | 7.8 | 0.3531 | 3.2 | 2.8902 | 6.32 | 8.1 | 3.79 |
| 23 | 0.38 | 2.07 | 0.1184 | -0.21 | 8.56 | 0.2414 | 4.08 | 1.9058 | 6.26 | 11.03 | 4.13 |
| 24 | 0.14 | 2.71 | 0.2823 | -0.12 | 8.7 | 6E-04 | NA | NA | 6.01 | 22 | 2.53 |
| 25 | 0.48 | 2.5 | 0 | -0.15 | 8.04 | 0.0671 | 3.57 | 0.3811 | 6.74 | 8.93 | 6.87 |
| 26 | 0.4 | 3.14 | 0 | -0.19 | 9.43 | 0.0144 | 4.54 | 0.0089 | 7.39 | 10.52 | 4.62 |
| 27 | 0.37 | 3.15 | 0.2873 | -0.3 | 12.49 | 0.3569 | 3.7 | 0.1305 | 8.29 | 13.89 | 3.22 |
| 28 | 0.26 | 3.75 | 0.0864 | -0.22 | 9.82 | 0.0783 | 3.09 | 0.6549 | 7.07 | 12.63 | 2.87 |
| 29 | 0.19 | 4.59 | 0.0937 | -0.27 | 10.42 | 0.0033 | 4.32 | 0.1053 | 7.01 | 12.85 | 2.07 |

| | | | | | | | | | | | |
|---|---|---|---|---|---|---|---|---|---|---|---|
| 30 | NA | NA | NA | NA | NA | NA | 4.23 | 0.5085 | NA | NA | NA |
| 31 | 0.13 | 4.53 | 0.0011 | -0.27 | 8.08 | 0.4891 | 2.69 | 0.3771 | 5.68 | 9 | 1.68 |
| 32 | 0.21 | 2.63 | 8E-04 | -0.16 | 6.59 | 0.0104 | 3.45 | 0.0072 | 4.84 | 10.72 | 2.78 |
| 33 | 0.43 | 2.15 | 0.0043 | -0.17 | 6.8 | 0.0967 | NA | NA | 5.64 | 8 | 5.53 |
| 34 | 0.46 | 2.67 | 0 | -0.16 | 8.86 | 0.5014 | NA | NA | 7.24 | 10 | 6.02 |
| 35 | NA | NA | NA | NA | NA | NA | 2.49 | 1.3984 | NA | NA | NA |
| 36 | NA | NA | NA | -0.12 | 7.64 | 1.9544 | NA | NA | NA | NA | NA |
| 37 | 0.11 | 3.65 | 0.2605 | -0.05 | 5.64 | 0.1822 | 3.02 | 0.0185 | 5.45 | 17 | 3.81 |
| 38 | NA | NA | NA | NA | NA | NA | NA | NA | NA | NA | NA |
| 39 | NA | NA | NA | NA | NA | NA | NA | NA | NA | NA | NA |
| 40 | 0.48 | 4.27 | 0 | -0.26 | 9.32 | 1.7897 | NA | NA | 7.76 | 9 | 4.62 |
| 41 | 0.32 | 3.66 | 0.014 | -0.24 | 10.73 | 0.5472 | NA | NA | 7.69 | 12.55 | 3.2 |
| 42 | 0.46 | 2.34 | 0.0903 | -0.11 | 8.99 | 0.0093 | 5.29 | 2.0203 | 7.67 | 11.54 | 8 |
| 43 | NA | NA | NA | -0.1 | 6.55 | 0.0016 | 4.96 | 0.2073 | NA | NA | NA |
| 44 | 0.33 | 1.71 | 0.0654 | -0.16 | 7.96 | 0.0183 | 3.78 | 6E-04 | 5.95 | 12.68 | 4.35 |
| 45 | NA | NA | NA | -0.05 | 5.4 | 0.0172 | 4.87 | 0.0081 | NA | NA | NA |
| 46 | 0.16 | 4.07 | 0.2138 | -0.15 | 8.86 | 0.0207 | 4.77 | 0.2143 | 6.53 | 15.74 | 2.41 |
| 47 | NA | NA | NA | NA | NA | NA | NA | NA | NA | NA | NA |
| 48 | 0.34 | 5.27 | 0 | -0.35 | 11.88 | 0.0509 | 5.22 | 0.0695 | 8.54 | 9.56 | 2.79 |
| 49 | 0.43 | 1.56 | 0.0427 | -0.11 | 7.74 | 0.0528 | 4.8 | 0.5065 | 6.46 | 11.53 | 7.41 |
| 50 | NA | NA | NA | -0.05 | 6.52 | 0.0192 | 4.37 | 0.015 | NA | NA | NA |
| 51 | 0.03 | 1.44 | 0.4048 | NA | NA | NA | 4.66 | 1.2892 | NA | NA | NA |
| 52 | 0.29 | 3.14 | 0 | -0.07 | 7.48 | 0.0014 | 5.34 | 0.0871 | 6.66 | 11 | 6.57 |
| 53 | NA | NA | NA | NA | NA | NA | 5.08 | 0.1126 | NA | NA | NA |
| 54 | NA | NA | NA | -0.06 | 6.26 | 0.0018 | 4.57 | 0.9262 | NA | NA | NA |
| 55 | 0.34 | 3.61 | 0.0463 | -0.13 | 8.05 | 0.1456 | 4.01 | 0.004 | 6.79 | 9.47 | 4.96 |
| 56 | NA | NA | NA | NA | NA | NA | NA | NA | NA | NA | NA |
| 57 | 0.39 | 2.75 | 0.0982 | -0.1 | 7.87 | 0.4488 | 4.6 | 0.7161 | 6.86 | 10.39 | 7.5 |
| 58 | 0.45 | 3.06 | 0.1568 | -0.14 | 9.15 | 0.0228 | 5.03 | 0.3094 | 7.69 | 10.39 | 6.49 |
| 59 | 0.36 | 2.35 | 0.2753 | -0.15 | 8.64 | 0.009 | 4.8 | 0.3294 | 6.77 | 12.16 | 4.84 |
| 60 | NA | NA | NA | -0.25 | 5.76 | 0.2266 | 2.78 | 0.927 | NA | NA | NA |
| 61 | 0.28 | 1.21 | 0.2635 | -0.26 | 10.58 | 0.0811 | 3.71 | 0.1297 | 6.14 | 17.4 | 2.8 |
| 62 | 0.54 | 1.95 | 9E-04 | -0.18 | 8.83 | 0.6186 | 4.93 | 0.1899 | 7.13 | 9.51 | 6.97 |
| 63 | NA | NA | NA | -0.19 | 6.42 | 0.0267 | 4.69 | 0.4413 | NA | NA | NA |
| 64 | 0.16 | 3.12 | 0.2906 | -0.05 | 6.67 | 0.0804 | 4.41 | 0.1355 | 6.09 | 18 | 5.35 |
| 65 | 0.32 | 4.62 | 0 | -0.04 | 7.25 | 0.1744 | 4.09 | 0.3025 | 6.95 | 7.3 | 12.16 |
| 66 | 0.05 | 5.3 | 2E-04 | -0.01 | 5.85 | 3E-04 | NA | NA | 5.84 | 11 | 9.72 |
| 67 | 0.55 | 2.65 | 0 | -0.11 | 8.07 | 0.1795 | 5.08 | 0.3339 | 7.16 | 8.13 | 10.36 |
| 68 | NA | NA | NA | -0.19 | 8.16 | 0.1075 | 4.36 | 0.0778 | NA | NA | NA |
| 69 | 0.29 | 2.94 | 0.1319 | -0.12 | 8.64 | 0.1516 | 5.07 | 0.2263 | 6.98 | 13.78 | 4.61 |
| 70 | NA | NA | NA | -0.02 | 5.02 | 0.0026 | 4.14 | 0.2032 | NA | NA | NA |
| 71 | 0.44 | 4.24 | 0.0561 | -0.14 | 9.48 | 0.1257 | 5.63 | 0.1775 | 8.21 | 9.03 | 6.4 |

| | | | | | | | | | | | |
|---|---|---|---|---|---|---|---|---|---|---|---|
| 72 | NA | NA | NA | NA | NA | NA | 4.12 | 0.1576 | NA | NA | NA |
| 73 | 0.6 | 1.93 | 0.0321 | -0.18 | 9.68 | 0.509 | 4.33 | 0.6403 | 7.89 | 9.88 | 7.91 |
| 74 | NA | NA | NA | -0.02 | 5.62 | 0.0052 | NA | NA | NA | NA | NA |
| 75 | NA | NA | NA | NA | NA | NA | NA | NA | NA | NA | NA |
| 76 | NA | NA | NA | NA | NA | NA | NA | NA | NA | NA | NA |
| 77 | NA | NA | NA | NA | NA | NA | NA | NA | NA | NA | NA |

Table S2: Parameter value estimates for the Standard model, along with the number of data points used to fit the model, the negative log likelihood (nll), AIC, BIC and AICc.

| **ID** | **$Log_{10}(p)$** | **delta** | **c** | **k** | **$t_0$** | **n** | **error value** | **Negative log likelihood** | **AIC** | **BIC** | **AICc** |
|---|---|---|---|---|---|---|---|---|---|---|---|
| 1 | 2.99 | 0.58 | 19.08 | 2.54E-08 | -17.59 | 11 | 0.0788 | 27.4 | 64.8 | 66.79 | 76.8 |
| 2 | 1.08 | 0.33 | 20.04 | 2.4238E-06 | -7.79 | 9 | 0.0837 | 16.78 | 43.57 | 44.55 | 63.57 |
| 4 | 0.75 | 0.35 | 10.97 | 2.23E-06 | -3.5 | 10 | 0.097 | 7.66 | 25.33 | 26.84 | 40.33 |
| 5 | 2.08 | 0.46 | 13 | 3.85E-07 | -4.6 | 8 | 0.0936 | 7.51 | 25.02 | 25.42 | 55.02 |
| 6 | 1.45 | 0.57 | 1.33 | 1.05E-07 | -12.62 | 8 | 0.0643 | 19.43 | 48.86 | 49.26 | 78.86 |
| 7 | 2.49 | 0.53 | 12.05 | 5.27E-08 | -14.7 | 12 | 0.0268 | 25.85 | 61.69 | 64.12 | 71.69 |
| 8 | 2.42 | 0.39 | 16.32 | 1.084E-07 | -4.35 | 9 | 0.0474 | 10.19 | 30.38 | 31.36 | 50.38 |
| 11 | 2.57 | 1.03 | 0.84 | 1.27E-08 | -4.03 | 9 | 0.0279 | 14.04 | 38.07 | 39.06 | 58.07 |
| 12 | 2.61 | 0.86 | 1.14 | 1.22E-08 | -5.21 | 10 | 0.0786 | 18.74 | 47.47 | 48.98 | 62.47 |
| 20 | 1.44 | 0.51 | 5.14 | 4.08E-07 | -5.71 | 8 | 0.2151 | 12.79 | 35.58 | 35.97 | 65.58 |
| 21 | 2.53 | 0.82 | 1.09 | 1.42E-08 | -4.91 | 9 | 0.0839 | 17.29 | 44.59 | 45.57 | 64.59 |
| 22 | 1.59 | 0.48 | 11.89 | 6.87E-07 | -4.89 | 10 | 0.107 | 17.65 | 45.3 | 46.81 | 60.3 |
| 23 | 0.92 | 0.36 | 6.7 | 1.73E-06 | -2.97 | 9 | 0.1841 | 10.47 | 30.93 | 31.92 | 50.93 |
| 24 | 3.45 | 6.13 | 13.99 | 3.29E-08 | -10.6 | 7 | 0.0581 | 14.84 | 39.68 | 39.41 | 99.68 |
| 25 | 2.42 | 0.62 | 20.27 | 1.34E-07 | -6.11 | 9 | 0.0369 | 16.53 | 43.07 | 44.05 | 63.07 |
| 26 | 1.81 | 0.35 | 4.72 | 1.14E-07 | -8.88 | 8 | 0.0615 | 16.16 | 42.33 | 42.72 | 72.33 |
| 27 | 2.55 | 0.94 | 1.2 | 1.33E-08 | -5.15 | 9 | 0.1613 | 17.9 | 45.8 | 46.79 | 65.8 |
| 28 | 2.64 | 0.99 | 10.86 | 4.72E-08 | -11.13 | 10 | 0.0657 | 22.64 | 55.29 | 56.8 | 70.29 |
| 29 | 2.37 | 0.41 | 21.78 | 1.12E-07 | -14.05 | 8 | 0.1089 | 19.02 | 48.04 | 48.44 | 78.04 |
| 31 | 0.81 | 0.83 | 1.96 | 1.13E-06 | -5.87 | 9 | 0.2009 | 17.87 | 45.74 | 46.72 | 65.74 |
| 32 | 0 | 0.39 | 9.84 | 1.08E-05 | -12.66 | 8 | 0.023 | 14.95 | 39.91 | 40.31 | 69.91 |
| 33 | 1.2 | 0.91 | 12.28 | 1.52E-06 | -6.95 | 9 | 0.0385 | 16.67 | 43.33 | 44.32 | 63.33 |
| 34 | 2.29 | 0.73 | 2.94 | 4.18E-08 | -4.52 | 11 | 0.0184 | 17.06 | 44.11 | 46.1 | 56.11 |
| 37 | 1.15 | 0.32 | 19.58 | 1.02E-06 | -24.25 | 13 | 0.0465 | 27.52 | 65.05 | 67.87 | 73.62 |
| 40 | 2.73 | 1.49 | 2.03 | 1.894E-08 | -6.33 | 9 | 0.0551 | 20.55 | 51.1 | 52.08 | 71.1 |
| 41 | 2.35 | 1.13 | 1 | 2.13E-08 | -6.11 | 9 | 0.1432 | 19.1 | 48.2 | 49.19 | 68.2 |
| 42 | 2.78 | 0.29 | 11.9 | 3.19E-08 | -5.07 | 10 | 0.0076 | 17.2 | 44.4 | 45.92 | 59.4 |
| 44 | 0.71 | 0.27 | 7.95 | 2.43E-06 | -3.35 | 11 | 0.1189 | 11.13 | 32.27 | 34.26 | 44.27 |
| 46 | 2.09 | 0.37 | 21.39 | 1.63E-07 | -18.63 | 11 | 0.075 | 25.5 | 61.01 | 63 | 73.01 |

| | | | | | | | | | | | |
|---|---|---|---|---|---|---|---|---|---|---|---|
| 48 | 2.49 | 0.63 | 0.98 | 1.27E-08 | -9.32 | 8 | 0.1086 | 19.64 | 49.28 | 49.67 | 79.28 |
| 49 | 1.79 | 0.39 | 12.88 | 3.19E-07 | -4.27 | 12 | 0.0072 | 15.77 | 41.54 | 43.96 | 51.54 |
| 52 | 1.99 | 0.19 | 18 | 3.34E-07 | -2.66 | 9 | 0.1425 | 8.43 | 26.86 | 27.84 | 46.86 |
| 55 | 2.27 | 0.5 | 14.66 | 1.15E-07 | -10.05 | 9 | 0.0126 | 20.02 | 50.03 | 51.02 | 70.03 |
| 57 | 2.23 | 0.32 | 14 | 1.23E-07 | -6.91 | 12 | 0.0333 | 21.62 | 53.24 | 55.66 | 63.24 |
| 58 | 2.55 | 0.37 | 6.03 | 3.78E-08 | -4 | 8 | 0.019 | 13.03 | 36.07 | 36.47 | 66.07 |
| 59 | 1.72 | 0.36 | 7.12 | 2.22E-07 | -4.64 | 10 | 0.0343 | 16.2 | 42.4 | 43.91 | 57.4 |
| 61 | 0.68 | 0.43 | 4.29 | 1.40E-06 | -3.33 | 11 | 0.0767 | 7.86 | 25.72 | 27.71 | 37.72 |
| 62 | 2.38 | 0.6 | 9.55 | 8.22E-08 | -4.29 | 9 | 0.0855 | 15.65 | 41.3 | 42.28 | 61.3 |
| 64 | 1.43 | 0.16 | 19.09 | 4.56E-07 | -17.54 | 13 | 0.0429 | 27.27 | 64.54 | 67.36 | 73.11 |
| 65 | 2.45 | 0.31 | 21.41 | 5.06E-08 | -37.98 | 8 | 0.2838 | 21.06 | 52.12 | 52.52 | 82.12 |
| 67 | 2.16 | 0.33 | 6.46 | 9.32E-08 | -4.67 | 10 | 0.0429 | 10.67 | 31.34 | 32.85 | 46.34 |
| 71 | 2.71 | 0.44 | 2.3 | 1.05E-08 | -8.67 | 11 | 0.0313 | 25.06 | 60.12 | 62.11 | 72.12 |
| 73 | 2.61 | 0.48 | 5.02 | 3.12E-08 | -3.25 | 10 | 0.0753 | 11.72 | 33.43 | 34.95 | 48.43 |

Table S3: Parameter value estimates for the Density-Dependent Cell Death model, along with the negative log likelihood (nll), AIC, BIC and AICc.

| ID | $Log_{10}(p)$ | delta | c | k | γ | $t_0$ | error value | Negative log likelihood | AIC | BIC | AICc |
|---|---|---|---|---|---|---|---|---|---|---|---|
| 1 | 3.74 | 5 | 4.03 | 4.93E-09 | 0.01 | -12.63 | 0.0811 | 26.74 | 65.49 | 67.87 | 86.49 |
| 2 | 2.38 | 1.44 | 23.26 | 3.056E-07 | 0.05 | -4.64 | 0.0514 | 14.54 | 41.07 | 42.26 | 83.07 |
| 4 | 2.08 | 1.21 | 23 | 4.22E-07 | 0.03 | -1.42 | 0.0916 | 4.13 | 20.26 | 22.08 | 48.26 |
| 5 | 4.34 | 2.38 | 22.89 | 3.79E-09 | 0.02 | -19.06 | 0.1144 | 15.77 | 43.54 | 44.01 | 127.54 |
| 6 | 3.28 | 1.88 | 9.83 | 1.62E-08 | 0.02 | -9.82 | 0.0484 | 18.03 | 48.07 | 48.55 | 132.07 |
| 7 | 2.85 | 1.03 | 8.31 | 2.17E-08 | 0.01 | -18 | 0.0259 | 27.03 | 66.06 | 68.97 | 82.86 |
| 8 | 3.57 | 2.58 | 10.37 | 1.29E-08 | 0.04 | -2.39 | 0.0286 | 11.18 | 34.36 | 35.54 | 76.36 |
| 11 | 3.91 | 1.89 | 5.44 | 2.79E-09 | 0.04 | -1.79 | 0.0289 | 9.98 | 31.95 | 33.13 | 73.95 |
| 12 | 4.1 | 4.42 | 3.35 | 2.27E-09 | 0.03 | -1.84 | 0.0629 | 12.75 | 37.5 | 39.32 | 65.5 |
| 20 | 3.44 | 3.09 | 22.91 | 4.05E-08 | 0.02 | -2.77 | 0.1563 | 10.63 | 33.26 | 33.74 | 117.26 |
| 21 | 3.15 | 1.2 | 1.98 | 6.19E-09 | 0.05 | -3.97 | 0.0596 | 15.73 | 43.46 | 44.64 | 85.46 |
| 22 | 3.03 | 3.13 | 13.9 | 6.75E-08 | 0.03 | -4.01 | 0.0449 | 17.59 | 47.17 | 48.99 | 75.17 |
| 23 | 3.18 | 3.99 | 22.91 | 9.15E-08 | 0.03 | -0.46 | 0.1063 | 7.12 | 26.23 | 27.42 | 68.23 |
| 24 | 3.28 | 3.39 | 23.02 | 4.65E-08 | 0 | -11.23 | 0.0603 | 14.4 | 40.81 | 40.48 | Inf |
| 25 | 2.29 | 0.61 | 14.93 | 1.36E-07 | 0 | -5.97 | 0.0378 | 16.52 | 45.03 | 46.22 | 87.03 |
| 26 | 3.34 | 1.32 | 10.27 | 1.49E-08 | 0.05 | -3.62 | 0.0397 | 12.62 | 37.25 | 37.72 | 121.25 |
| 27 | 3.75 | 2.63 | 3.51 | 3.27E-09 | 0.02 | -2.99 | 0.1286 | 15.85 | 43.7 | 44.88 | 85.7 |
| 28 | 3.31 | 1.94 | 13.09 | 1.93E-08 | 0.01 | -8.42 | 0.0465 | 21.74 | 55.48 | 57.29 | 83.48 |
| 29 | 3.57 | 3.18 | 10.71 | 1.39E-08 | 0.03 | -6.75 | 0.0791 | 16.13 | 44.25 | 44.73 | 128.25 |
| 31 | 2.55 | 5 | 11.59 | 1.96E-07 | 0 | -13.45 | 0.2452 | 21.84 | 55.68 | 56.86 | 97.68 |
| 32 | 1.28 | 1.66 | 12.43 | 1.92E-06 | 0.04 | -5.57 | 0.0159 | 12.86 | 37.72 | 38.19 | 121.72 |

| | | | | | | | | | | | |
|---|---|---|---|---|---|---|---|---|---|---|---|
| 33 | 1.55 | 0.94 | 23.04 | 1.29E-06 | 0 | -6.09 | 0.0361 | 16.34 | 44.68 | 45.86 | 86.68 |
| 34 | 2.92 | 0.76 | 9.47 | 2.62E-08 | 0.03 | -4.16 | 0.0149 | 15.78 | 43.55 | 45.94 | 64.55 |
| 37 | 2.43 | 1.08 | 43.1 | 2.45E-07 | 0.01 | -19.89 | 0.0551 | 26.71 | 65.41 | 68.8 | 79.41 |
| 40 | 3.96 | 4.99 | 10.24 | 7.62E-09 | 0.01 | -9.51 | 0.1516 | 22.06 | 56.13 | 57.31 | 98.13 |
| 41 | 3.92 | 4.64 | 5.06 | 4.46E-09 | 0.02 | -4.04 | 0.1175 | 17.39 | 46.79 | 47.97 | 88.79 |
| 42 | 2.46 | 0.17 | 4.49 | 2.74E-08 | 0.06 | -4.79 | 0.0074 | 16.85 | 45.7 | 47.52 | 73.7 |
| 44 | 2.63 | 1.83 | 22.91 | 1.88E-07 | 0.04 | -0.17 | 0.0743 | 6.65 | 25.29 | 27.68 | 46.29 |
| 46 | 3.35 | 1.82 | 22.99 | 2.71E-08 | 0.02 | -10.45 | 0.0472 | 22.83 | 57.65 | 60.04 | 78.65 |
| 48 | 3.45 | 0.76 | 2 | 3.06E-09 | 0.09 | -6.08 | 0.0765 | 17.6 | 47.19 | 47.67 | 131.19 |
| 49 | 2.14 | 0.34 | 22.9 | 2.45E-07 | 0.03 | -3.97 | 0.0065 | 15.1 | 42.2 | 45.11 | 59 |
| 52 | 1.96 | 0.02 | 9.51 | 1.89E-07 | 0.2 | -2.34 | 0.1299 | 8.2 | 28.4 | 29.59 | 70.4 |
| 55 | 3.3 | 2.02 | 12.52 | 2.23E-08 | 0.03 | -5.55 | 0.0043 | 17.54 | 47.08 | 48.26 | 89.08 |
| 57 | 3.7 | 1.89 | 18.62 | 1.27E-08 | 0.04 | -2.93 | 0.0161 | 18.5 | 49.01 | 51.92 | 65.81 |
| 58 | 1.93 | 0.49 | 0.64 | 4.30E-08 | 0.04 | -4.58 | 0.0083 | 14.37 | 40.74 | 41.21 | 124.74 |
| 59 | 3.24 | 1.6 | 15.57 | 2.9E-08 | 0.05 | -2.1 | 0.0185 | 12.26 | 36.51 | 38.33 | 64.51 |
| 61 | 2.48 | 1.33 | 32.34 | 2.7E-07 | 0.03 | -0.15 | 0.0545 | 2.5 | 17.01 | 19.4 | 38.01 |
| 62 | 3.61 | 3.55 | 8.82 | 1.27E-08 | 0.03 | -1.27 | 0.0498 | 12.56 | 37.13 | 38.31 | 79.13 |
| 64 | 2.45 | 1.8 | 3.87 | 4.32E-08 | 0.04 | -7.8 | 0.0357 | 21.79 | 55.57 | 58.96 | 69.57 |
| 65 | 2.64 | 2.17 | 4.71 | 3.92E-08 | 0.04 | -16.57 | 0.4188 | 18.41 | 48.83 | 49.31 | 132.83 |
| 67 | 3.46 | 0.72 | 15.62 | 1.44E-08 | 0.09 | -2.92 | 0.0203 | 9.98 | 31.95 | 33.77 | 59.95 |
| 71 | 3.99 | 3.19 | 2.2 | 1.74E-09 | 0.05 | -5.02 | 0.0139 | 21.89 | 55.78 | 58.17 | 76.78 |
| 73 | 2.58 | 0.55 | 1.2 | 1.57E-08 | 0.07 | -2.78 | 0.0209 | 11.92 | 35.83 | 37.65 | 63.83 |

Table S4: Parameter value estimates for the MOI model, along the negative log likelihood (nll), AIC, BIC and AICc.

| ID | $Log_{10}(\lambda)$ | α | η | k | β | $t_0$ | error value | Negative log likelihood | AIC | BIC | AICc |
|---|---|---|---|---|---|---|---|---|---|---|---|
| 1 | 2.87 | 0.61 | 12.04 | 6.4 | 2.14E-08 | -20.73 | 0.0642 | 29.89 | 71.78 | 74.17 | 92.78 |
| 2 | 0.4 | 0.11 | 11.52 | 4.08 | 5.55E-06 | -16.53 | 0.0491 | 22.71 | 57.42 | 58.6 | 99.42 |
| 4 | 1.26 | 0.24 | 42.97 | 13.63 | 2.36E-06 | -8.46 | 0.0557 | 19.9 | 51.8 | 53.61 | 79.8 |
| 5 | 0.82 | 0.06 | 15.4 | 1.01 | 2.24E-06 | -20.47 | 0.1244 | 17.32 | 46.64 | 47.12 | 130.64 |
| 6 | 1.84 | 0.78 | 1.44 | 1.88 | 5.64E-08 | -11.67 | 0.0621 | 19.71 | 51.41 | 51.89 | 135.41 |
| 7 | 2.48 | 0.72 | 6.95 | 2.04 | 3.55E-08 | -17.99 | 0.0217 | 28.84 | 69.68 | 72.59 | 86.48 |
| 8 | 3.4 | 1.3 | 24.87 | 0.32 | 2.65E-08 | -6.04 | 0.0451 | 11.23 | 34.47 | 35.65 | 76.47 |
| 11 | 3.41 | 0.95 | 3.65 | 0.9 | 4.55E-09 | -3.83 | 0.0612 | 14.21 | 40.41 | 41.6 | 82.41 |
| 12 | 3.64 | 1.12 | 7.8 | 0.64 | 5.38E-09 | -4.41 | 0.1621 | 17.89 | 47.77 | 49.59 | 75.77 |
| 20 | 2.48 | 2.08 | 8.77 | 0.5 | 8.36E-08 | -18.6 | 0.2516 | 19.89 | 51.77 | 52.25 | 135.77 |
| 21 | 2.42 | 0.95 | 0.85 | 4.05 | 1.57E-08 | -6.35 | 0.0706 | 18.94 | 49.87 | 51.06 | 91.87 |
| 22 | 1.83 | 0.51 | 17.86 | 1.99 | 4.57E-07 | -9.37 | 0.0982 | 23.69 | 59.38 | 61.19 | 87.38 |
| 23 | 0.72 | 0.21 | 7.22 | 1.72 | 2.25E-06 | -7.44 | 0.1212 | 18.87 | 49.74 | 50.92 | 91.74 |

| | | | | | | | | | | | |
|---|---|---|---|---|---|---|---|---|---|---|---|
| 24 | 2.43 | 3.32 | 4.77 | 3.59 | 7.19E-08 | -18.58 | 0.059 | 15.83 | 43.67 | 43.34 | Inf |
| 25 | 2.73 | 1.5 | 11.19 | 0.58 | 5.623E-08 | -8.4 | 0.04 | 19.41 | 50.83 | 52.01 | 92.83 |
| 26 | 1.81 | 0.2 | 5.31 | 13.57 | 1.09E-07 | -10.3 | 0.0271 | 18.28 | 48.55 | 49.03 | 132.55 |
| 27 | 2.58 | 0.83 | 1.34 | 10.75 | 1.13E-08 | -7.21 | 0.131 | 19.89 | 51.77 | 52.96 | 93.77 |
| 28 | 2.2 | 1.3 | 2.55 | 2.67 | 4.35E-08 | -12.34 | 0.0491 | 24.1 | 60.2 | 62.02 | 88.2 |
| 29 | 2.16 | 0.39 | 13.44 | 2.53 | 1.01E-07 | -20.19 | 0.096 | 21.73 | 55.46 | 55.94 | 139.46 |
| 31 | 2.29 | 3.89 | 9.82 | 0.25 | 2.72E-07 | -10.74 | 0.2508 | 21.49 | 54.98 | 56.17 | 96.98 |
| 32 | 0.38 | 0.41 | 14.09 | 2.01 | 1.08E-05 | -15.19 | 0.0132 | 19.04 | 50.08 | 50.56 | 134.08 |
| 33 | 1.58 | 1.12 | 16.78 | 3.92 | 1.01E-06 | -8.5 | 0.0351 | 19.91 | 51.82 | 53.01 | 93.82 |
| 34 | 3.48 | 0.84 | 38.13 | 1.56 | 2.73E-08 | -6.05 | 0.0194 | 21.02 | 54.03 | 56.42 | 75.03 |
| 37 | 2.86 | 5.01 | 11.59 | 0.07 | 9.25E-08 | -15.93 | 0.068 | 27.77 | 67.53 | 70.92 | 81.53 |
| 40 | 2.83 | 1.52 | 2.02 | 16.53 | 1.53E-08 | -6.59 | 0.0467 | 21.07 | 54.13 | 55.31 | 96.13 |
| 41 | 2.14 | 0.78 | 1.33 | 3.22 | 2.85E-08 | -8.7 | 0.1206 | 21.38 | 54.77 | 55.95 | 96.77 |
| 42 | 3.32 | 1.02 | 6.86 | 0.24 | 8.26-09 | -6.08 | 0.0093 | 20.54 | 53.07 | 54.89 | 81.07 |
| 44 | 0.82 | 0.21 | 10.13 | 2.13 | 2.14E-06 | -6.6 | 0.0774 | 21.22 | 54.45 | 56.83 | 75.45 |
| 46 | 1.44 | 0.72 | 1.89 | 0.7 | 1.11E-07 | -20.06 | 0.0606 | 27.43 | 66.87 | 69.25 | 87.87 |
| 48 | 2.66 | 0.33 | 2.01 | 6.04 | 9.46E-09 | -11.53 | 0.0526 | 20.78 | 53.56 | 54.04 | 137.56 |
| 49 | 2.77 | 1.61 | 14.92 | 0.26 | 7.15E-08 | -5.53 | 0.0087 | 21.53 | 55.05 | 57.96 | 71.85 |
| 52 | 3.1 | 2.29 | 9.15 | 0.06 | 3.19E-08 | -2.23 | 0.1409 | 11.79 | 35.59 | 36.77 | 77.59 |
| 55 | 2.48 | 0.53 | 18 | 2.64 | 8.47E-08 | -13.21 | 0.0074 | 22.92 | 57.85 | 59.03 | 99.85 |
| 57 | 2.65 | 0.66 | 11.68 | 0.56 | 4.65E-08 | -8.72 | 0.0277 | 26.6 | 65.2 | 68.11 | 82 |
| 58 | 2.08 | 0.62 | 1.11 | 0.46 | 2.98E-08 | -7.99 | 0.0239 | 17.81 | 47.61 | 48.09 | 131.61 |
| 59 | 2.35 | 0.56 | 14.94 | 0.62 | 1.15E-07 | -7.04 | 0.0299 | 21 | 54 | 55.81 | 82 |
| 61 | 1.68 | 1.01 | 12.14 | 0.55 | 5.24E-07 | -4.28 | 0.0713 | 16.83 | 45.66 | 48.04 | 66.66 |
| 62 | 2.58 | 0.61 | 11.54 | 4.15 | 5.79E-08 | -6.17 | 0.0763 | 19.76 | 51.53 | 52.71 | 93.53 |
| 64 | 2.55 | 0.98 | 17.05 | 0.15 | 7.19E-08 | -20.39 | 0.0386 | 32.05 | 76.09 | 79.48 | 90.09 |
| 65 | 2.75 | 0.31 | 39.55 | 4.66 | 4.69E-08 | -49.59 | 0.2866 | 23.38 | 58.75 | 59.23 | 142.75 |
| 67 | 2.57 | 1.52 | 2.76 | 0.16 | 3.03E-08 | -5.56 | 0.0288 | 14.05 | 40.09 | 41.91 | 68.09 |
| 71 | 2.94 | 0.87 | 1.48 | 0.56 | 5.64E-09 | -9.88 | 0.0263 | 26.3 | 64.59 | 66.98 | 85.59 |
| 73 | 2.66 | 0.48 | 2.28 | 5.49 | 1.53E-08 | -3.85 | 0.0322 | 12.48 | 36.95 | 38.77 | 64.95 |

Table S5: Parameter value estimates for the Density-Dependent Cell Death & MOI model, along the negative log likelihood (nll), AIC, BIC and AICc.

| ID | $Log_{10}(\lambda)$ | α | η | k | β | γ | $t_0$ | error value | Negative log likelihood | AIC | BIC | AICc |
|---|---|---|---|---|---|---|---|---|---|---|---|---|
| 1 | 2.59 | 0.6 | 6.56 | 9.93 | 2.26E-08 | 1E-04 | -20.38 | 0.0638 | 29.56 | 73.12 | 75.9 | 110.45 |
| 2 | 0.33 | 0.1 | 9.92 | 48.46 | 5.88E-06 | 1E-04 | -16.5 | 0.037 | 22.73 | 59.46 | 60.85 | 171.46 |
| 4 | 0.9 | 0.24 | 19.08 | 24.91 | 2.55E-06 | 1E-04 | -7.55 | 0.0513 | 18.82 | 51.64 | 53.76 | 107.64 |
| 5 | 0 | 0.01 | 12.08 | 2.86 | 4.3E-05 | 6E-04 | -13.28 | 0.0547 | 18.11 | 50.21 | 50.77 | Inf |
| 6 | 3.79 | 4.32 | 11.81 | 3.57 | 9.49E-09 | 0.012 | -12.92 | 0.0607 | 19.42 | 52.83 | 53.39 | Inf |

| | | | | | | | | | | | | |
|---|---|---|---|---|---|---|---|---|---|---|---|---|
| 7 | 2.83 | 0.88 | 12.01 | 6.15 | 2.74E-08 | 0.0109 | -19 | 0.0226 | 29.02 | 72.04 | 75.43 | 100.04 |
| 8 | 3.91 | 5.76 | 11.88 | 3.71 | 9.17E-09 | 0.0268 | -3.56 | 0.0284 | 10.46 | 34.92 | 36.3 | 146.92 |
| 11 | 3.11 | 0.56 | 12.03 | 3.18 | 1.27E-08 | 5E-04 | -14.85 | 0.249 | 22.42 | 58.85 | 60.23 | 170.85 |
| 12 | 2.81 | 0.45 | 7.6 | 2.77 | 1.75E-08 | 6E-04 | -11.96 | 0.187 | 24.84 | 63.68 | 65.8 | 119.68 |
| 20 | 1.74 | 0.45 | 11.92 | 7.74 | 3.2E-07 | 1E-04 | -10.44 | 0.1921 | 18.6 | 51.2 | 51.75 | Inf |
| 21 | 3.22 | 0.72 | 9.11 | 3.65 | 9.96E-09 | 0.0127 | -8.98 | 0.1 | 21.24 | 56.48 | 57.86 | 168.48 |
| 22 | 2.96 | 4.31 | 11.59 | 6.99 | 6.645E-08 | 0.0266 | -5.5 | 0.0449 | 19.37 | 52.73 | 54.85 | 108.73 |
| 23 | 0.49 | 0.21 | 2.35 | 54.72 | 1.97E-06 | 1E-04 | -6.58 | 0.0701 | 18.01 | 50.01 | 51.39 | 162.01 |
| 24 | 2.87 | 2.99 | 11.9 | 1.32 | 5.08E-08 | 0.0177 | -10.77 | 0.0822 | 12.95 | 39.9 | 39.52 | -72.1 |
| 25 | 2.47 | 0.91 | 12.3 | 1.47 | 8.43E-08 | 1E-04 | -8.9 | 0.0417 | 19.89 | 53.78 | 55.16 | 165.78 |
| 26 | 2.18 | 0.21 | 11.53 | 23.92 | 9.52E-08 | 1E-04 | -10.4 | 0.0265 | 18.59 | 51.19 | 51.74 | Inf |
| 27 | 4.23 | 3.16 | 9.95 | 8.63 | 2.46E-09 | 0.0178 | -5.39 | 0.1364 | 17.76 | 49.52 | 50.9 | 161.52 |
| 28 | 2.71 | 0.92 | 12 | 20 | 4.34E-08 | 1E-04 | -12.03 | 0.06 | 24.63 | 63.26 | 65.37 | 119.26 |
| 29 | 2.08 | 0.35 | 12.04 | 3.34 | 1.08E-07 | 0.0079 | -17.69 | 0.0932 | 21.33 | 56.66 | 57.21 | Inf |
| 31 | 3.46 | 17.99 | 11.18 | 7.26 | 7.8E-08 | 0.0095 | -5.26 | 0.1416 | 17.18 | 48.37 | 49.75 | 160.37 |
| 32 | 0.13 | 0.37 | 7.77 | 89.85 | 1.52E-05 | 0.0361 | -15.12 | 0.0091 | 18.94 | 51.89 | 52.45 | Inf |
| 33 | 1.39 | 1.02 | 12.07 | 12.46 | 1.11E-06 | 1E-04 | -8.31 | 0.0409 | 19.75 | 53.51 | 54.89 | 165.51 |
| 34 | 2.68 | 0.95 | 9.11 | 8.93 | 3.40E-08 | 0.0176 | -11.98 | 0.0515 | 24.74 | 63.49 | 66.27 | 100.82 |
| 37 | 1.55 | 1.67 | 12.1 | 0.03 | 1.57E-06 | 0.001 | -2.25 | 0.2932 | 14.8 | 43.61 | 47.56 | 66.01 |
| 40 | 4.06 | 9.46 | 3.97 | 10.61 | 4.74E-09 | 0.011 | -5.72 | 0.0448 | 19.78 | 53.57 | 54.95 | 165.57 |
| 41 | 2.94 | 2.1 | 2.37 | 9.41 | 1.048E-08 | 0.0329 | -6.22 | 0.1143 | 19.26 | 52.53 | 53.91 | 164.53 |
| 42 | 3.61 | 1.18 | 12.06 | 0.35 | 6.67E-09 | 0.0313 | -5.13 | 0.0078 | 19.37 | 52.74 | 54.86 | 108.74 |
| 44 | 2.02 | 2.1 | 11.77 | 7.09 | 2.9E-07 | 0.067 | -2.72 | 0.0713 | 13.7 | 41.4 | 44.19 | 78.73 |
| 46 | 3.39 | 3.67 | 11.92 | 2.08 | 1.97E-08 | 0.0169 | -11.03 | 0.0477 | 23.72 | 61.44 | 64.22 | 98.77 |
| 48 | 5.06 | 6.2 | 11.44 | 8.48 | 7.08E-10 | 0.027 | -5.91 | 0.0741 | 17.07 | 48.15 | 48.7 | Inf |
| 49 | 2.6 | 1.58 | 11.97 | 1.14 | 7.17E-08 | 0.0302 | -4.63 | 0.0141 | 19.71 | 53.41 | 56.81 | 81.41 |
| 52 | 3.01 | 0.97 | 12 | 0.99 | 1.95E-08 | 5E-04 | -14.99 | 0.7707 | 21.74 | 57.48 | 58.86 | 169.48 |
| 55 | 2.8 | 1.37 | 11.99 | 1.36 | 4.09E-08 | 0.0254 | -10.11 | 0.0066 | 21.06 | 56.12 | 57.5 | 168.12 |
| 57 | 3.71 | 3.96 | 12.13 | 4.39 | 1.05E-08 | 0.0315 | -4.67 | 0.0161 | 21.11 | 56.23 | 59.62 | 84.23 |
| 58 | 2.91 | 0.31 | 11.96 | 1.67 | 2.36E-08 | 1E-04 | -7.74 | 0.0213 | 18.4 | 50.81 | 51.36 | Inf |
| 59 | 3.11 | 2.76 | 11.94 | 3.25 | 2.99E-08 | 0.0478 | -3.79 | 0.018 | 15.93 | 45.86 | 47.98 | 101.86 |
| 61 | 1.97 | 1.78 | 11.95 | 1.4 | 3.21E-07 | 0.0274 | -2.76 | 0.0542 | 12.78 | 39.55 | 42.34 | 76.89 |
| 62 | 2.98 | 1.04 | 12 | 1 | 2.6E-08 | 6E-04 | -9.97 | 0.2945 | 21.89 | 57.78 | 59.16 | 169.78 |
| 64 | 3.06 | 3.01 | 11.91 | 1.09 | 3.08E-08 | 0.0329 | -9.14 | 0.036 | 24.47 | 62.95 | 66.9 | 85.35 |
| 65 | 1.98 | 0.78 | 12.03 | 0.08 | 3.64E-07 | 7E-04 | -5.52 | 0.4364 | 16.86 | 47.72 | 48.27 | Inf |
| 67 | 3.55 | 2.79 | 11.99 | 2.89 | 1.1E-08 | 0.0692 | -4.16 | 0.0201 | 10.12 | 34.24 | 36.36 | 90.24 |
| 71 | 4.61 | 3.14 | 11.9 | 3.4 | 1.1E-09 | 0.0338 | -7.14 | 0.016 | 24.22 | 62.45 | 65.23 | 99.78 |
| 73 | 2.44 | 0.46 | 4.66 | 2.82 | 3.11E-08 | 7E-04 | -7.58 | 0.1742 | 17.94 | 49.89 | 52 | 105.89 |

Table S6: Summary of which model is selected by AIC, BIC and AICc for each study participant.

| ID | AIC | BIC | AICc | ID | AIC | BIC | AICc |
|---|---|---|---|---|---|---|---|
| 1 | Standard | Standard | Standard | 34 | DDDI | DDDI | Standard |
| 2 | DDDI | DDDI | Standard | 37 | DDDDI & MOI | DDDDI & MOI | DDDDI & MOI |
| 4 | DDDI | DDDI | Standard | 40 | Standard | Standard | Standard |
| 5 | Standard | Standard | Standard | 41 | DDDI | DDDI | Standard |
| 6 | DDDI | DDDI | Standard | 42 | Standard | Standard | Standard |
| 7 | Standard | Standard | Standard | 44 | DDDI | DDDI | Standard |
| 8 | Standard | Standard | Standard | 46 | DDDI | DDDI | Standard |
| 11 | DDDI | DDDI | Standard | 48 | DDDI | DDDI | Standard |
| 12 | DDDI | DDDI | Standard | 49 | Standard | Standard | Standard |
| 20 | DDDI | DDDI | Standard | 52 | Standard | Standard | Standard |
| 21 | DDDI | DDDI | Standard | 55 | DDDI | DDDI | Standard |
| 22 | Standard | Standard | Standard | 57 | DDDI | DDDI | Standard |
| 23 | DDDI | DDDI | Standard | 58 | Standard | Standard | Standard |
| 24 | Standard | Standard | DDDDI & MOI | 59 | DDDI | DDDI | Standard |
| 25 | Standard | Standard | Standard | 61 | DDDI | DDDI | Standard |
| 26 | DDDI | DDDI | Standard | 62 | DDDI | DDDI | Standard |
| 27 | DDDI | DDDI | Standard | 64 | DDDI | DDDI | DDDI |
| 28 | Standard | Standard | Standard | 65 | Standard | Standard | Standard |
| 29 | DDDI | DDDI | Standard | 67 | DDDI | DDDI | Standard |
| 31 | Standard | Standard | Standard | 71 | Standard | Standard | Standard |
| 32 | DDDI | DDDI | Standard | 73 | Standard | Standard | Standard |
| 33 | Standard | Standard | Standard | | | | |

Table S7: Data-derived growth rate and model-derived growth rates for each study participants, along with the squared difference for the data-and-model derived rates for each model.

| ID | Data Growth Rate | Standard Growth Rate | Error Standard | DDDI Growth Rate | Error DDDI | MOI Growth Rate | Error MOI | DDDDI & MOI Growth Rate | Error DDDDI & MOI | Best Model |
|---|---|---|---|---|---|---|---|---|---|---|
| 1 | 0.25 | 0.29 | 0.00147 | 0.29 | 0.0012 | 0.27 | 0.00037 | 0.27 | 0.00018 | DDDDI & MOI |
| 2 | 0.32 | 0.45 | 0.0162 | 0.44 | 0.01329 | 0.29 | 0.00129 | 0.29 | 0.00138 | DDDI |
| 4 | 0.28 | 0.31 | 0.00116 | 0.3 | 0.00075 | 0.29 | 0.00022 | 0.29 | 0.8E-4 | DDDDI & MOI |
| 5 | 0.19 | 1.08 | 0.78659 | 0.41 | 0.04914 | 0.32 | 0.01715 | 0.32 | 0.01746 | DDDI |
| 6 | 0.24 | 0.35 | 0.01297 | 0.37 | 0.01745 | 0.37 | 0.01892 | 0.34 | 0.01109 | DDDDI & MOI |
| 7 | 0.19 | 0.32 | 0.01845 | 0.26 | 0.00574 | 0.29 | 0.01108 | 0.29 | 0.00983 | MOI |
| 8 | 0.52 | 0.54 | 0.00035 | 0.49 | 0.00075 | 0.53 | 1E-04 | 0.49 | 0.00083 | DDDI |
| 11 | 0.35 | 0.53 | 0.03308 | 0.54 | 0.03493 | 0.59 | 0.0562 | 0.31 | 0.00195 | DDDDI & MOI |

| | | | | | | | | | | |
|---|---|---|---|---|---|---|---|---|---|---|
| 12 | 0.31 | 0.53 | 0.04923 | 0.57 | 0.06657 | 0.62 | 0.09704 | 0.38 | 0.00428 | DDDDI & MOI |
| 20 | 0.17 | 0.54 | 0.1396 | 0.51 | 0.11865 | 0.25 | 0.00734 | 0.39 | 0.04766 | DDDI |
| 21 | 0.43 | 0.54 | 0.01167 | 0.53 | 0.00987 | 0.48 | 0.00291 | 0.43 | 0.3E-4 | DDDDI & MOI |
| 22 | 0.32 | 0.66 | 0.11254 | 0.53 | 0.04211 | 0.46 | 0.02031 | 0.53 | 0.04525 | DDDI |
| 23 | 0.38 | 0.62 | 0.05572 | 0.53 | 0.02196 | 0.4 | 0.00034 | 0.4 | 0.00057 | DDDI |
| 24 | 0.14 | 0.16 | 0.2E-3 | 0.16 | 0.00036 | 0.16 | 0.00023 | 0.21 | 0.004 | Standard |
| 25 | 0.48 | 0.45 | 0.00074 | 0.45 | 0.00072 | 0.42 | 0.00331 | 0.41 | 0.00421 | MOI |
| 26 | 0.4 | 0.41 | 0.4E-4 | 0.48 | 0.00578 | 0.39 | 0.00017 | 0.41 | 1.06E-6 | DDDDI & MOI |
| 27 | 0.37 | 0.48 | 0.01265 | 0.49 | 0.01524 | 0.43 | 0.00415 | 0.49 | 0.01508 | DDDI |
| 28 | 0.26 | 0.34 | 0.00537 | 0.35 | 0.0081 | 0.32 | 0.00354 | 0.35 | 0.00758 | DDDI |
| 29 | 0.19 | 0.33 | 0.01896 | 0.43 | 0.06048 | 0.27 | 0.0069 | 0.3 | 0.01169 | DDDI |
| 31 | 0.13 | 0.59 | 0.21668 | 0.28 | 0.02266 | 0.4 | 0.0751 | 0.56 | 0.18672 | MOI |
| 32 | 0.21 | 0.22 | 2E-04 | 0.29 | 0.00765 | 0.26 | 0.00257 | 0.25 | 0.00198 | Standard |
| 33 | 0.43 | 0.4 | 0.00086 | 0.42 | 0.00028 | 0.4 | 0.00129 | 0.39 | 0.00161 | MOI |
| 34 | 0.46 | 0.53 | 0.00505 | 0.53 | 0.0048 | 0.55 | 0.00836 | 0.36 | 0.00937 | MOI |
| 37 | 0.11 | 0.17 | 0.00339 | 0.16 | 0.00263 | 0.21 | 0.01003 | 0.83 | 0.51031 | MOI |
| 40 | 0.48 | 0.63 | 0.02104 | 0.45 | 0.00086 | 0.62 | 0.02002 | 0.64 | 0.02427 | MOI |
| 41 | 0.32 | 0.49 | 0.02946 | 0.47 | 0.02329 | 0.41 | 0.00754 | 0.47 | 0.02246 | DDDI |
| 42 | 0.46 | 0.51 | 0.00229 | 0.51 | 0.00227 | 0.48 | 0.00045 | 0.52 | 0.00294 | DDDI |
| 44 | 0.33 | 0.48 | 0.02197 | 0.43 | 0.0094 | 0.38 | 0.0018 | 0.45 | 0.01241 | DDDI |
| 46 | 0.16 | 0.23 | 0.00556 | 0.27 | 0.01344 | 0.21 | 0.00314 | 0.26 | 0.01064 | DDDI |
| 48 | 0.34 | 0.52 | 0.03033 | 0.62 | 0.07535 | 0.46 | 0.01402 | 0.65 | 0.09556 | DDDI |
| 49 | 0.43 | 0.44 | 0.00033 | 0.44 | 0.2E-4 | 0.43 | 0.32E-4 | 0.42 | 0.1E-4 | DDDDI & MOI |
| 52 | 0.29 | 0.65 | 0.12639 | 0.64 | 0.1222 | 0.64 | 0.12419 | 0.26 | 0.00077 | DDDDI & MOI |
| 55 | 0.34 | 0.37 | 0.00127 | 0.41 | 0.00487 | 0.36 | 0.00042 | 0.39 | 0.00307 | DDDI |
| 57 | 0.39 | 0.47 | 0.00509 | 0.44 | 0.00185 | 0.42 | 0.00049 | 0.45 | 0.00261 | DDDI |
| 58 | 0.45 | 0.62 | 0.03128 | 0.56 | 0.01339 | 0.44 | 4.23E-9 | 0.49 | 0.00228 | DDDI |
| 59 | 0.36 | 0.47 | 0.01031 | 0.46 | 0.00997 | 0.44 | 0.00619 | 0.47 | 0.01022 | DDDI |
| 61 | 0.28 | 0.38 | 0.00944 | 0.36 | 0.00653 | 0.36 | 0.00649 | 0.37 | 0.00805 | DDDI |
| 62 | 0.54 | 0.53 | 0.00016 | 0.54 | 0.2E-4 | 0.49 | 0.00334 | 0.38 | 0.02616 | MOI |
| 64 | 0.16 | 0.2 | 0.00119 | 0.22 | 0.00369 | 0.2 | 0.00112 | 0.24 | 0.00576 | DDDI |
| 65 | 0.32 | 0.15 | 0.02861 | 0.24 | 0.00573 | 0.14 | 0.03127 | 0.73 | 0.16905 | MOI |
| 67 | 0.55 | 0.61 | 0.00278 | 0.57 | 0.00022 | 0.55 | 0.9E-4 | 0.61 | 0.00252 | DDDI |
| 71 | 0.44 | 0.49 | 0.00253 | 0.51 | 0.00439 | 0.46 | 0.00044 | 0.52 | 0.00575 | DDDI |
| 73 | 0.6 | 0.65 | 0.00188 | 0.64 | 0.0012 | 0.61 | 0.3E-4 | 0.45 | 0.0239 | DDDI |

Table S8: Data-derived decay rate and model-derived decay rates for each study participants, along with the squared difference for the data-and-model derived rates for each model.

| ID | Data Decay Rate | Standard Decay Rate | Error Standard | DDDI Decay Rate | Error DDDI | MOI Decay Rate | Error MOI | DDDDI & MOI Decay Rate | Error DDDDI & MOI | Best Model |
|---|---|---|---|---|---|---|---|---|---|---|
| 1 | -0.13 | -0.13 | 1.36E-6 | -0.17 | 0.00166 | -0.15 | 0.00025 | -0.15 | 0.00026 | Standard |
| 2 | -0.14 | -0.11 | 0.00069 | -0.15 | 0.00028 | -0.15 | 0.00028 | -0.2 | 0.00438 | MOI |
| 4 | -0.21 | -0.1 | 0.01223 | -0.14 | 0.00591 | -0.15 | 0.00423 | -0.17 | 0.00154 | DDDDI & MOI |
| 5 | -0.19 | -0.16 | 0.00099 | -0.17 | 0.00037 | -0.13 | 0.00425 | -0.22 | 6E-04 | MOI |
| 6 | -0.24 | -0.15 | 0.00664 | -0.17 | 0.00388 | -0.2 | 0.00136 | -0.17 | 0.00484 | DDDI |
| 7 | -0.11 | -0.14 | 0.00093 | -0.14 | 0.00136 | -0.15 | 0.00172 | -0.15 | 0.00167 | Standard |
| 8 | -0.15 | -0.13 | 0.00036 | -0.18 | 0.00082 | -0.17 | 0.00027 | -0.17 | 0.00033 | DDDI |
| 11 | -0.18 | -0.23 | 0.0026 | -0.24 | 0.00289 | -0.27 | 0.00723 | -0.16 | 0.00049 | DDDDI & MOI |
| 12 | -0.24 | -0.25 | 0.00013 | -0.25 | 7E-05 | -0.25 | 0.00028 | -0.19 | 0.00256 | MOI |
| 20 | -0.35 | -0.17 | 0.02955 | -0.23 | 0.01363 | -0.14 | 0.04168 | -0.2 | 0.02206 | MOI |
| 21 | -0.26 | -0.24 | 0.00043 | -0.23 | 0.00105 | -0.25 | 0.00017 | -0.22 | 0.00219 | DDDI |
| 22 | -0.18 | -0.16 | 4E-04 | -0.24 | 0.00285 | -0.2 | 0.00016 | -0.23 | 0.0025 | DDDI |
| 23 | -0.21 | -0.13 | 0.00649 | -0.18 | 8E-04 | -0.2 | 0.00012 | -0.35 | 0.01963 | DDDI |
| 24 | NA | NA | NA | NA | NA | NA | NA | NA | NA | NA |
| 25 | -0.15 | -0.18 | 0.00101 | -0.18 | 0.00107 | -0.18 | 0.00136 | -0.19 | 0.00168 | Standard |
| 26 | -0.19 | -0.12 | 0.00582 | -0.16 | 0.00092 | -0.25 | 0.00323 | -0.25 | 0.00334 | MOI |
| 27 | -0.3 | -0.25 | 0.00292 | -0.25 | 0.00229 | -0.28 | 0.00034 | -0.25 | 0.00301 | DDDI |
| 28 | -0.22 | -0.19 | 0.00055 | -0.22 | 1E-05 | -0.21 | 0.00012 | -0.2 | 4E-04 | MOI |
| 29 | -0.27 | -0.12 | 0.02153 | -0.15 | 0.01289 | -0.13 | 0.01745 | -0.14 | 0.01592 | MOI |
| 31 | -0.27 | -0.27 | 1.05E-7 | -0.19 | 0.00718 | -0.2 | 0.00484 | -0.31 | 0.00121 | Standard |
| 32 | -0.16 | -0.1 | 0.00394 | -0.12 | 0.00196 | -0.13 | 0.00094 | -0.16 | 1E-05 | DDDDI & MOI |
| 33 | -0.17 | -0.21 | 0.00189 | -0.21 | 0.00223 | -0.22 | 0.00291 | -0.23 | 0.0043 | Standard |
| 34 | -0.16 | -0.23 | 0.00398 | -0.22 | 0.0037 | -0.25 | 0.00691 | -0.18 | 0.00034 | DDDDI & MOI |
| 37 | -0.05 | -0.07 | 0.00036 | -0.08 | 0.00102 | -0.09 | 0.00154 | -0.05 | 1.55E-6 | DDDDI & MOI |
| 40 | -0.26 | -0.37 | 0.01159 | -0.3 | 0.00157 | -0.39 | 0.01608 | -0.37 | 0.01339 | MOI |
| 41 | -0.24 | -0.26 | 0.00031 | -0.25 | 1E-05 | -0.24 | 1.99E-6 | -0.25 | 0.00014 | DDDI |
| 42 | -0.11 | -0.1 | 0.00018 | -0.11 | 1.67E-6 | -0.14 | 0.00043 | -0.13 | 0.00026 | MOI |
| 44 | -0.16 | -0.09 | 0.00413 | -0.15 | 1E-04 | -0.19 | 0.001 | -0.14 | 0.00045 | MOI |
| 46 | -0.15 | -0.09 | 0.00306 | -0.11 | 0.00154 | -0.1 | 0.00189 | -0.12 | 0.00061 | DDDDI & MOI |
| 48 | -0.35 | -0.19 | 0.02578 | -0.23 | 0.01432 | -0.31 | 0.00173 | -0.24 | 0.01151 | DDDI |
| 49 | -0.11 | -0.12 | 0.00019 | -0.14 | 0.00064 | -0.14 | 0.00082 | -0.17 | 0.00326 | Standard |
| 52 | -0.07 | -0.06 | 9E-05 | -0.09 | 0.00038 | -0.08 | 2E-05 | -0.13 | 0.00356 | DDDI |

| | | | | | | | | | |
|---|---|---|---|---|---|---|---|---|---|
| 55 | -0.13 | -0.14 | 0.5E-4 | -0.18 | 0.00189 | -0.18 | 0.00246 | -0.16 | 0.00082 | Standard |
| 57 | -0.1 | -0.11 | 1E-04 | -0.16 | 0.00424 | -0.16 | 0.00334 | -0.16 | 0.00357 | Standard |
| 58 | -0.14 | -0.13 | 0.00024 | -0.16 | 0.00056 | -0.17 | 0.00089 | -0.22 | 0.00685 | Standard |
| 59 | -0.15 | -0.12 | 0.00098 | -0.16 | 5E-05 | -0.15 | 2.96E-6 | -0.15 | 3E-05 | DDDI |
| 61 | -0.26 | -0.13 | 0.01501 | -0.16 | 0.00815 | -0.14 | 0.01289 | -0.16 | 0.0096 | MOI |
| 62 | -0.18 | -0.2 | 0.00028 | -0.26 | 0.0058 | -0.25 | 0.00559 | -0.18 | 1E-05 | DDDDI & MOI |
| 64 | -0.05 | -0.05 | 1.6E-6 | -0.06 | 0.00022 | -0.05 | 6E-05 | -0.06 | 0.00017 | Standard |
| 65 | NA | NA | NA | NA | NA | NA | NA | NA | NA | NA |
| 67 | -0.11 | -0.12 | 2E-05 | -0.16 | 0.00219 | -0.14 | 0.00106 | -0.17 | 0.00295 | Standard |
| 71 | -0.14 | -0.16 | 0.00024 | -0.19 | 0.00194 | -0.21 | 0.00419 | -0.18 | 0.00141 | Standard |
| 73 | -0.18 | -0.18 | 2E-05 | -0.27 | 0.00801 | -0.42 | 0.05464 | -0.24 | 0.00388 | Standard |

Table S9: Data-derived setpoint and model-derived setpoint for each study participants, along with the squared difference for the data-and-model derived setpoint for each model.

| ID | Data Setpoint | Standard Setpoint | Error Standard | DDDI Setpoint | Error DDDI | MOI Setpoint | Error MOI | DDDDI & MOI Setpoint | Error DDDDI & MOI | Model |
|---|---|---|---|---|---|---|---|---|---|---|
| 1 | 3.9 | 5.69 | 3.18 | 5.75 | 3.4 | 5.72 | 3.28 | 5.72 | 3.3 | Standard |
| 2 | 3.69 | 4.14 | 0.2 | 4.21 | 0.26 | 4.17 | 0.23 | 4.25 | 0.31 | Standard |
| 4 | 3.81 | 4 | 0.04 | 4.03 | 0.05 | 4.08 | 0.08 | 4.09 | 0.08 | Standard |
| 5 | 4.3 | 5.25 | 0.89 | 5.94 | 2.68 | 4.76 | 0.21 | 4.62 | 0.1 | DDDDI & MOI |
| 6 | 4.9 | 5.44 | 0.3 | 5.43 | 0.29 | 5.63 | 0.53 | 5.39 | 0.24 | DDDDI & MOI |
| 7 | 3.38 | 5.47 | 4.37 | 5.43 | 4.22 | 5.5 | 4.48 | 5.47 | 4.4 | DDDI |
| 8 | 3.9 | 5.51 | 2.57 | 5.48 | 2.5 | 5.58 | 2.81 | 5.47 | 2.44 | DDDDI & MOI |
| 11 | NA | NA | NA | NA | NA | NA | NA | NA | NA | NA |
| 12 | 5.2 | 6.52 | 1.75 | 6.42 | 1.48 | 6.49 | 1.67 | 6.1 | 0.81 | DDDDI & MOI |
| 20 | 3.46 | 4.9 | 2.07 | 4.94 | 2.16 | 4.64 | 1.38 | 4.84 | 1.88 | MOI |
| 21 | 5.08 | 6.48 | 1.98 | 6.34 | 1.59 | 6.41 | 1.77 | 6.21 | 1.29 | DDDDI & MOI |
| 22 | 3.2 | 4.73 | 2.33 | 4.75 | 2.41 | 4.7 | 2.25 | 4.77 | 2.45 | MOI |
| 23 | 4.08 | 4.46 | 0.15 | 4.48 | 0.17 | 4.43 | 0.12 | 4.7 | 0.38 | MOI |
| 24 | NA | NA | NA | NA | NA | NA | NA | NA | NA | NA |
| 25 | 3.57 | 5.13 | 2.43 | 5.13 | 2.44 | 5.14 | 2.45 | 5.15 | 2.48 | Standard |
| 26 | 4.54 | 5.48 | 0.89 | 5.6 | 1.14 | 5.68 | 1.32 | 5.7 | 1.35 | Standard |
| 27 | 3.7 | 6.38 | 7.18 | 6.32 | 6.83 | 6.4 | 7.26 | 6.26 | 6.56 | DDDDI & MOI |
| 28 | 3.09 | 5.29 | 4.84 | 5.37 | 5.21 | 5.37 | 5.24 | 5.36 | 5.16 | Standard |
| 29 | 4.32 | 5.24 | 0.85 | 5.31 | 0.98 | 5.23 | 0.83 | 5.28 | 0.93 | MOI |
| 31 | 2.69 | 4.49 | 3.23 | 3.96 | 1.6 | 4.12 | 2.04 | 4.36 | 2.77 | DDDI |

| | | | | | | | | | |
|---|---|---|---|---|---|---|---|---|---|
| 32 | 3.45 | 3.23 | 0.05 | 3.28 | 0.03 | 3.37 | 0.01 | 3.48 | 0 | DDDDI & MOI |
| 33 | NA | NA | NA | NA | NA | NA | NA | NA | NA | NA |
| 34 | NA | NA | NA | NA | NA | NA | NA | NA | NA | NA |
| 37 | 3.02 | 4.1 | 1.15 | 4.07 | 1.1 | 4.19 | 1.35 | 3.93 | 0.82 | DDDDI & MOI |
| 40 | NA | NA | NA | NA | NA | NA | NA | NA | NA | NA |
| 41 | NA | NA | NA | NA | NA | NA | NA | NA | NA | NA |
| 42 | 5.29 | 6.16 | 0.76 | 6.24 | 0.92 | 6.22 | 0.87 | 6.23 | 0.9 | Standard |
| 44 | 3.78 | 4.3 | 0.27 | 4.32 | 0.29 | 4.39 | 0.37 | 4.35 | 0.32 | Standard |
| 46 | 4.77 | 4.97 | 0.04 | 5 | 0.05 | 5.01 | 0.06 | 5.04 | 0.07 | Standard |
| 48 | 5.22 | 6.63 | 1.97 | 6.76 | 2.37 | 6.74 | 2.31 | 6.67 | 2.09 | Standard |
| 49 | 4.8 | 4.96 | 0.03 | 4.99 | 0.04 | 5 | 0.04 | 5.01 | 0.05 | Standard |
| 52 | 5.34 | 5.42 | 0.01 | 5.61 | 0.07 | 5.42 | 0.01 | 5.55 | 0.04 | Standard |
| 55 | 4.01 | 5.21 | 1.44 | 5.26 | 1.56 | 5.29 | 1.62 | 5.26 | 1.56 | Standard |
| 57 | 4.6 | 5.47 | 0.76 | 5.5 | 0.82 | 5.54 | 0.88 | 5.49 | 0.8 | Standard |
| 58 | 5.03 | 6.13 | 1.21 | 6.23 | 1.44 | 6.13 | 1.2 | 6.22 | 1.42 | MOI |
| 59 | 4.8 | 5.21 | 0.17 | 5.2 | 0.17 | 5.23 | 0.19 | 5.21 | 0.17 | DDDI |
| 61 | 3.71 | 4.28 | 0.33 | 4.25 | 0.3 | 4.27 | 0.31 | 4.26 | 0.3 | DDDI |
| 62 | 4.93 | 5.47 | 0.29 | 5.5 | 0.33 | 5.56 | 0.4 | 5.57 | 0.41 | Standard |
| 64 | 4.41 | 4.81 | 0.16 | 4.81 | 0.16 | 4.82 | 0.17 | 4.8 | 0.15 | DDDDI & MOI |
| 65 | 4.09 | 5.37 | 1.64 | 4.85 | 0.58 | 5.35 | 1.58 | 4.81 | 0.52 | DDDDI & MOI |
| 67 | 5.08 | 5.76 | 0.47 | 5.77 | 0.48 | 5.72 | 0.42 | 5.75 | 0.45 | MOI |
| 71 | 5.63 | 6.61 | 0.96 | 6.62 | 0.97 | 6.69 | 1.11 | 6.6 | 0.93 | DDDDI & MOI |
| 73 | 4.33 | 6.14 | 3.27 | 6.38 | 4.23 | 6.53 | 4.86 | 5.97 | 2.71 | DDDDI & MOI |

Table S10: Data-derived peak magnitude and model-derived peak magnitude for each study participants, along with the squared difference for the data-and-model derived peak magnitude for each model.

| ID | Data Peak magnitude | Standard Peak magnitude | Error Standard | DDDI Peak magnitude | Error DDDI | MOI Peak magnitude | SQ_MOI | DDDDI & MOI Peak magnitude | Error DDDDI & MOI | Best Model |
|---|---|---|---|---|---|---|---|---|---|---|
| 1 | 7.51 | 6.99 | 0.2708 | 7.1 | 0.17 | 7.06 | 0.204 | 7.06 | 0.2005 | DDDI |
| 2 | 5.71 | 5.41 | 0.0884 | 5.47 | 0.0552 | 5.4 | 0.0961 | 5.5 | 0.045 | DDDDI & MOI |
| 4 | 5.94 | 5.22 | 0.5122 | 5.28 | 0.4361 | 5.4 | 0.2923 | 5.39 | 0.2968 | MOI |
| 5 | 5.74 | 6.74 | 0.9936 | 7.3 | 2.4102 | 5.97 | 0.0504 | 5.7 | 0.0022 | DDDDI & MOI |
| 6 | 7.27 | 6.76 | 0.2504 | 6.8 | 0.2206 | 7.04 | 0.0516 | 6.74 | 0.2752 | MOI |
| 7 | 6.78 | 6.78 | 2.5E-8 | 6.72 | 0.0039 | 6.85 | 0.0042 | 6.8 | 4E-04 | Standard |

| | | | | | | | | | | |
|---|---|---|---|---|---|---|---|---|---|---|
| 8 | 7.25 | 6.85 | 0.1573 | 6.81 | 0.1892 | 7 | 0.0618 | 6.8 | 0.1955 | MOI |
| 11 | 7.78 | 7.98 | 0.0377 | 7.82 | 0.0012 | 8.26 | 0.233 | 7.4 | 0.148 | DDDI |
| 12 | 8.07 | 7.97 | 0.0098 | 7.88 | 0.0368 | 8.06 | 3E-04 | 7.52 | 0.3049 | MOI |
| 20 | 6.68 | 6.32 | 0.1293 | 6.39 | 0.0852 | 5.96 | 0.5221 | 6.28 | 0.1564 | DDDI |
| 21 | 8.27 | 7.93 | 0.1181 | 7.73 | 0.2899 | 7.87 | 0.1593 | 7.67 | 0.3559 | Standard |
| 22 | 6.32 | 6.16 | 0.0246 | 6.2 | 0.0131 | 6.18 | 0.0196 | 6.22 | 0.0095 | DDDDI & MOI |
| 23 | 6.26 | 5.8 | 0.2108 | 5.84 | 0.1748 | 5.77 | 0.242 | 6.08 | 0.0336 | DDDDI & MOI |
| 24 | 6.01 | 5.69 | 0.1034 | 5.67 | 0.1149 | 5.68 | 0.1067 | 5.57 | 0.1954 | Standard |
| 25 | 6.74 | 6.55 | 0.0383 | 6.55 | 0.0369 | 6.59 | 0.024 | 6.61 | 0.0161 | DDDDI & MOI |
| 26 | 7.39 | 6.75 | 0.4078 | 6.9 | 0.24 | 7.12 | 0.0745 | 7.15 | 0.0607 | DDDDI & MOI |
| 27 | 8.29 | 7.84 | 0.2078 | 7.79 | 0.2496 | 7.89 | 0.1591 | 7.76 | 0.2846 | MOI |
| 28 | 7.07 | 6.71 | 0.1283 | 6.82 | 0.0642 | 6.81 | 0.0699 | 6.83 | 0.0571 | DDDDI & MOI |
| 29 | 7.01 | 6.5 | 0.2626 | 6.58 | 0.1895 | 6.51 | 0.249 | 6.58 | 0.1875 | DDDDI & MOI |
| 31 | 5.68 | 6 | 0.1043 | 5.35 | 0.1068 | 5.58 | 0.0106 | 5.93 | 0.0606 | MOI |
| 32 | 4.84 | 4.44 | 0.1532 | 4.48 | 0.1273 | 4.56 | 0.0781 | 4.58 | 0.0634 | DDDDI & MOI |
| 33 | 5.64 | 5.36 | 0.0805 | 5.41 | 0.0549 | 5.48 | 0.025 | 5.51 | 0.017 | DDDDI & MOI |
| 34 | 7.24 | 7.3 | 0.0044 | 7.21 | 7E-04 | 7.32 | 0.006 | 6.87 | 0.135 | DDDI |
| 37 | 5.45 | 5.16 | 0.0814 | 5.15 | 0.0882 | 5.32 | 0.0157 | 4.9 | 0.302 | MOI |
| 40 | 7.76 | 7.71 | 0.0023 | 7.17 | 0.3442 | 7.84 | 0.0072 | 7.72 | 0.0016 | DDDDI & MOI |
| 41 | 7.69 | 7.65 | 0.0019 | 7.42 | 0.0721 | 7.44 | 0.0629 | 7.47 | 0.0476 | Standard |
| 42 | 7.67 | 7.41 | 0.0686 | 7.47 | 0.0418 | 7.55 | 0.0152 | 7.53 | 0.0194 | MOI |
| 44 | 5.95 | 5.53 | 0.1799 | 5.57 | 0.1463 | 5.73 | 0.0471 | 5.59 | 0.1291 | MOI |
| 46 | 6.53 | 6.14 | 0.1555 | 6.2 | 0.1106 | 6.18 | 0.1226 | 6.25 | 0.0783 | DDDDI & MOI |
| 48 | 8.54 | 8.01 | 0.2802 | 8.12 | 0.1725 | 8.24 | 0.0856 | 8.12 | 0.1695 | MOI |
| 49 | 6.46 | 6.27 | 0.0369 | 6.28 | 0.0334 | 6.35 | 0.0121 | 6.36 | 0.0118 | DDDDI & MOI |
| 52 | 6.66 | 6.56 | 0.0104 | 6.68 | 6E-04 | 6.62 | 0.0018 | 6.84 | 0.034 | DDDI |
| 55 | 6.79 | 6.55 | 0.0616 | 6.59 | 0.0399 | 6.69 | 0.0107 | 6.61 | 0.0339 | MOI |
| 57 | 6.86 | 6.74 | 0.0128 | 6.8 | 0.0032 | 6.9 | 0.002 | 6.8 | 0.0032 | MOI |
| 58 | 7.69 | 7.48 | 0.0429 | 7.53 | 0.0244 | 7.46 | 0.051 | 7.69 | 3.1E-5 | DDDDI & MOI |
| 59 | 6.77 | 6.51 | 0.0689 | 6.48 | 0.0875 | 6.6 | 0.0283 | 6.49 | 0.0805 | MOI |
| 61 | 6.14 | 5.59 | 0.3029 | 5.57 | 0.3225 | 5.62 | 0.27 | 5.58 | 0.3142 | MOI |
| 62 | 7.13 | 6.92 | 0.043 | 6.96 | 0.0278 | 7.1 | 8E-04 | 7 | 0.0164 | MOI |
| 64 | 6.09 | 5.77 | 0.1053 | 5.75 | 0.1136 | 5.81 | 0.0808 | 5.74 | 0.1196 | MOI |
| 65 | 6.95 | 6.4 | 0.3055 | 5.87 | 1.1796 | 6.42 | 0.288 | 6 | 0.9118 | MOI |

| | | | | | | | | | |
|---|---|---|---|---|---|---|---|---|---|
| 67 | 7.16 | 7.08 | 0.007 | 7.03 | 0.018 | 7.07 | 0.0093 | 7.03 | 0.0165 | Standard |
| 71 | 8.21 | 7.97 | 0.0605 | 7.94 | 0.0731 | 8.09 | 0.0142 | 7.96 | 0.061 | MOI |
| 73 | 7.89 | 7.57 | 0.0998 | 7.78 | 0.0122 | 8.14 | 0.0663 | 7.46 | 0.1803 | DDDI |

Table S11: Data-derived peak time and model-derived peak time for each study participants, along with the squared difference for the data-and-model derived peak time for each model.

| ID | Data peak time | Standard peak time | Error Standard | DDDI peak time | SQ_DDDI | MOI peak time | Error MOI | DDDDI & MOI peak time | Error DDDDI & MOI | Best Model |
|---|---|---|---|---|---|---|---|---|---|---|
| 1 | 11 | 9.03 | 3.881 | 14.08 | 9.49 | 11.47 | 0.221 | 11.63 | 0.397 | MOI |
| 2 | 7.41 | 6.27 | 1.292 | 8.74 | 1.78 | 8.99 | 2.507 | 9.31 | 3.623 | Standard |
| 4 | 18.91 | 15.95 | 8.783 | 17.84 | 1.15 | 17.9 | 1.027 | 18.29 | 0.389 | DDDDI & MOI |
| 5 | 12.17 | 2.47 | 94.183 | 0.1 | 145.8 | 4.43 | 59.982 | 11.77 | 0.164 | DDDDI & MOI |
| 6 | 11.34 | 9.1 | 5.01 | 10.32 | 1.04 | 9.81 | 2.336 | 9.94 | 1.955 | DDDI |
| 7 | 9.28 | 8.7 | 0.337 | 10.02 | 0.55 | 10.37 | 1.187 | 10.49 | 1.463 | Standard |
| 8 | 9.83 | 10.24 | 0.165 | 12.72 | 8.33 | 10.72 | 0.785 | 12.05 | 4.912 | Standard |
| 11 | 14.01 | 12.76 | 1.552 | 14.04 | 0.001 | 12.23 | 3.153 | 14.54 | 0.285 | DDDI |
| 12 | 12.92 | 11.49 | 2.043 | 13.15 | 0.05 | 10.67 | 5.06 | 11.81 | 1.231 | DDDI |
| 20 | 13 | 7.62 | 28.944 | 10.61 | 5.71 | 9.99 | 9.06 | 10.15 | 8.123 | DDDI |
| 21 | 12.12 | 11.65 | 0.217 | 12.46 | 0.12 | 12.04 | 0.006 | 12.23 | 0.013 | MOI |
| 22 | 8.1 | 5.91 | 4.781 | 8.98 | 0.78 | 7.74 | 0.127 | 8.3 | 0.041 | DDDDI & MOI |
| 23 | 11.03 | 8.01 | 9.134 | 11.51 | 0.23 | 11.17 | 0.019 | 11.63 | 0.357 | MOI |
| 24 | 22 | 28.95 | 48.303 | 27.06 | 25.6 | 22.98 | 0.96 | 22.27 | 0.073 | DDDDI & MOI |
| 25 | 8.93 | 10.35 | 2.018 | 10.44 | 2.28 | 10.87 | 3.766 | 11.02 | 4.371 | Standard |
| 26 | 10.52 | 9.74 | 0.608 | 11.9 | 1.9 | 11.51 | 0.98 | 11.38 | 0.74 | Standard |
| 27 | 13.89 | 13.01 | 0.773 | 14.39 | 0.25 | 13.25 | 0.408 | 13.01 | 0.773 | DDDI |
| 28 | 12.63 | 11.02 | 2.589 | 12.84 | 0.04 | 12.05 | 0.335 | 11.94 | 0.475 | DDDI |
| 29 | 12.85 | 8.33 | 20.4 | 11.15 | 2.88 | 10.38 | 6.084 | 10.36 | 6.183 | DDDI |
| 31 | 9 | 5.72 | 10.758 | 7.79 | 1.46 | 6.13 | 8.237 | 6.71 | 5.244 | DDDI |
| 32 | 10.72 | 6.26 | 19.858 | 10.87 | 0.02 | 10.36 | 0.127 | 10.93 | 0.046 | DDDI |
| 33 | 8 | 8.2 | 0.04 | 8.63 | 0.4 | 9.39 | 1.932 | 9.59 | 2.528 | Standard |
| 34 | 10 | 10.97 | 0.934 | 11.1 | 1.2 | 10.91 | 0.821 | 11.26 | 1.578 | MOI |
| 37 | 17 | 10.19 | 46.376 | 15.52 | 2.19 | 14.9 | 4.41 | 6.23 | 115.993 | DDDI |
| 40 | 9 | 7.39 | 2.592 | 7.72 | 1.64 | 7.58 | 2.016 | 7.68 | 1.742 | DDDI |
| 41 | 12.55 | 11.26 | 1.666 | 13.01 | 0.21 | 12.01 | 0.292 | 11.98 | 0.326 | DDDI |
| 42 | 11.54 | 11.39 | 0.023 | 11.75 | 0.04 | 12.55 | 1.018 | 12.73 | 1.414 | Standard |
| 44 | 12.68 | 10.1 | 6.658 | 13.97 | 1.66 | 13.21 | 0.281 | 12.93 | 0.062 | DDDDI & MOI |
| 46 | 15.74 | 11.13 | 21.272 | 15.38 | 0.13 | 14.23 | 2.287 | 14.35 | 1.938 | DDDI |

| | | | | | | | | | |
|---|---|---|---|---|---|---|---|---|---|
| 48 | 9.56 | 8.18 | 1.903 | 8.75 | 0.66 | 8.81 | 0.562 | 8.36 | 1.439 | MOI |
| 49 | 11.53 | 11.86 | 0.11 | 12.26 | 0.54 | 12.96 | 2.052 | 13.91 | 5.675 | Standard |
| 52 | 11 | 9.26 | 3.028 | 9.97 | 1.06 | 10.64 | 0.13 | 17.19 | 38.316 | MOI |
| 55 | 9.47 | 9.78 | 0.098 | 11.82 | 5.54 | 10.62 | 1.331 | 10.66 | 1.425 | Standard |
| 57 | 10.39 | 9.57 | 0.671 | 13.58 | 10.18 | 11.77 | 1.906 | 12.65 | 5.111 | Standard |
| 58 | 10.39 | 9.61 | 0.612 | 10.78 | 0.15 | 11.4 | 1.015 | 11.24 | 0.718 | DDDI |
| 59 | 12.16 | 11.36 | 0.635 | 13.43 | 1.62 | 11.85 | 0.094 | 12.78 | 0.388 | MOI |
| 61 | 17.4 | 13.6 | 14.427 | 16.75 | 0.42 | 15.53 | 3.491 | 15.98 | 2.012 | DDDI |
| 62 | 9.51 | 10.32 | 0.662 | 12.81 | 10.91 | 11.55 | 4.175 | 12.33 | 7.971 | Standard |
| 64 | 18 | 15.52 | 6.15 | 19.96 | 3.84 | 18.56 | 0.314 | 19.58 | 2.496 | MOI |
| 65 | 7.3 | 8.82 | 2.323 | 7.73 | 0.19 | 11.69 | 19.307 | 5.23 | 4.268 | DDDI |
| 67 | 8.13 | 8.59 | 0.21 | 10.77 | 6.96 | 9.7 | 2.46 | 10.24 | 4.445 | Standard |
| 71 | 9.03 | 9.48 | 0.2 | 11.58 | 6.49 | 10.07 | 1.076 | 10.88 | 3.413 | Standard |
| 73 | 9.88 | 9.88 | 2.76E-7 | 11.06 | 1.39 | 11.26 | 1.906 | 11.93 | 4.205 | Standard |

Table S12: Data-derived joint peak measurement and model-derived joint peak measurement for each study participants, along with the squared difference for the data-and-model derived joint peak measurement for each model. The joint measurement for the peak is defined as $\frac{\text{peak magnitude} - \text{mean peak magnitude}}{\text{mean peak magnitude}} + \frac{\text{peak timing} - \text{mean peak timing}}{\text{mean peak timing}}$.

| ID | Data peak joint | Standard peak joint | Error Standard | DDDI peak joint | Error DDDI | MOI peak joint | Error MOI | DDDDI & MOI peak joint | Error DDDDI & MOI | Best Model |
|---|---|---|---|---|---|---|---|---|---|---|
| 1 | 2.03 | 1.94 | 0.009 | 2.24 | 0.042 | 2.05 | 2E-04 | 2.05 | 3E-04 | MOI |
| 2 | 1.46 | 1.43 | 0.0011 | 1.55 | 0.0076 | 1.59 | 0.0149 | 1.62 | 0.0239 | Standard |
| 4 | 2.48 | 2.35 | 0.0178 | 2.27 | 0.0435 | 2.36 | 0.0157 | 2.36 | 0.015 | DDDDI & MOI |
| 5 | 1.88 | 1.26 | 0.3833 | 1.11 | 0.5877 | 1.28 | 0.3616 | 1.86 | 4E-04 | DDDDI & MOI |
| 6 | 2.03 | 1.91 | 0.0134 | 1.88 | 0.0212 | 1.9 | 0.0155 | 1.86 | 0.0278 | Standard |
| 7 | 1.78 | 1.88 | 0.0089 | 1.85 | 0.0042 | 1.92 | 0.0201 | 1.92 | 0.0184 | DDDI |
| 8 | 1.9 | 2.04 | 0.0198 | 2.08 | 0.035 | 1.98 | 0.0064 | 2.05 | 0.0233 | MOI |
| 11 | 2.33 | 2.45 | 0.0149 | 2.34 | 2E-04 | 2.3 | 0.0012 | 2.35 | 3E-04 | DDDI |
| 12 | 2.28 | 2.33 | 0.0023 | 2.28 | 0 | 2.13 | 0.0227 | 2.14 | 0.0206 | DDDI |
| 20 | 2.08 | 1.7 | 0.1481 | 1.84 | 0.0578 | 1.76 | 0.1076 | 1.81 | 0.0758 | DDDI |
| 21 | 2.24 | 2.34 | 0.0094 | 2.2 | 0.0016 | 2.22 | 4E-04 | 2.2 | 0.002 | MOI |
| 22 | 1.61 | 1.51 | 0.0107 | 1.68 | 0.0048 | 1.59 | 3E-04 | 1.64 | 0.001 | MOI |
| 23 | 1.86 | 1.66 | 0.0382 | 1.84 | 4E-04 | 1.83 | 6E-04 | 1.9 | 0.0023 | DDDI |
| 24 | 2.76 | 3.69 | 0.8746 | 3.1 | 0.1153 | 2.84 | 0.0069 | 2.72 | 0.0012 | DDDDI & MOI |
| 25 | 1.74 | 2 | 0.0657 | 1.85 | 0.012 | 1.93 | 0.0332 | 1.93 | 0.0353 | DDDI |
| 26 | 1.98 | 1.97 | 8.46E-6 | 2.03 | 0.0028 | 2.06 | 0.0075 | 2.04 | 0.0046 | Standard |
| 27 | 2.39 | 2.46 | 0.0038 | 2.37 | 7E-04 | 2.33 | 0.0044 | 2.27 | 0.0147 | DDDI |
| 28 | 2.11 | 2.09 | 3E-04 | 2.09 | 3E-04 | 2.06 | 0.0023 | 2.04 | 0.0044 | DDDI |
| 29 | 2.12 | 1.8 | 0.1046 | 1.92 | 0.0409 | 1.87 | 0.0606 | 1.87 | 0.0616 | DDDI |

| | | | | | | | | | | |
|---|---|---|---|---|---|---|---|---|---|---|
| 31 | 1.6 | 1.47 | 0.0171 | 1.45 | 0.0203 | 1.37 | 0.0535 | 1.46 | 0.0174 | Standard |
| 32 | 1.62 | 1.28 | 0.1138 | 1.58 | 0.0019 | 1.58 | 0.0017 | 1.62 | 1.31E-5 | DDDDI & MOI |
| 33 | 1.51 | 1.61 | 0.0112 | 1.53 | 7E-04 | 1.63 | 0.0165 | 1.64 | 0.0194 | DDDI |
| 34 | 1.91 | 2.18 | 0.0715 | 2.01 | 0.0101 | 2.04 | 0.017 | 1.99 | 0.0068 | DDDDI & MOI |
| 37 | 2.25 | 1.78 | 0.2222 | 2.06 | 0.0341 | 2.09 | 0.026 | 1.27 | 0.9604 | MOI |
| 40 | 1.9 | 1.89 | 1E-04 | 1.72 | 0.0307 | 1.83 | 0.0047 | 1.82 | 0.0065 | Standard |
| 41 | 2.19 | 2.26 | 0.004 | 2.2 | 3.83E-5 | 2.15 | 0.0016 | 2.14 | 0.0024 | DDDI |
| 42 | 2.1 | 2.23 | 0.0168 | 2.1 | 4.11E-6 | 2.22 | 0.0127 | 2.22 | 0.0126 | DDDI |
| 44 | 1.95 | 1.82 | 0.0165 | 2 | 0.0022 | 2 | 0.0026 | 1.94 | 2E-04 | DDDDI & MOI |
| 46 | 2.3 | 2.02 | 0.0796 | 2.21 | 0.0076 | 2.16 | 0.0197 | 2.16 | 0.0194 | DDDI |
| 48 | 2.06 | 2.01 | 0.0026 | 1.95 | 0.0116 | 2 | 0.004 | 1.94 | 0.015 | Standard |
| 49 | 1.93 | 2.11 | 0.0326 | 1.96 | 0.0014 | 2.07 | 0.0214 | 2.14 | 0.0442 | DDDI |
| 52 | 1.91 | 1.9 | 2E-04 | 1.84 | 0.0055 | 1.91 | 2.04E-6 | 2.49 | 0.3339 | MOI |
| 55 | 1.8 | 1.95 | 0.0216 | 1.98 | 0.0314 | 1.92 | 0.015 | 1.9 | 0.0106 | DDDDI & MOI |
| 57 | 1.89 | 1.95 | 0.0046 | 2.15 | 0.0706 | 2.05 | 0.0274 | 2.1 | 0.0448 | Standard |
| 58 | 2.01 | 2.07 | 0.0038 | 2.03 | 5E-04 | 2.1 | 0.0092 | 2.11 | 0.0115 | DDDI |
| 59 | 2.03 | 2.09 | 0.0047 | 2.09 | 0.0043 | 2.01 | 1E-04 | 2.06 | 0.0013 | MOI |
| 61 | 2.38 | 2.17 | 0.0432 | 2.23 | 0.0236 | 2.19 | 0.0387 | 2.19 | 0.0356 | DDDI |
| 62 | 1.85 | 2.05 | 0.0417 | 2.11 | 0.0689 | 2.06 | 0.0451 | 2.1 | 0.0631 | Standard |
| 64 | 2.43 | 2.39 | 0.0014 | 2.52 | 0.009 | 2.48 | 0.0024 | 2.52 | 0.0092 | Standard |
| 65 | 1.64 | 1.83 | 0.0375 | 1.53 | 0.0118 | 1.97 | 0.1136 | 1.35 | 0.0813 | DDDI |
| 67 | 1.74 | 1.91 | 0.0294 | 1.95 | 0.0469 | 1.9 | 0.0255 | 1.93 | 0.0371 | MOI |
| 71 | 1.97 | 2.13 | 0.0265 | 2.16 | 0.0369 | 2.08 | 0.0134 | 2.13 | 0.025 | MOI |
| 73 | 1.99 | 2.11 | 0.0137 | 2.09 | 0.0097 | 2.19 | 0.0404 | 2.14 | 0.0211 | DDDI |

**Derivation of simplified terms in the *Density-Dependent Cell Death & MOI* model**

We again assume that the death rate of infected cells $\alpha_i$ scales linearly with MOI $i$. Therefore,

$$H\left(\sum_{i=0}^{\infty}\alpha_i p_i\right)^{\gamma} = H\left(\sum_{i=0}^{\infty} i\alpha p_i\right)^{\gamma}$$

$$= H\alpha^{\gamma}\left(\sum_{i=0}^{\infty} i p_i\right)^{\gamma}$$

$$= H\alpha^{\gamma}\left(\frac{P}{H}\right)^{\gamma}$$

$$= \alpha^{\gamma}\frac{P^{\gamma}}{H^{\gamma-1}}$$

and

$$H\left(\sum_{i=0}^{\infty} i\ \alpha_i p_i\right)^{\gamma} = H\left(\sum_{i=0}^{\infty} i^2 \alpha p_i\right)^{\gamma}$$

$$= H\alpha^{\gamma}\left(\sum_{i=0}^{\infty} i^2 p_i\right)^{\gamma}$$

$$= H\alpha^{\gamma}\left[\frac{P}{H} + \left(\frac{P}{H}\right)^2 + \frac{1+k}{k}\right]^{\gamma}$$

$$= \alpha^{\gamma}\frac{P^{\gamma}}{H^{\gamma-1}}\left[1 + \frac{P}{H}\left(\frac{1+k}{k}\right)\right]^{\gamma}$$

**Model comparison Figures**

In the figures below we provide the fitted curves of the four models (solid lines) to the viral load measurements (points) for each of the 43 study participants. The black line is for the Standard model, the green line for the Density-dependent cell death model, the pink line for the MOI model and orange line for the Density-dependent cell death & MOI model.

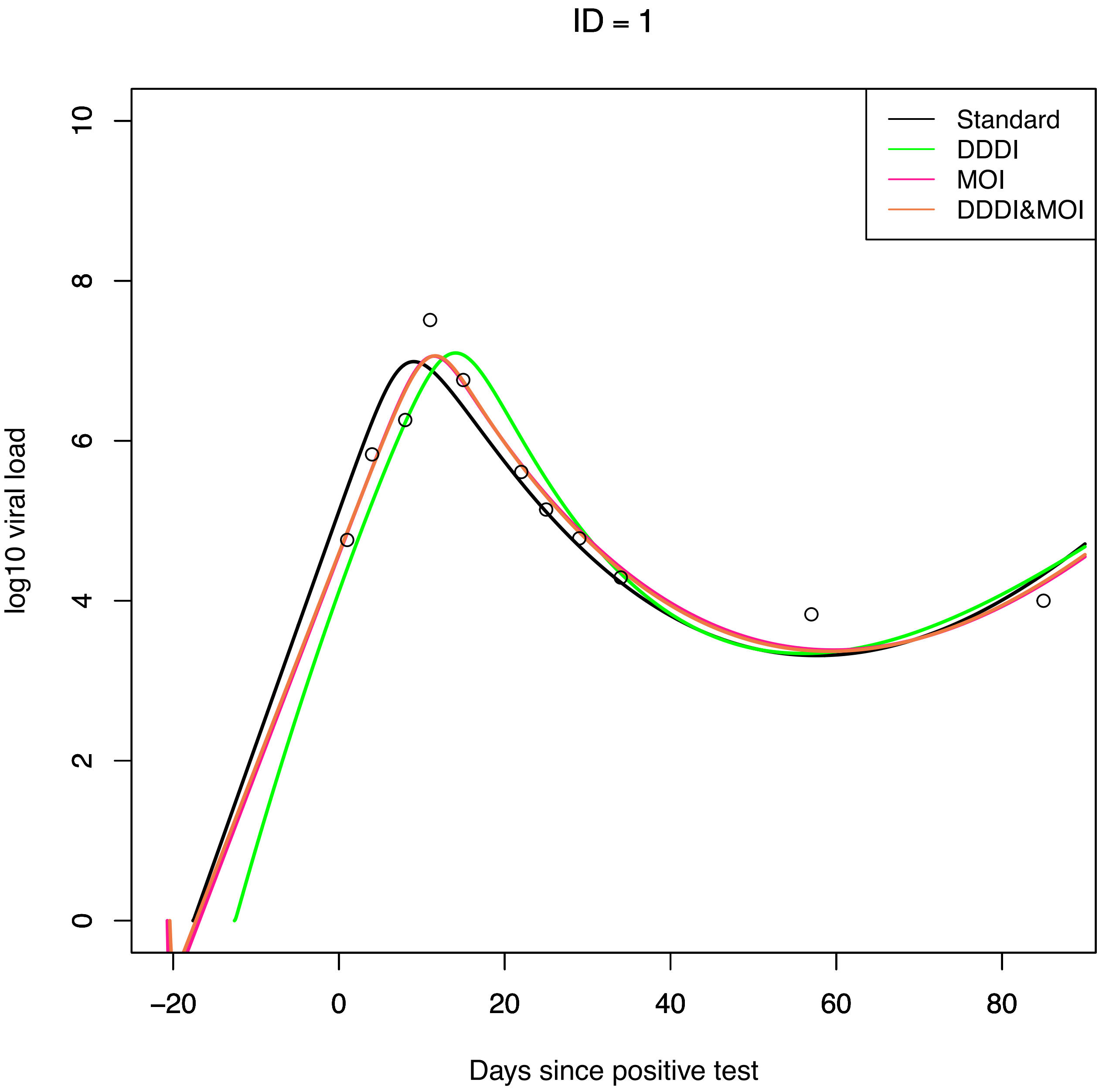
ID = 1
Standard
DDDI
MOI
DDDI&MOI
log10 viral load
0
2
4
6
8
10
−20
0
20
40
60
80
Days since positive test

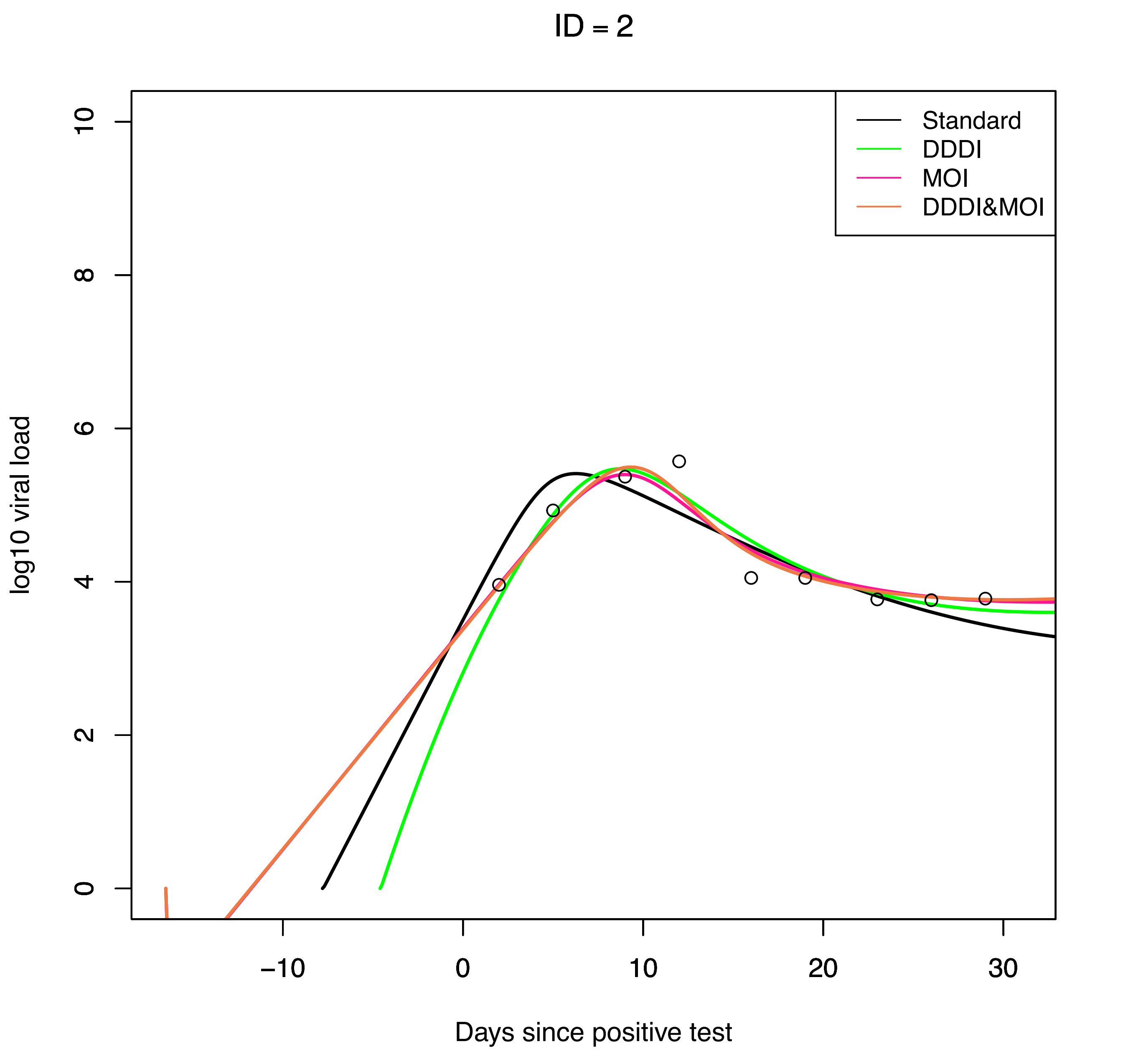

ID = 2
Standard
DDDI
MOI
DDDI&MOI
log10 viral load
Days since positive test
0
2
4
6
8
10
−10
0
10
20
30

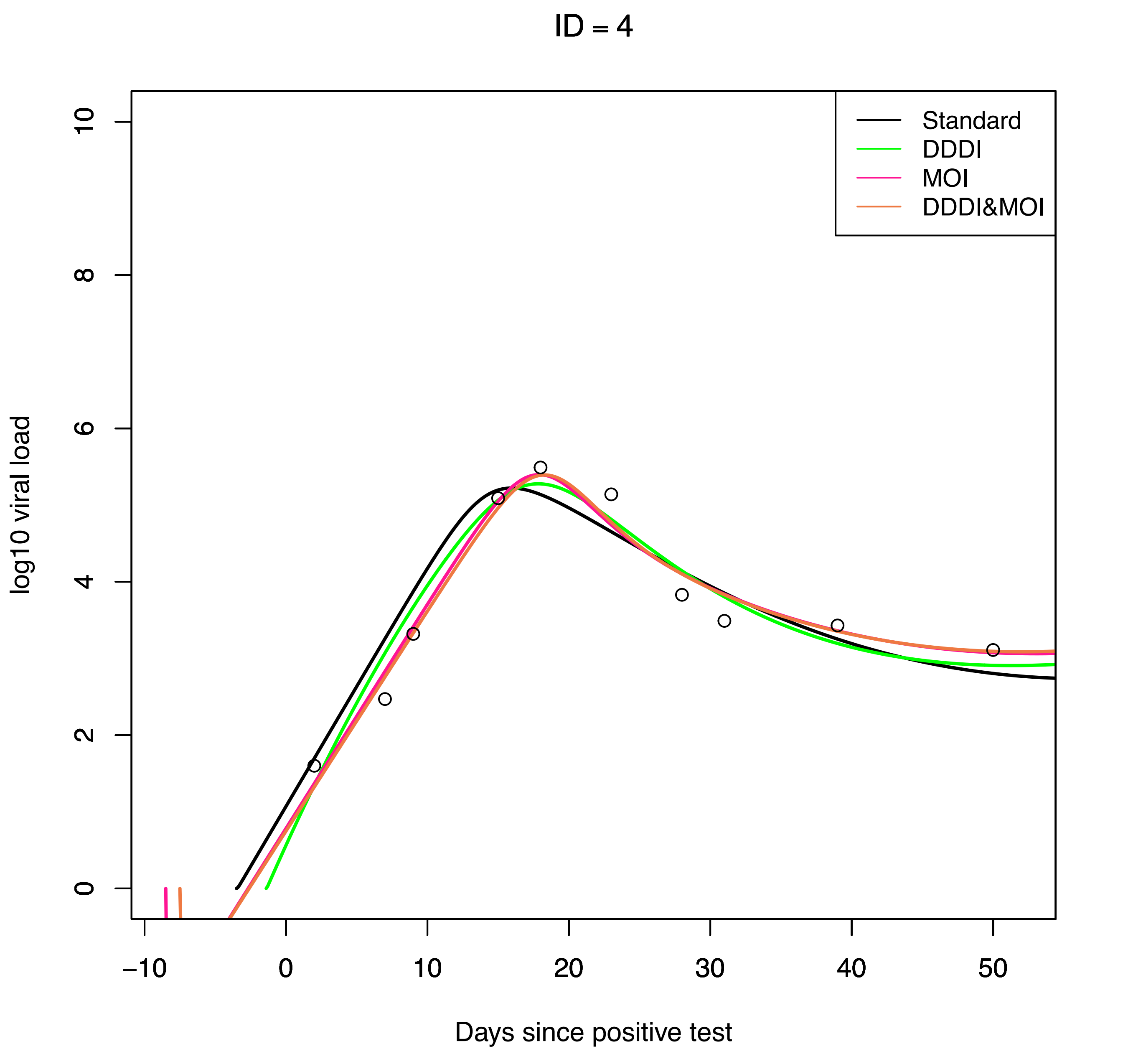
ID = 4
Standard
DDDI
MOI
DDDI&MOI
log10 viral load
Days since positive test
0
2
4
6
8
10
−10
0
10
20
30
40
50

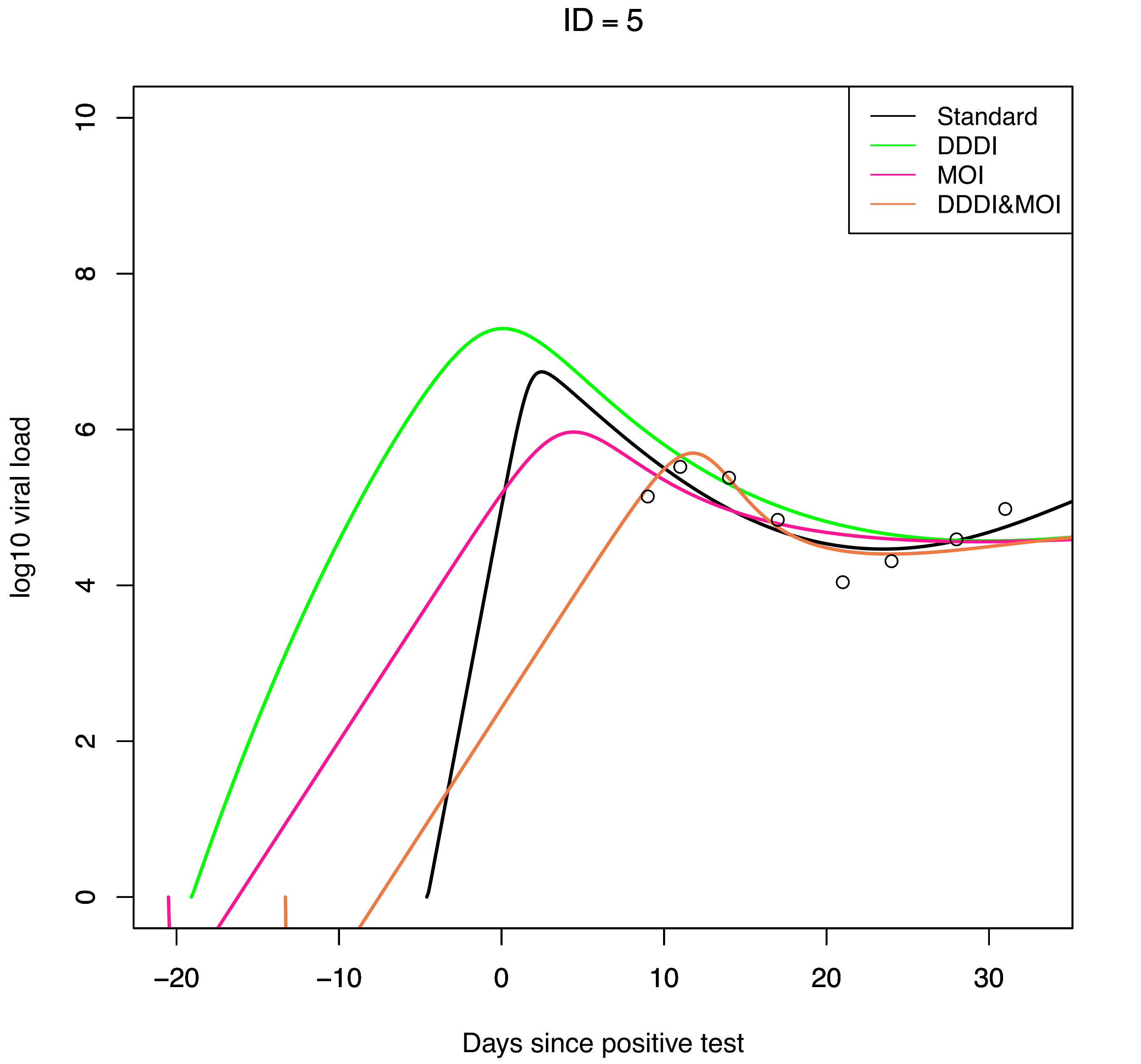
ID = 5
Standard
DDDI
MOI
DDDI&MOI
log10 viral load
Days since positive test
0
2
4
6
8
10
−20
−10
0
10
20
30

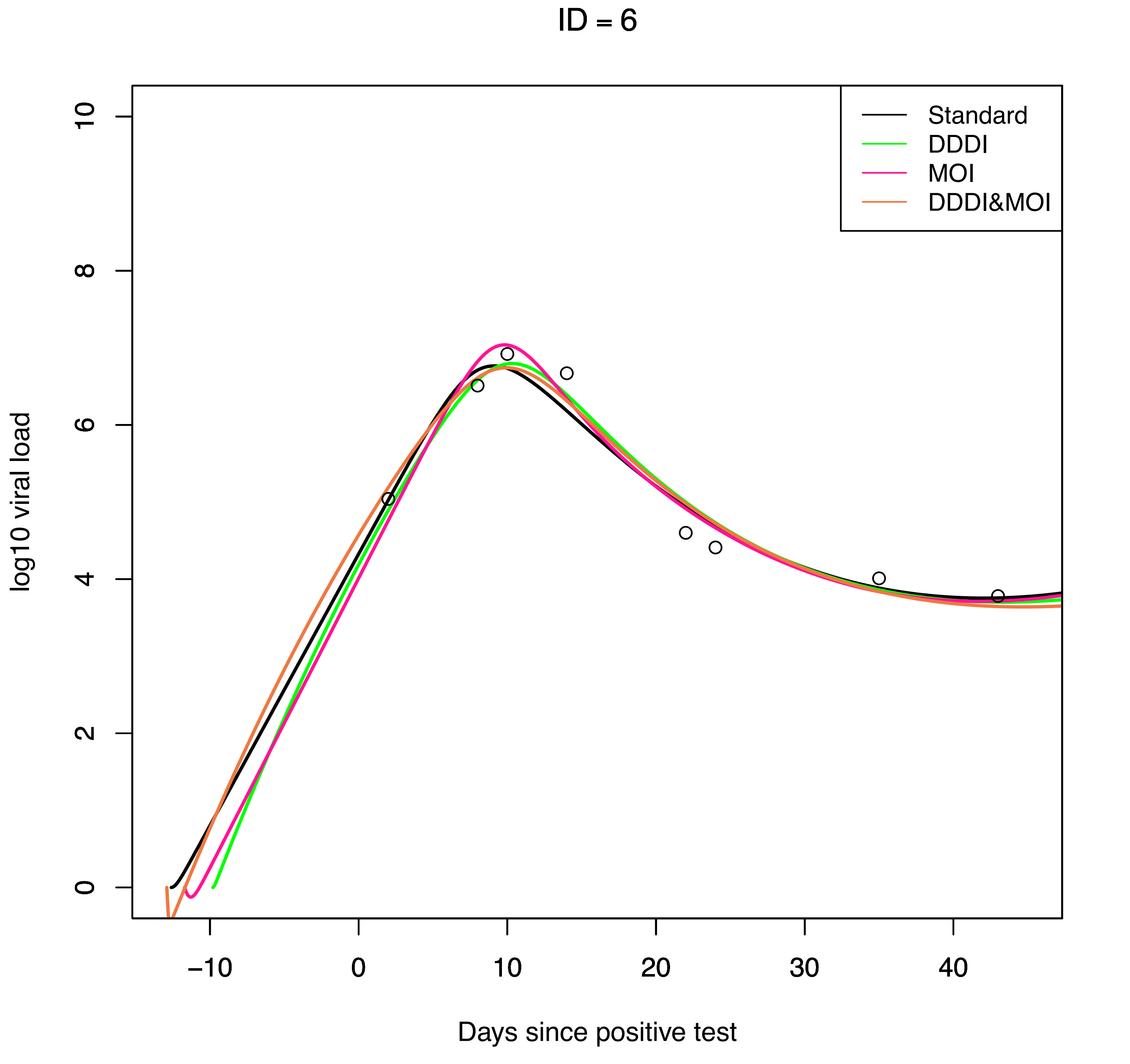

ID = 6
Standard
DDDI
MOI
DDDI&MOI
log10 viral load
Days since positive test
0
2
4
6
8
10
−10
0
10
20
30
40

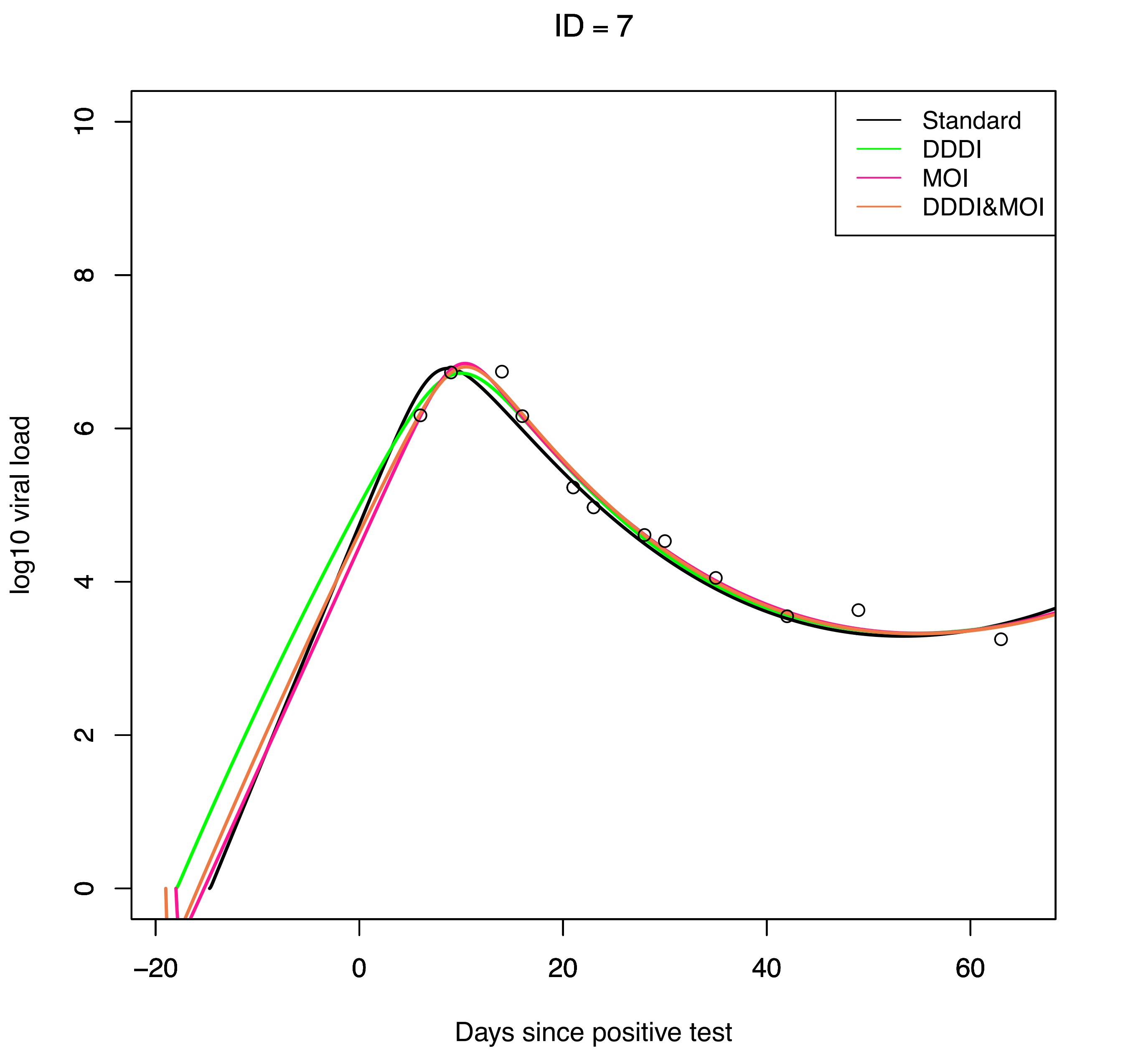
ID = 7
Standard
DDDI
MOI
DDDI&MOI
log10 viral load
Days since positive test
0
2
4
6
8
10
−20
0
20
40
60

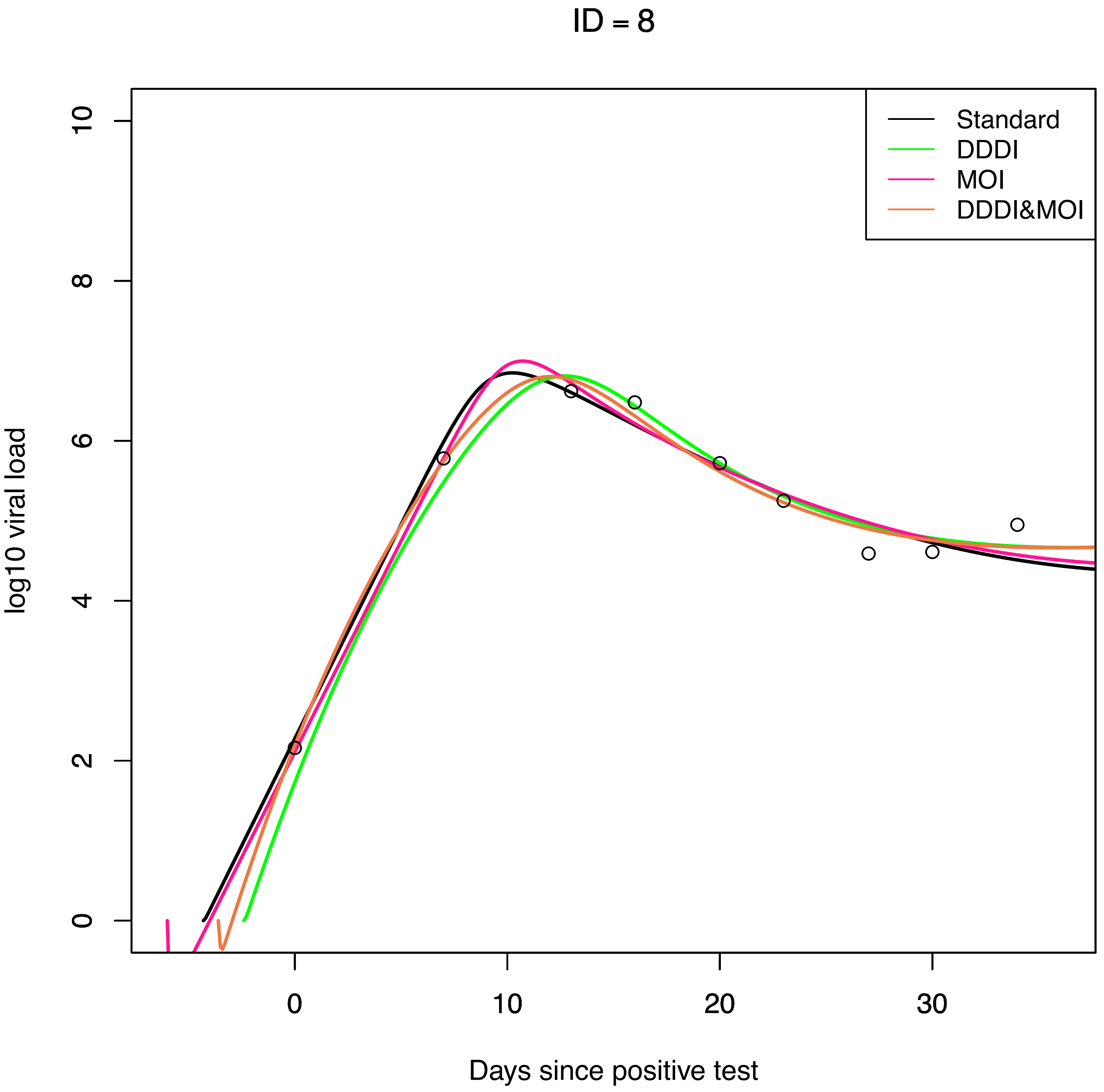
ID = 8
Standard
DDDI
MOI
DDDI&MOI
log10 viral load
0
2
4
6
8
10
0
10
20
30
Days since positive test

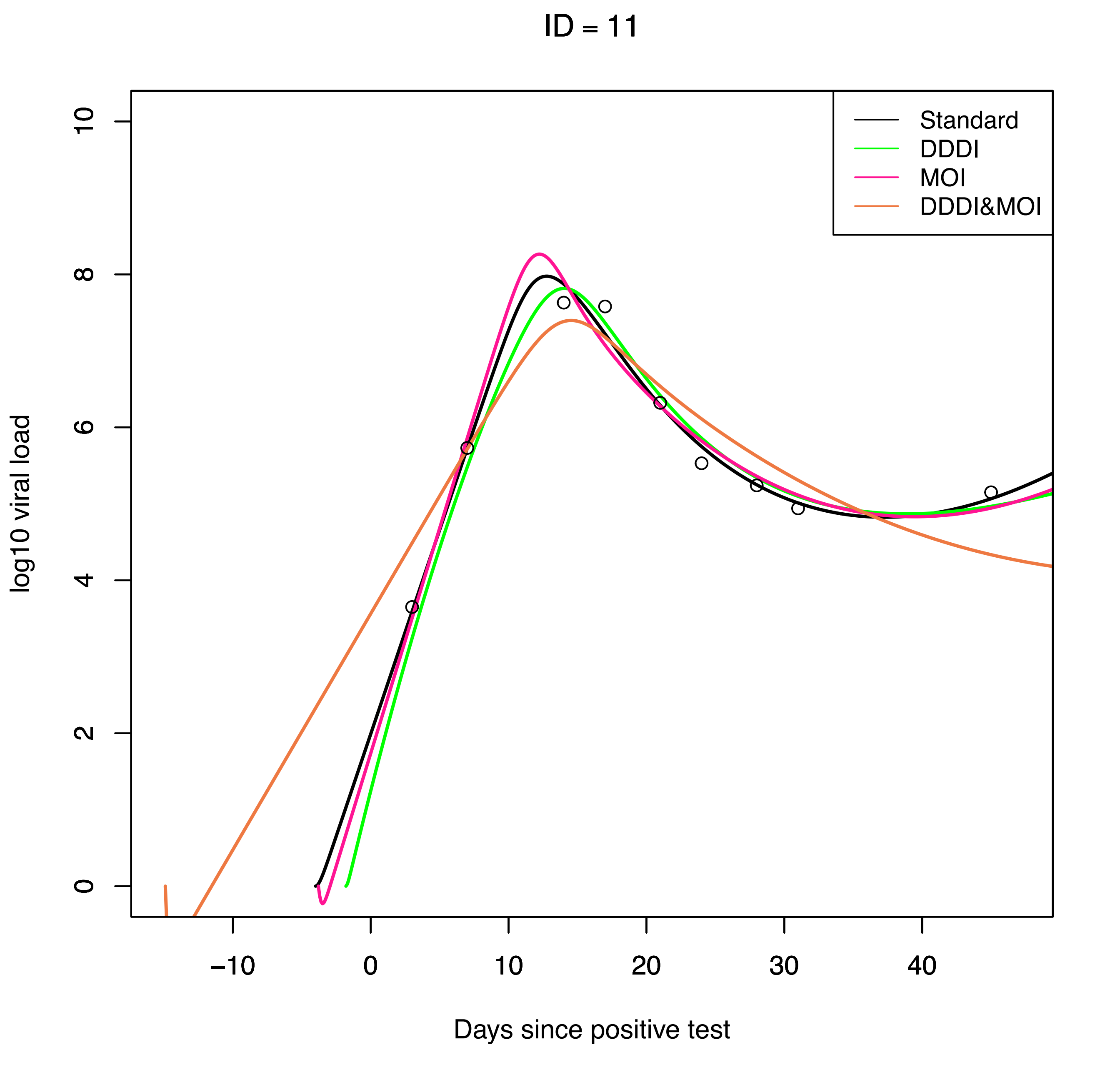
ID = 11
Standard
DDDI
MOI
DDDI&MOI
log10 viral load
0
2
4
6
8
10
−10
0
10
20
30
40
Days since positive test

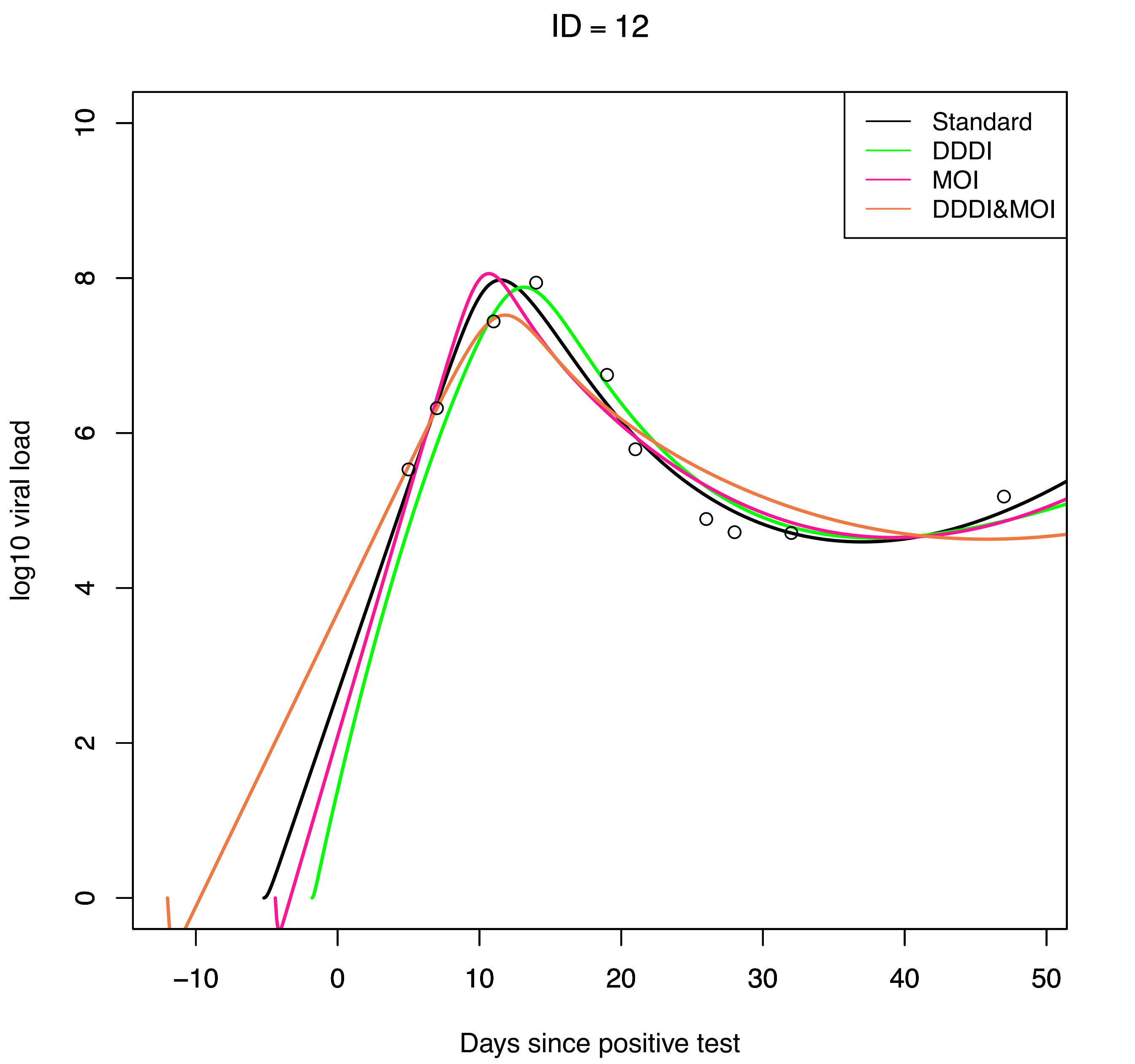
ID = 12
Standard
DDDI
MOI
DDDI&MOI
log10 viral load
0
2
4
6
8
10
−10
0
10
20
30
40
50
Days since positive test

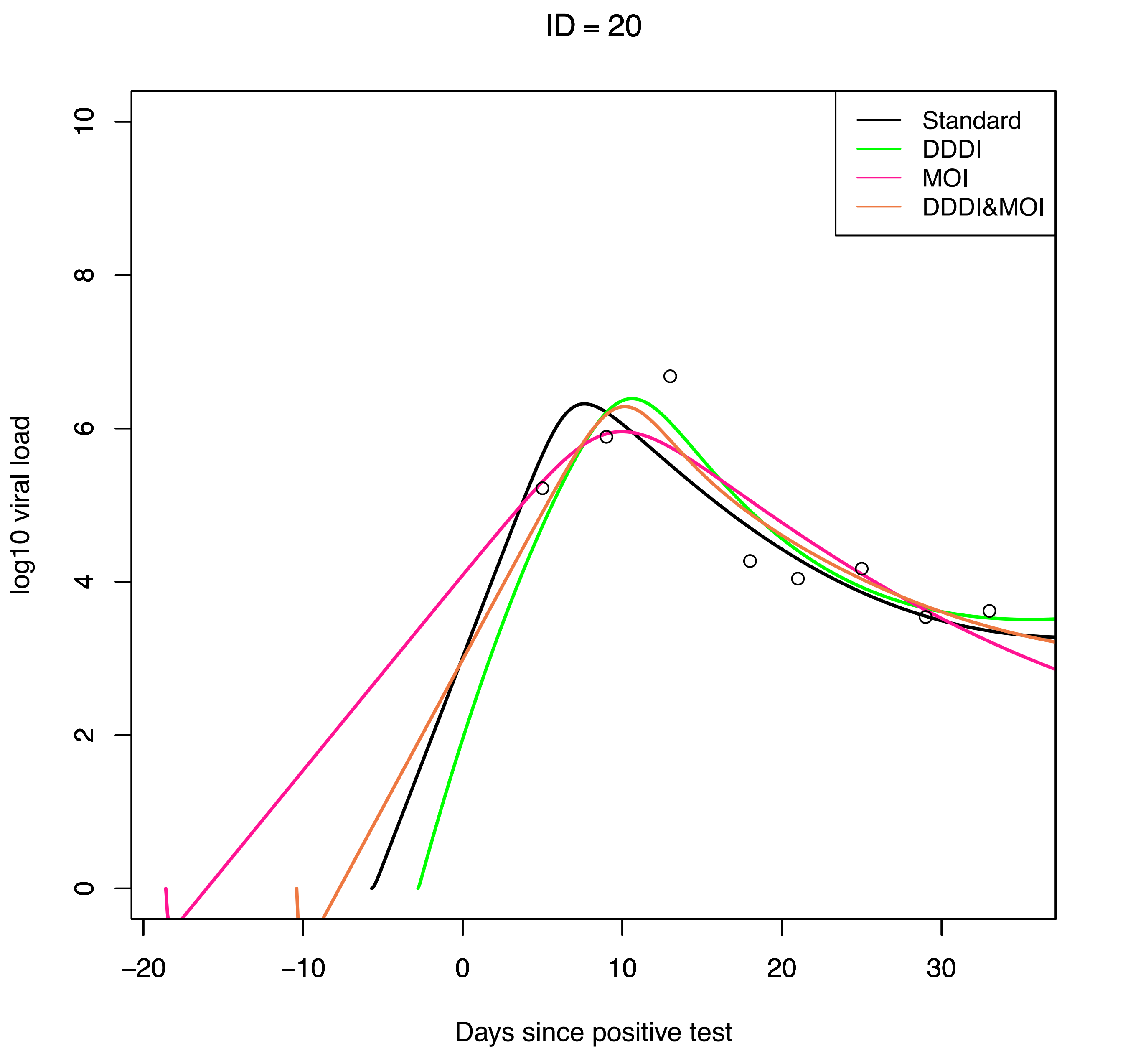

ID = 20
Standard
DDDI
MOI
DDDI&MOI
log10 viral load
0
2
4
6
8
10
−20
−10
0
10
20
30
Days since positive test

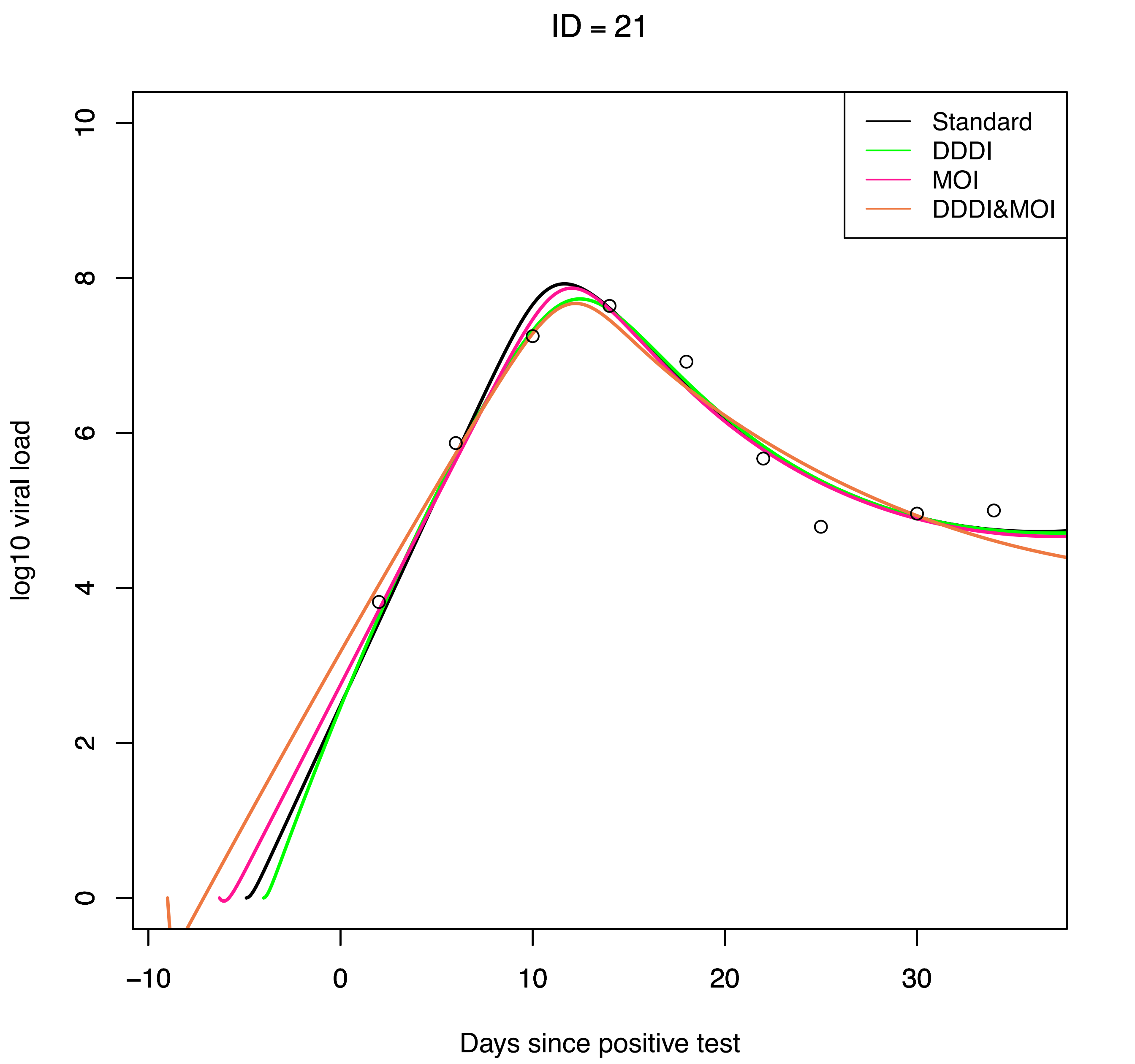
ID = 21
Standard
DDDI
MOI
DDDI&MOI
log10 viral load
0
2
4
6
8
10
−10
0
10
20
30
Days since positive test

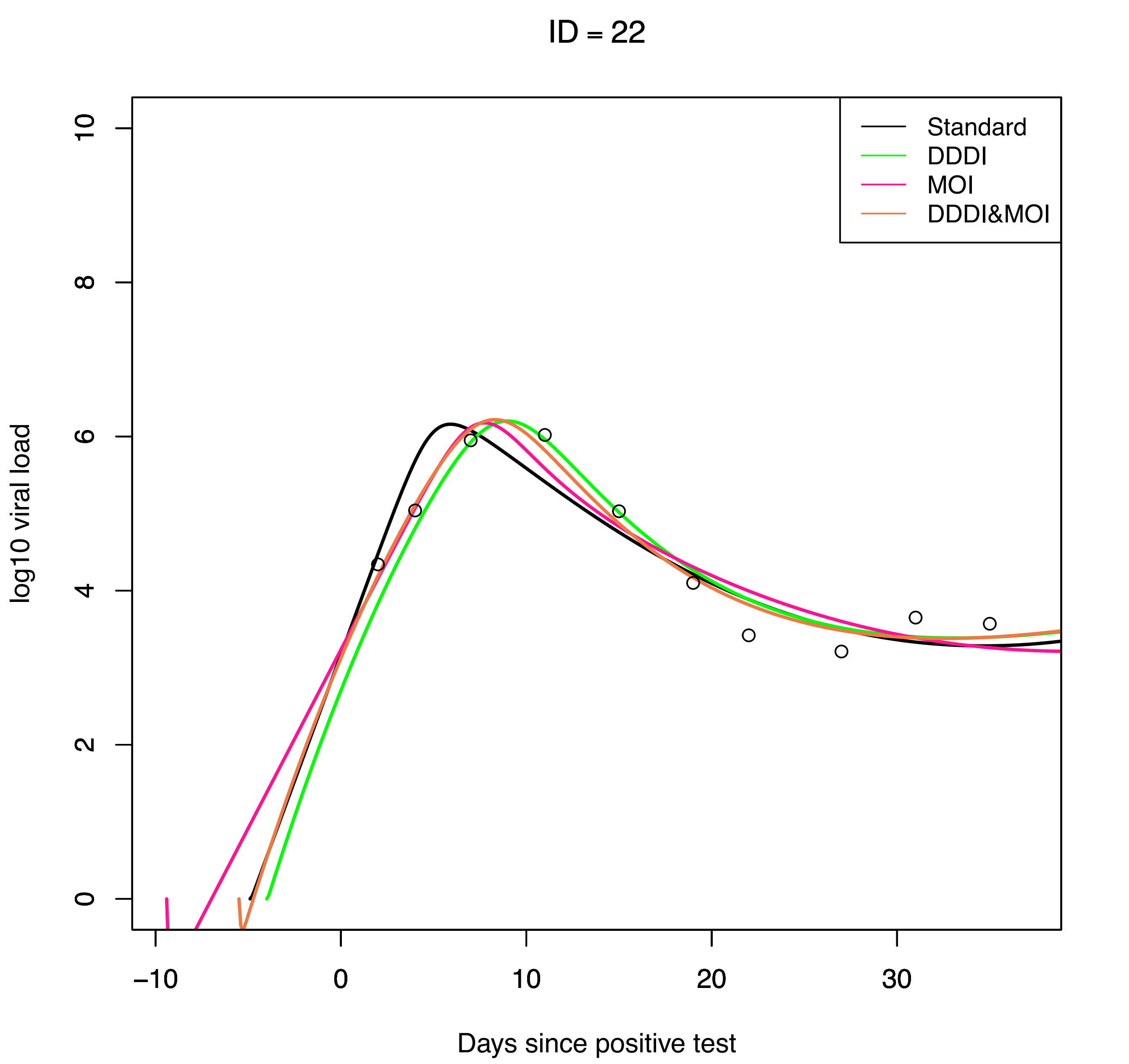

ID = 22
Standard
DDDI
MOI
DDDI&MOI
log10 viral load
0
2
4
6
8
10
−10
0
10
20
30
Days since positive test

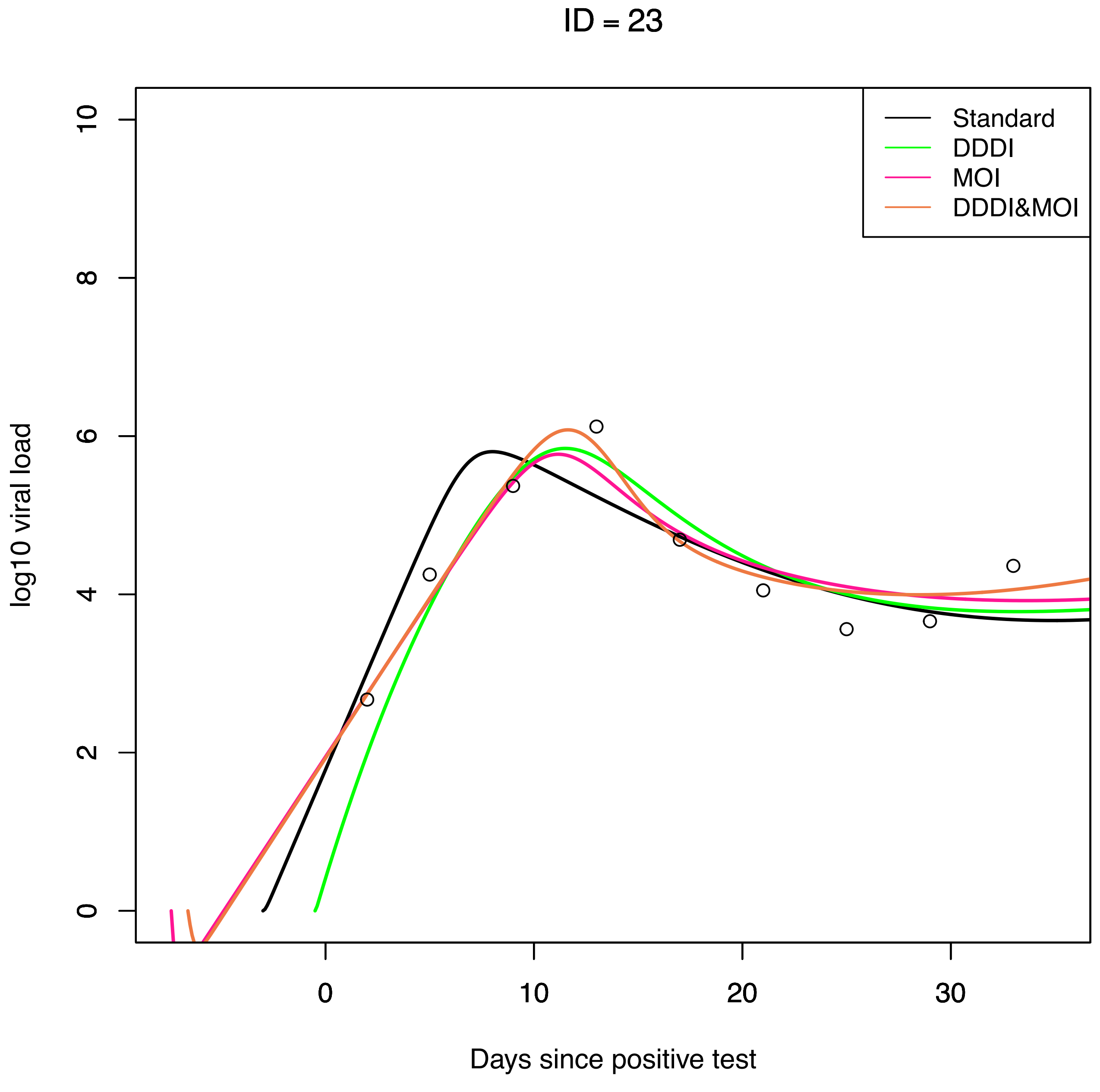
ID = 23
Standard
DDDI
MOI
DDDI&MOI
log10 viral load
0
2
4
6
8
10
0
10
20
30
Days since positive test

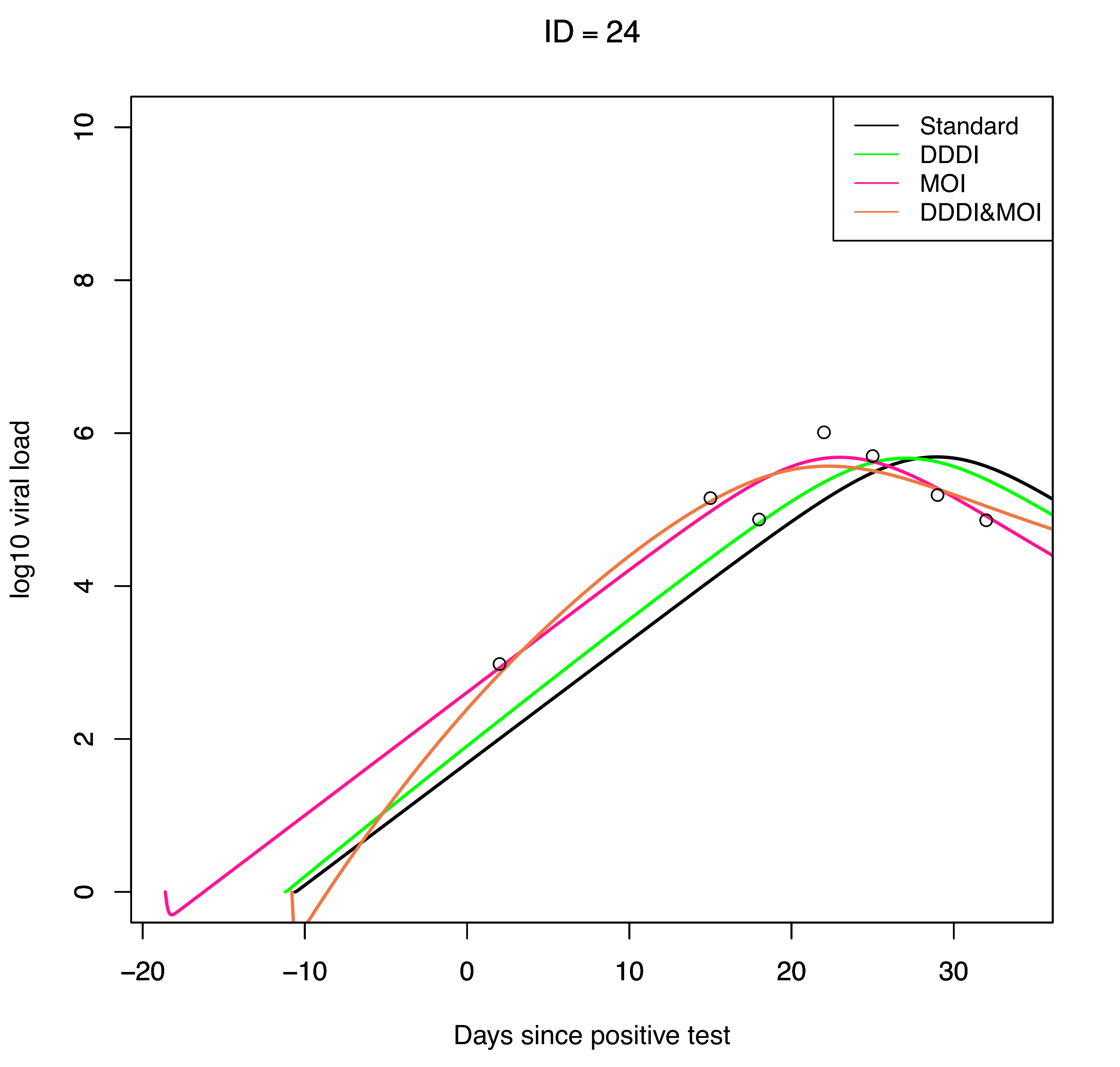

ID = 24
Standard
DDDI
MOI
DDDI&MOI
log10 viral load
0
2
4
6
8
10
−20
−10
0
10
20
30
Days since positive test

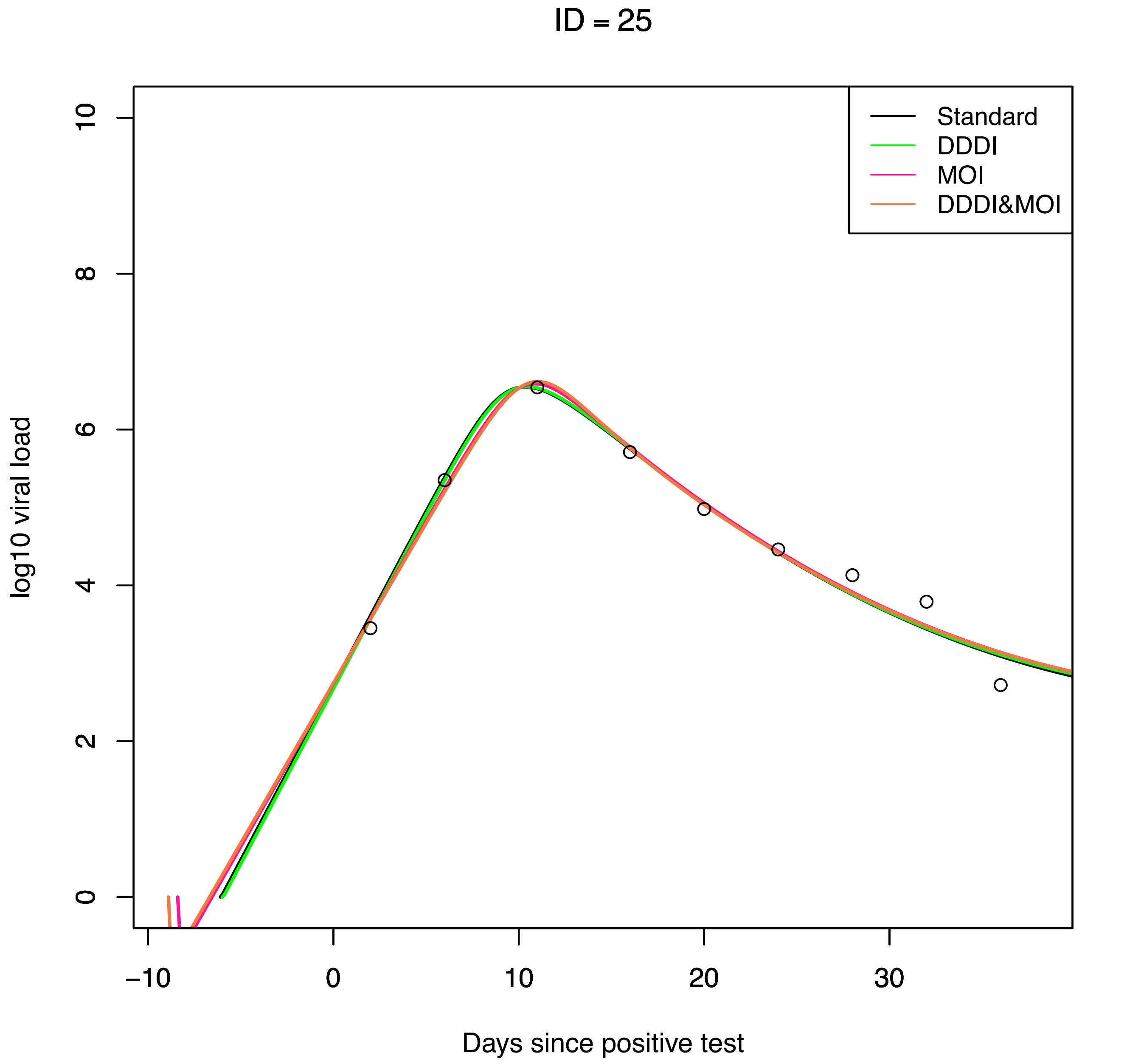
ID = 25
Standard
DDDI
MOI
DDDI&MOI
log10 viral load
0
2
4
6
8
10
−10
0
10
20
30
Days since positive test

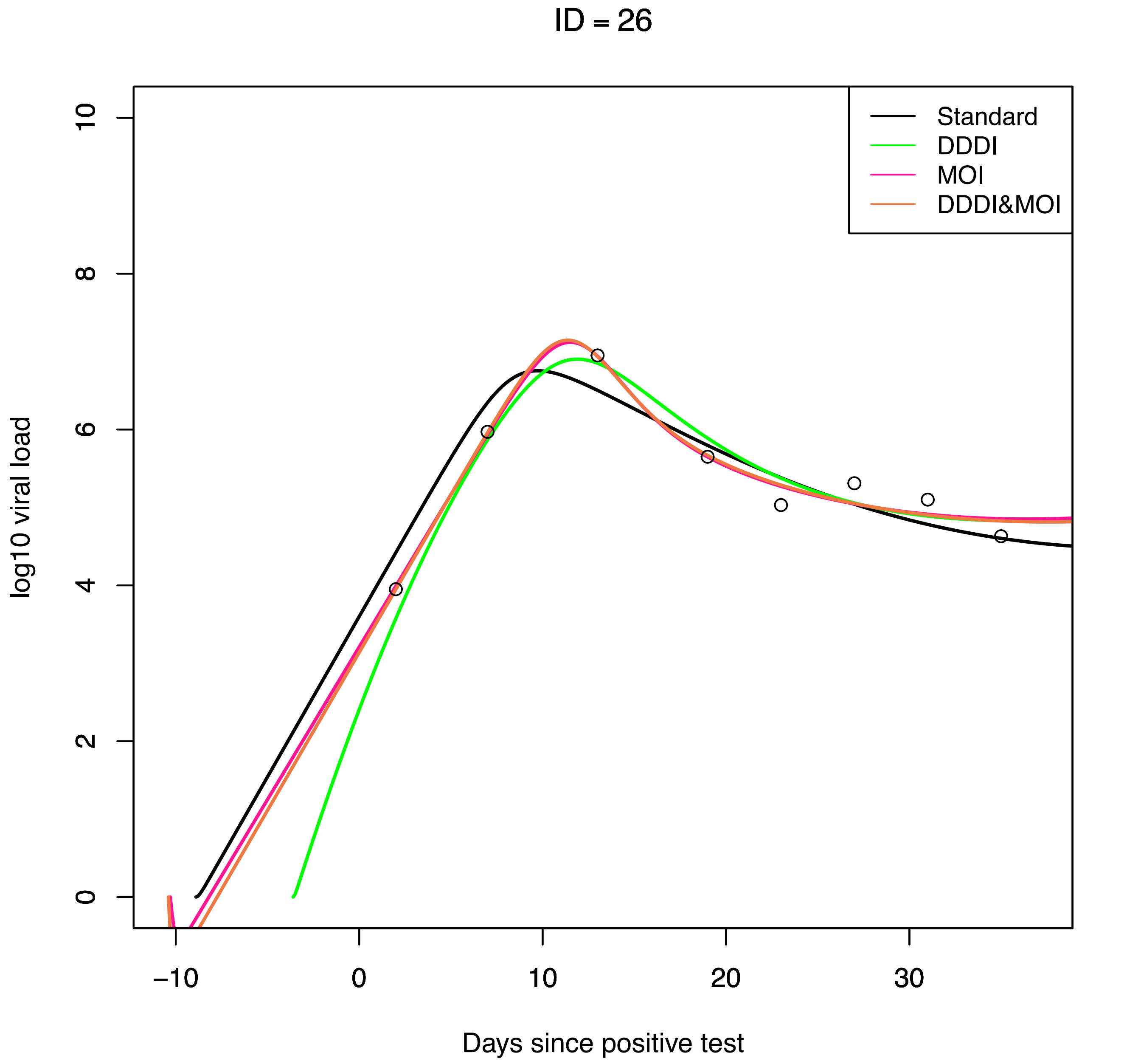

ID = 26
Standard
DDDI
MOI
DDDI&MOI
log10 viral load
0
2
4
6
8
10
−10
0
10
20
30
Days since positive test

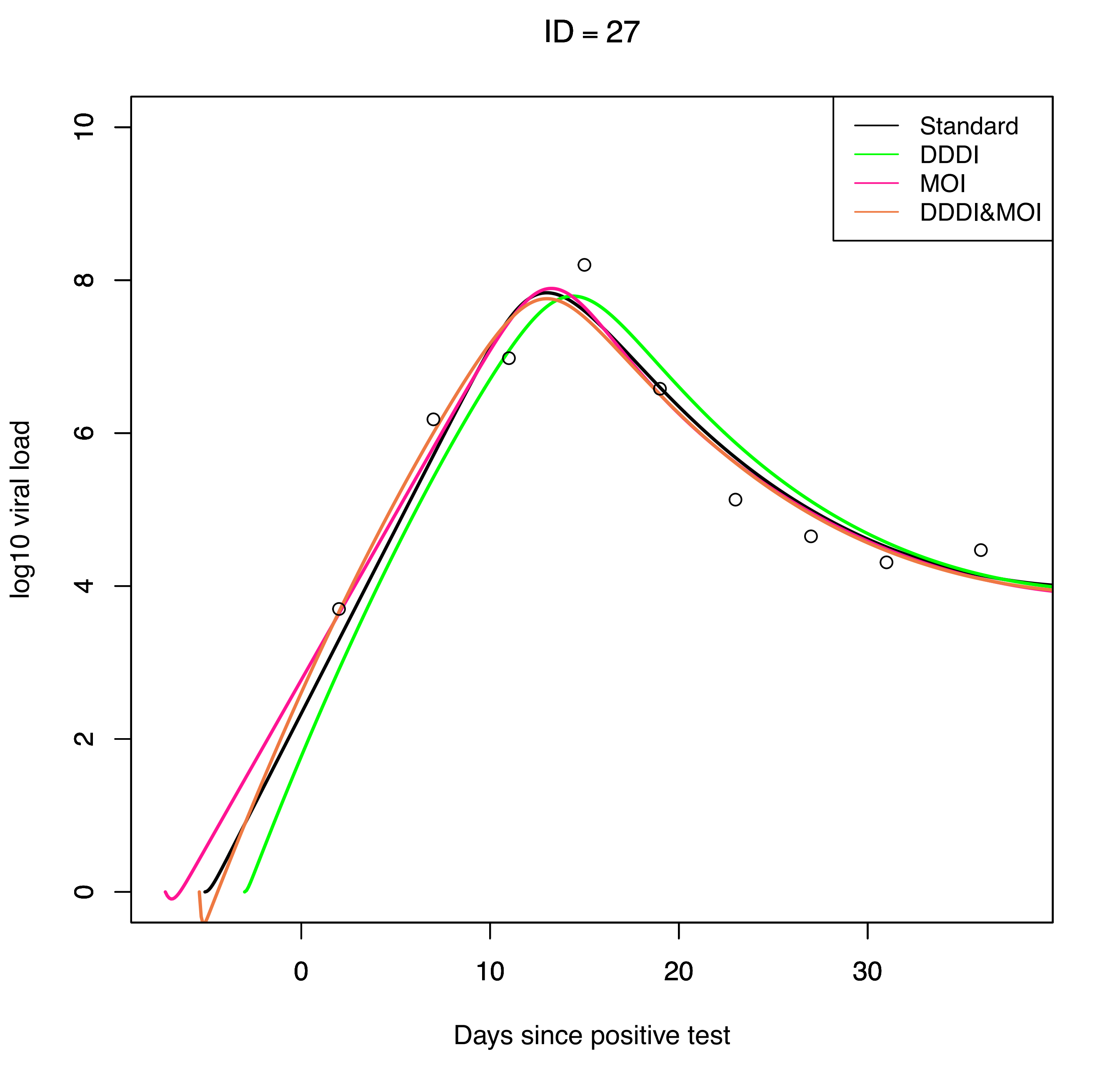
ID = 27
Standard
DDDI
MOI
DDDI&MOI
log10 viral load
0
2
4
6
8
10
0
10
20
30
Days since positive test

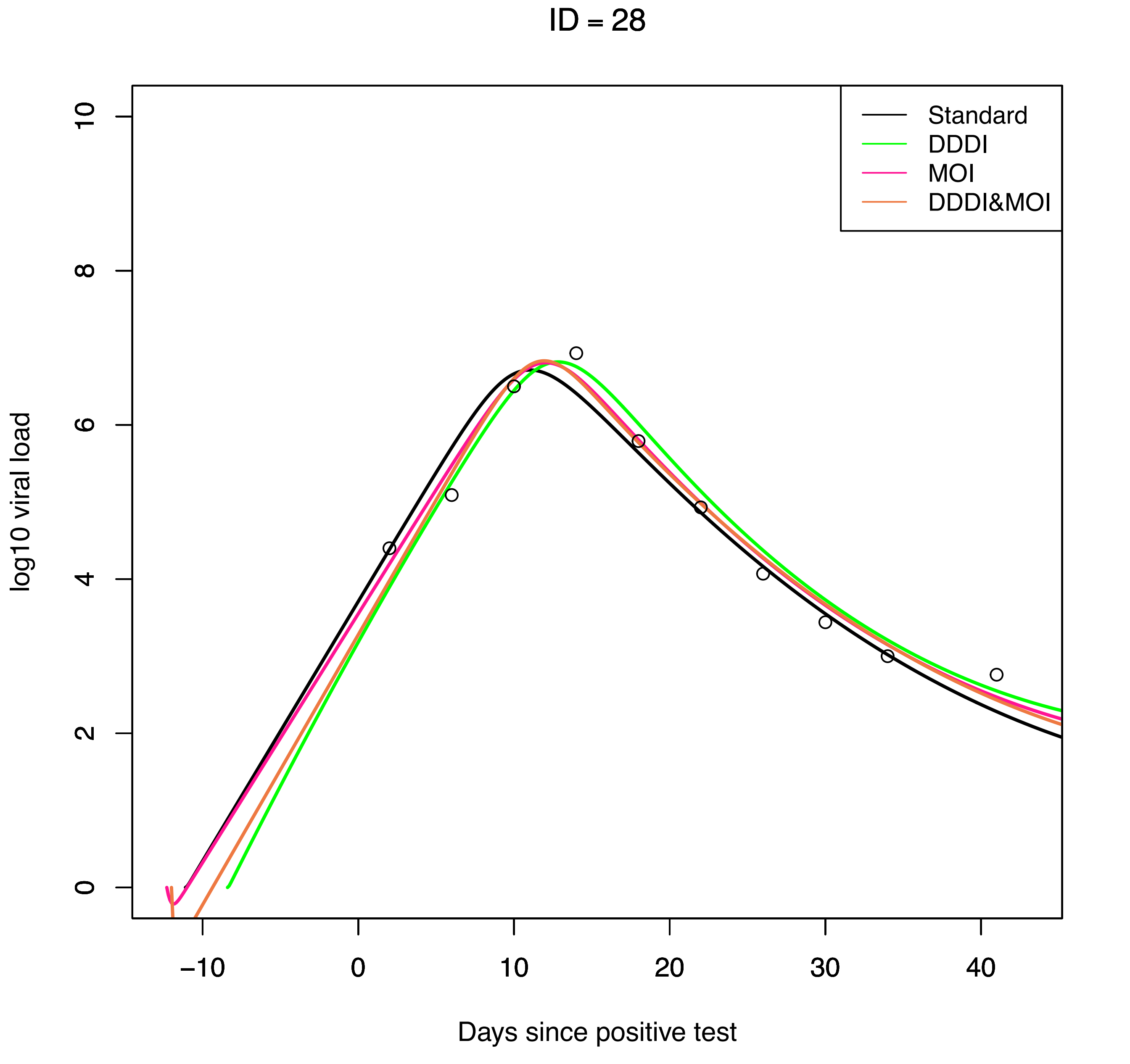
ID = 28
Standard
DDDI
MOI
DDDI&MOI
log10 viral load
0
2
4
6
8
10
−10
0
10
20
30
40
Days since positive test

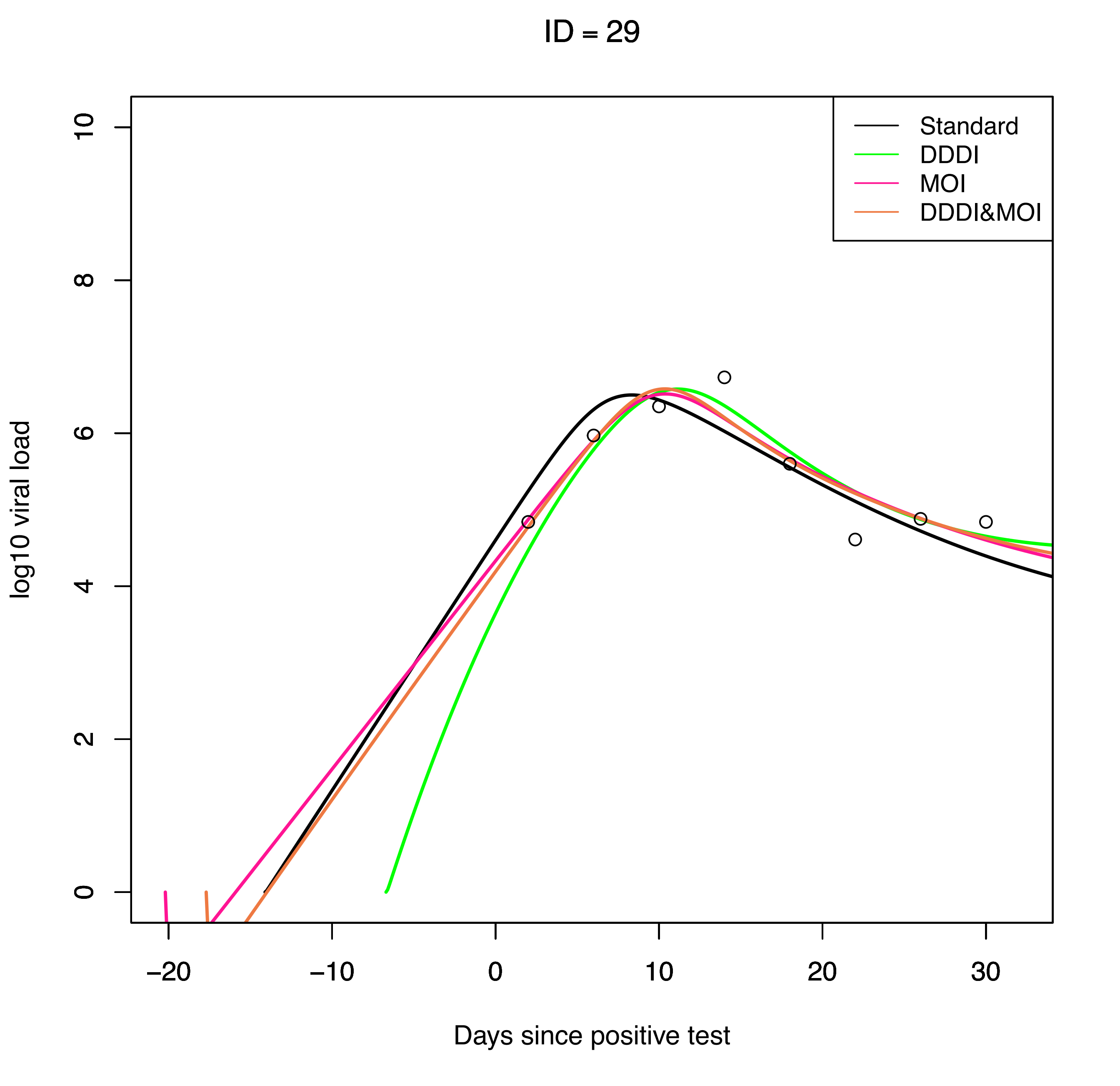
ID = 29
Standard
DDDI
MOI
DDDI&MOI
log10 viral load
0
2
4
6
8
10
−20
−10
0
10
20
30
Days since positive test

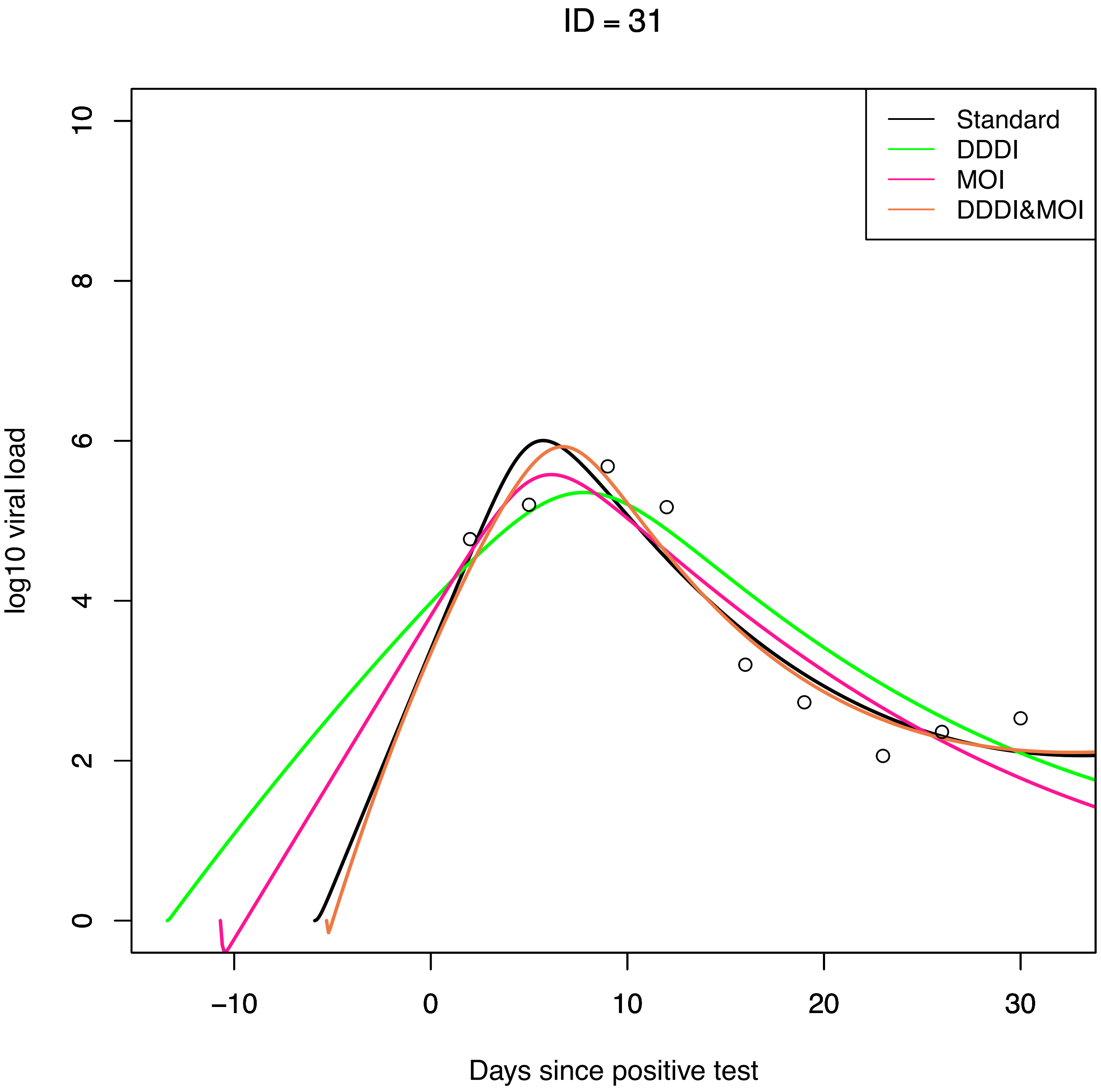
ID = 31
Standard
DDDI
MOI
DDDI&MOI
log10 viral load
0
2
4
6
8
10
−10
0
10
20
30
Days since positive test

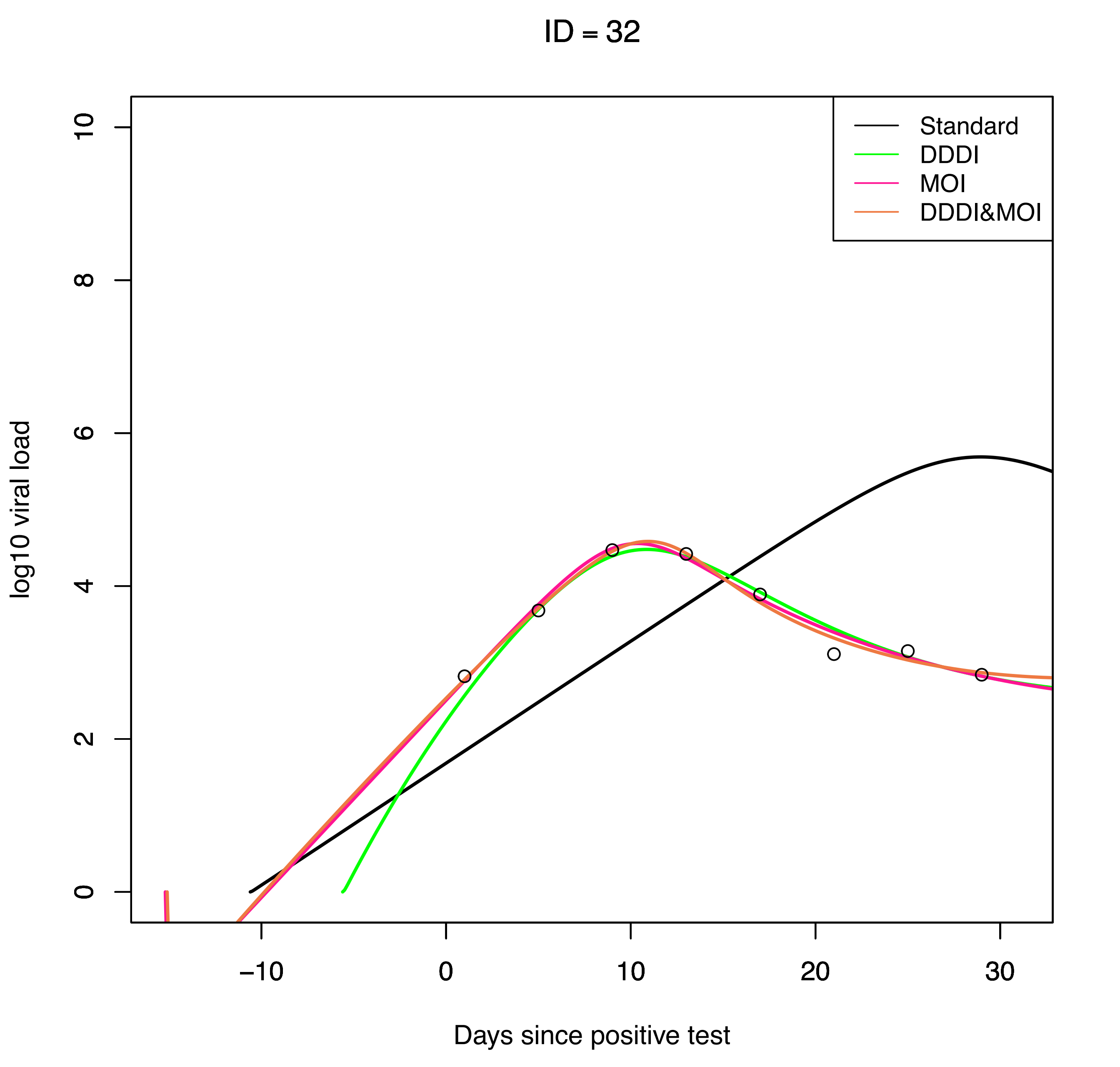
ID = 32
Standard
DDDI
MOI
DDDI&MOI
log10 viral load
0
2
4
6
8
10
−10
0
10
20
30
Days since positive test

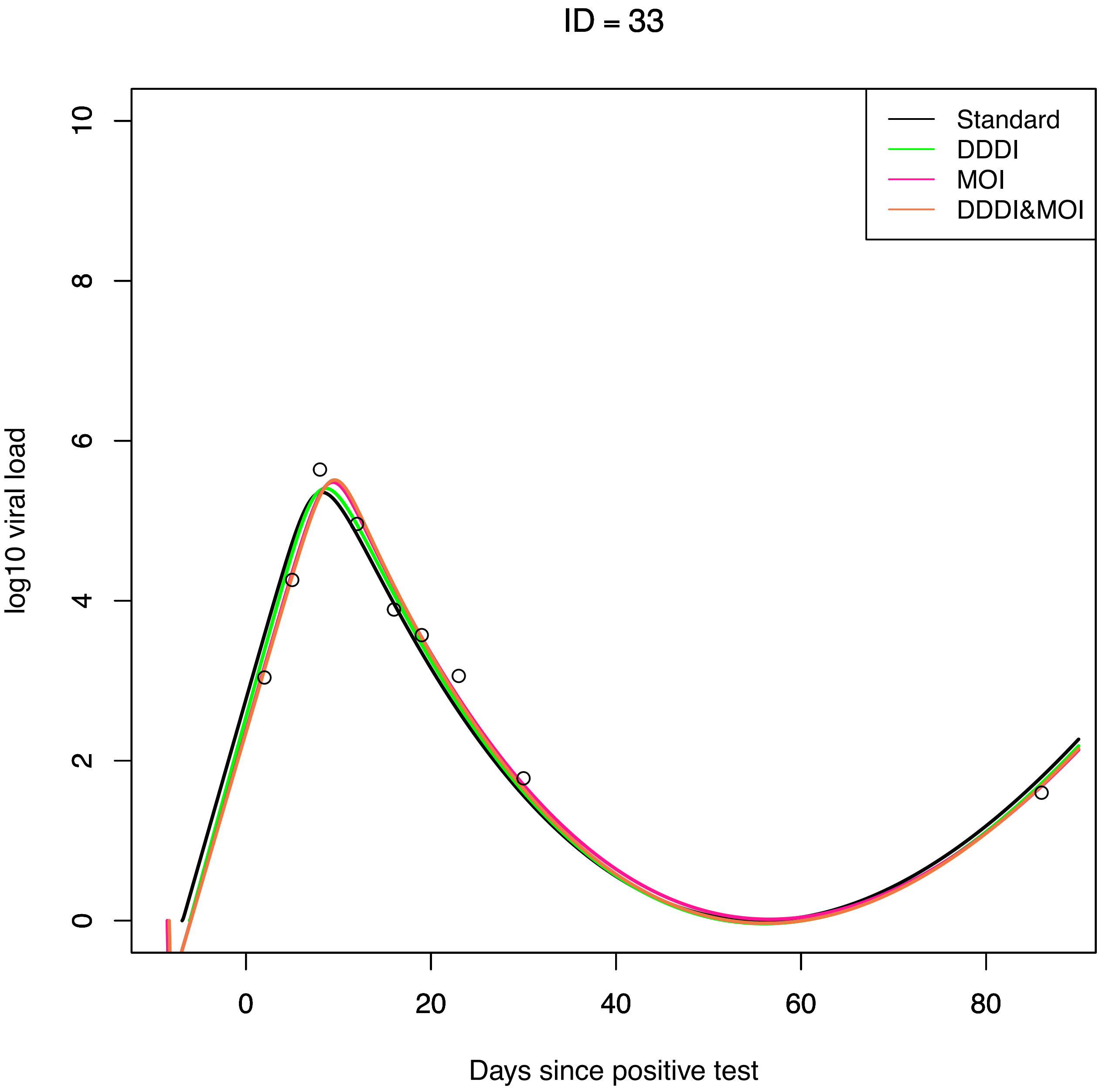
ID = 33
Standard
DDDI
MOI
DDDI&MOI
log10 viral load
0
2
4
6
8
10
0
20
40
60
80
Days since positive test

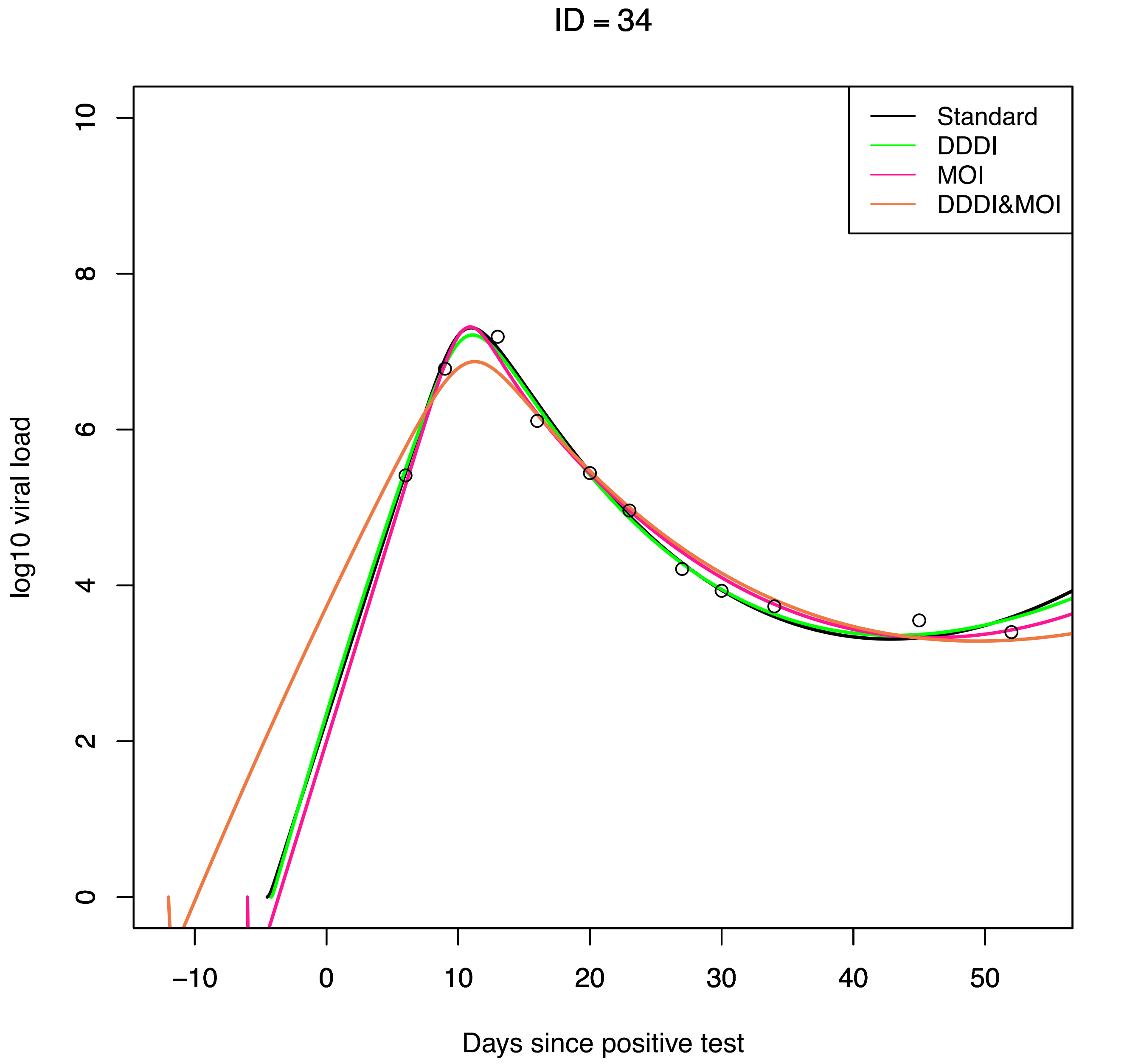

ID = 34
Standard
DDDI
MOI
DDDI&MOI
log10 viral load
Days since positive test
0
2
4
6
8
10
−10
0
10
20
30
40
50

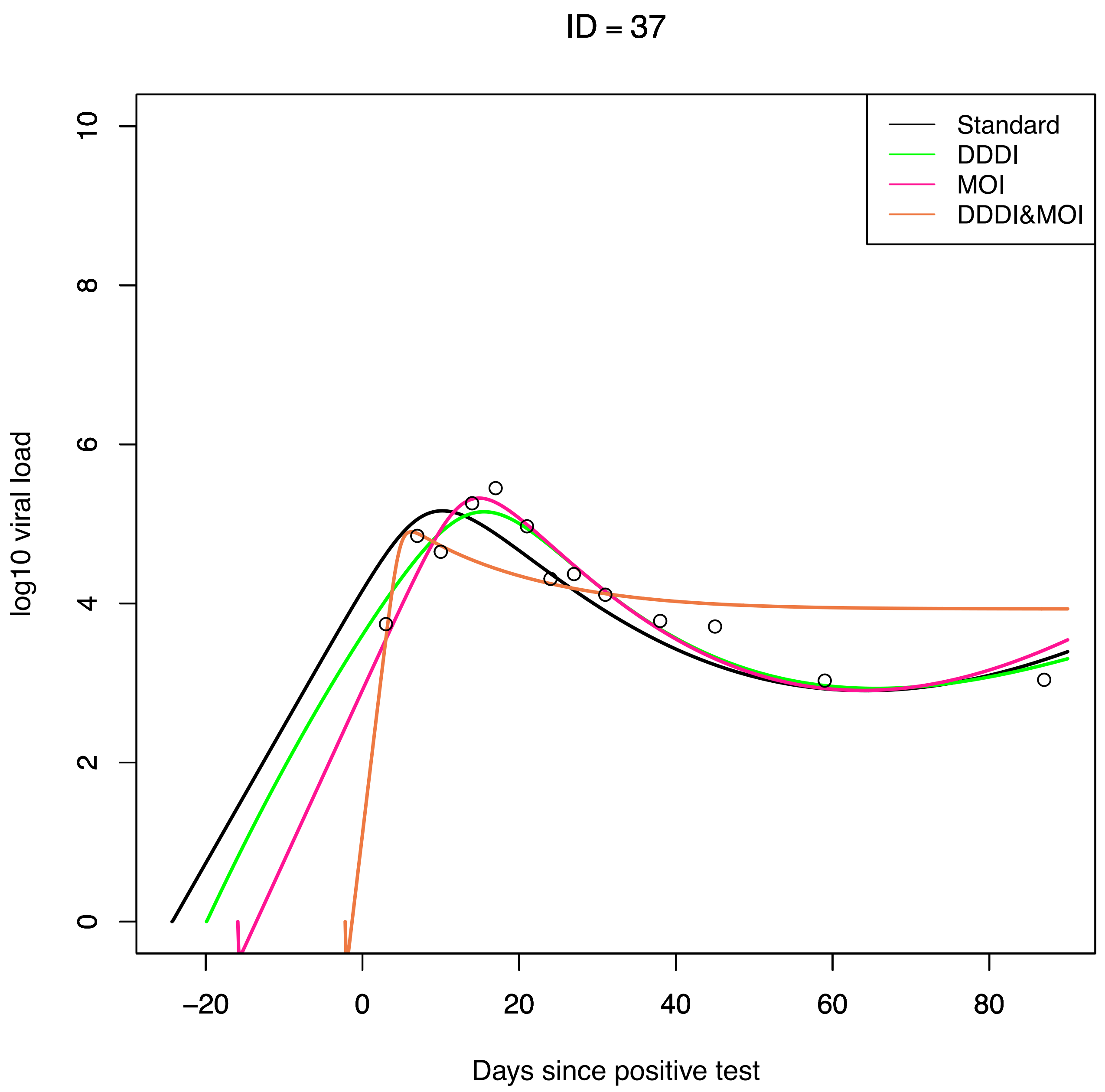

ID = 37
Standard
DDDI
MOI
DDDI&MOI
log10 viral load
0
2
4
6
8
10
−20
0
20
40
60
80
Days since positive test

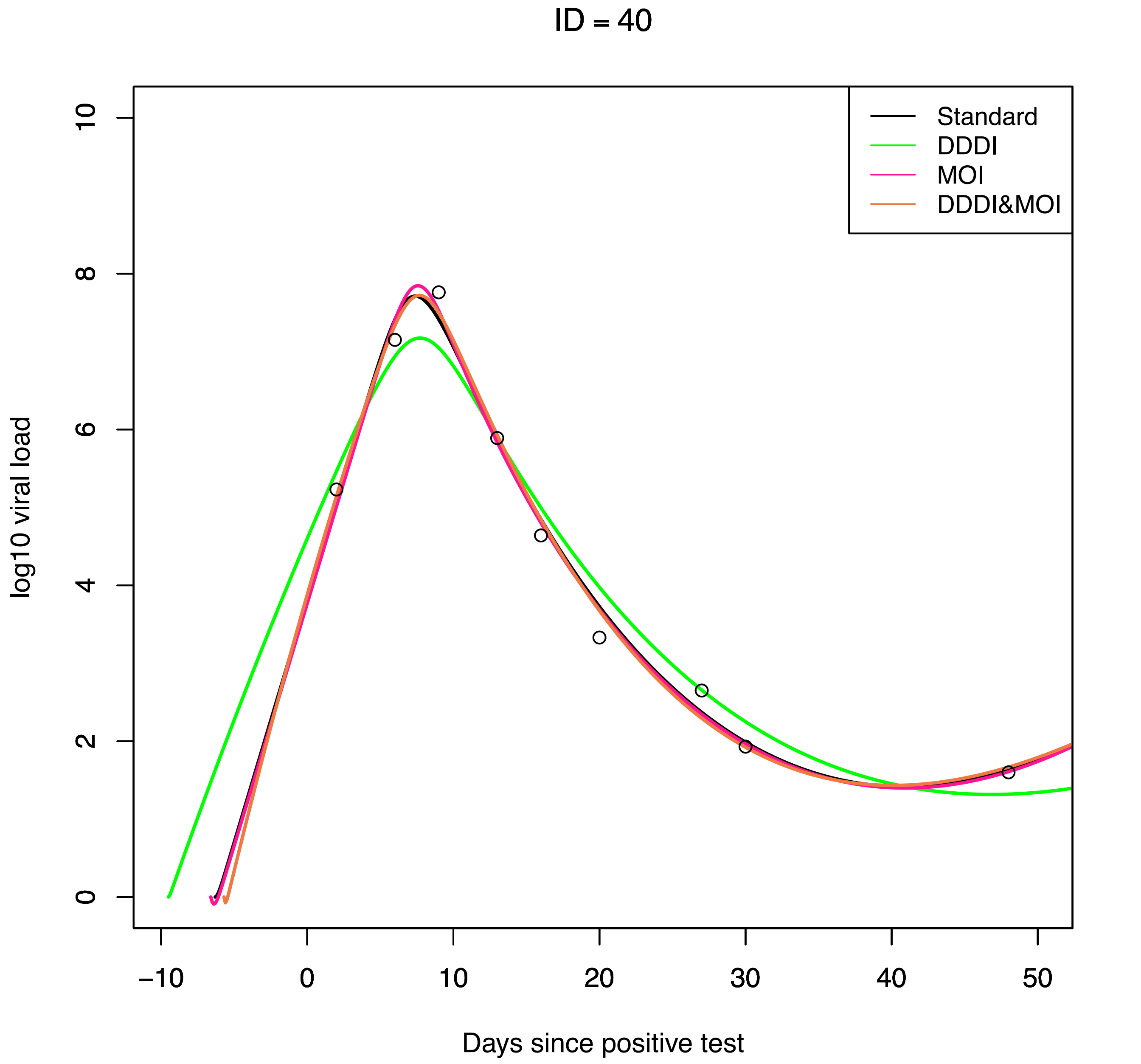
ID = 40
Standard
DDDI
MOI
DDDI&MOI
log10 viral load
Days since positive test
0
2
4
6
8
10
−10
0
10
20
30
40
50

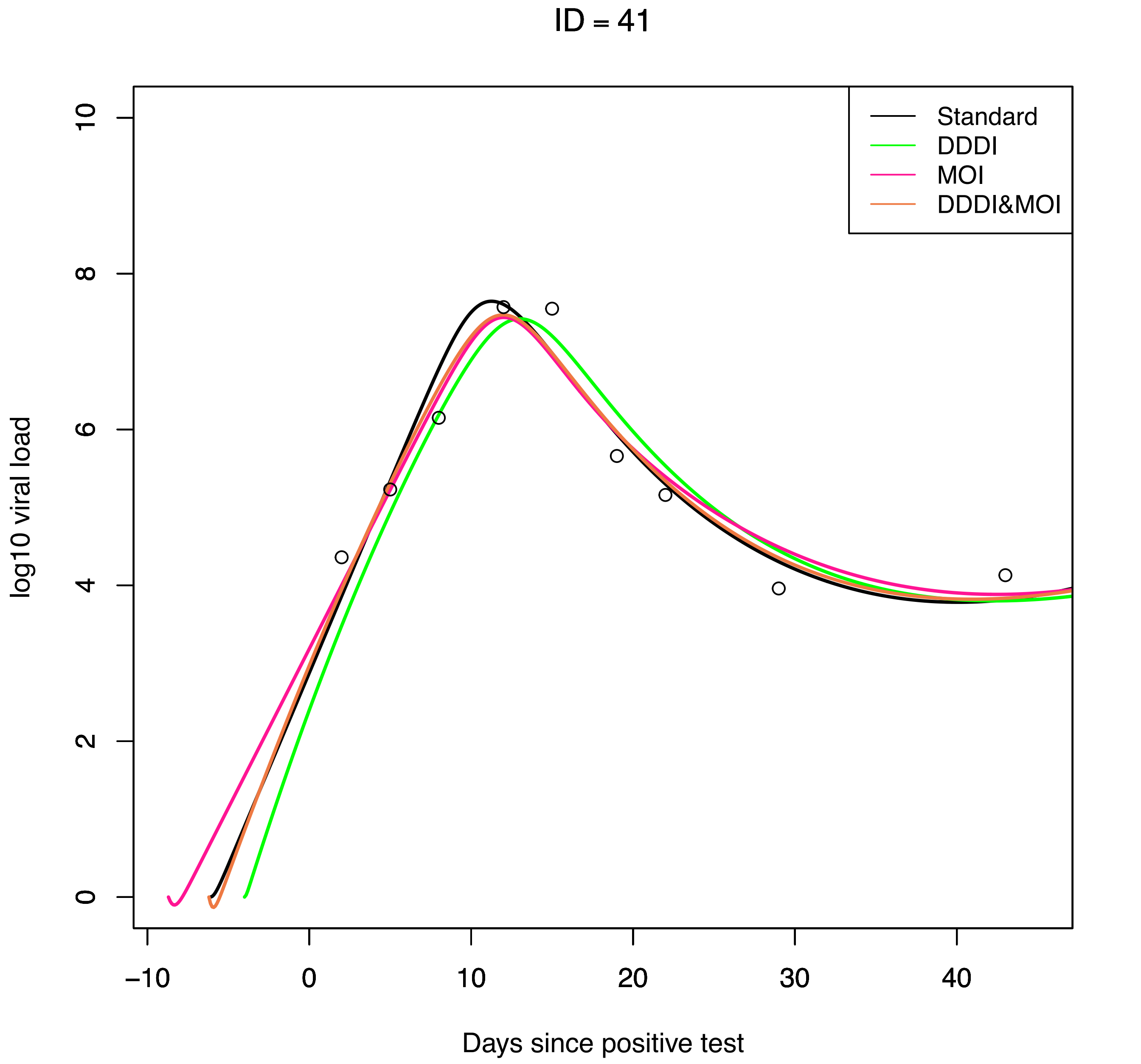

ID = 41
Standard
DDDI
MOI
DDDI&MOI
log10 viral load
Days since positive test
0
2
4
6
8
10
−10
0
10
20
30
40

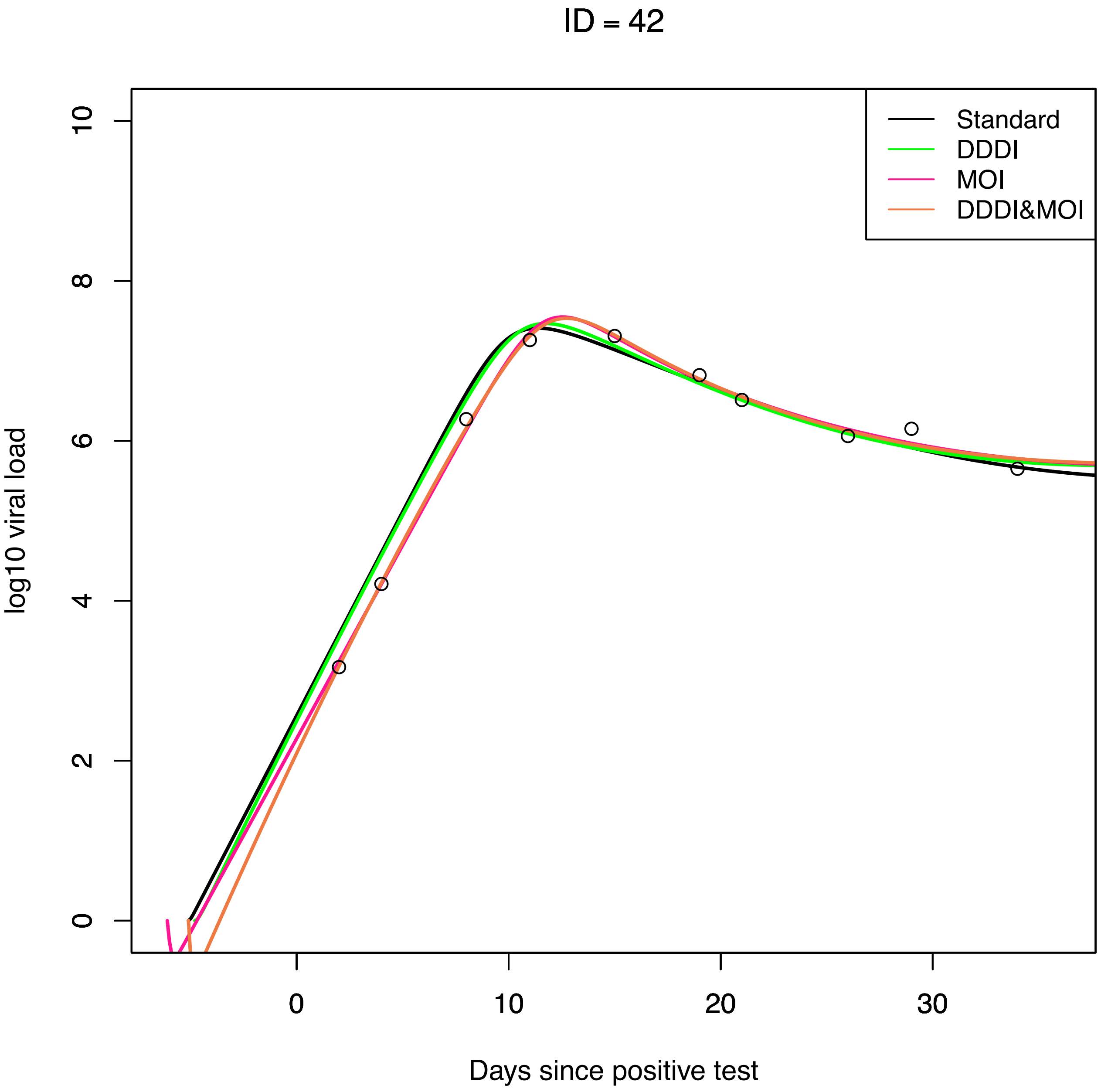

ID = 42
Standard
DDDI
MOI
DDDI&MOI
log10 viral load
0
2
4
6
8
10
0
10
20
30
Days since positive test

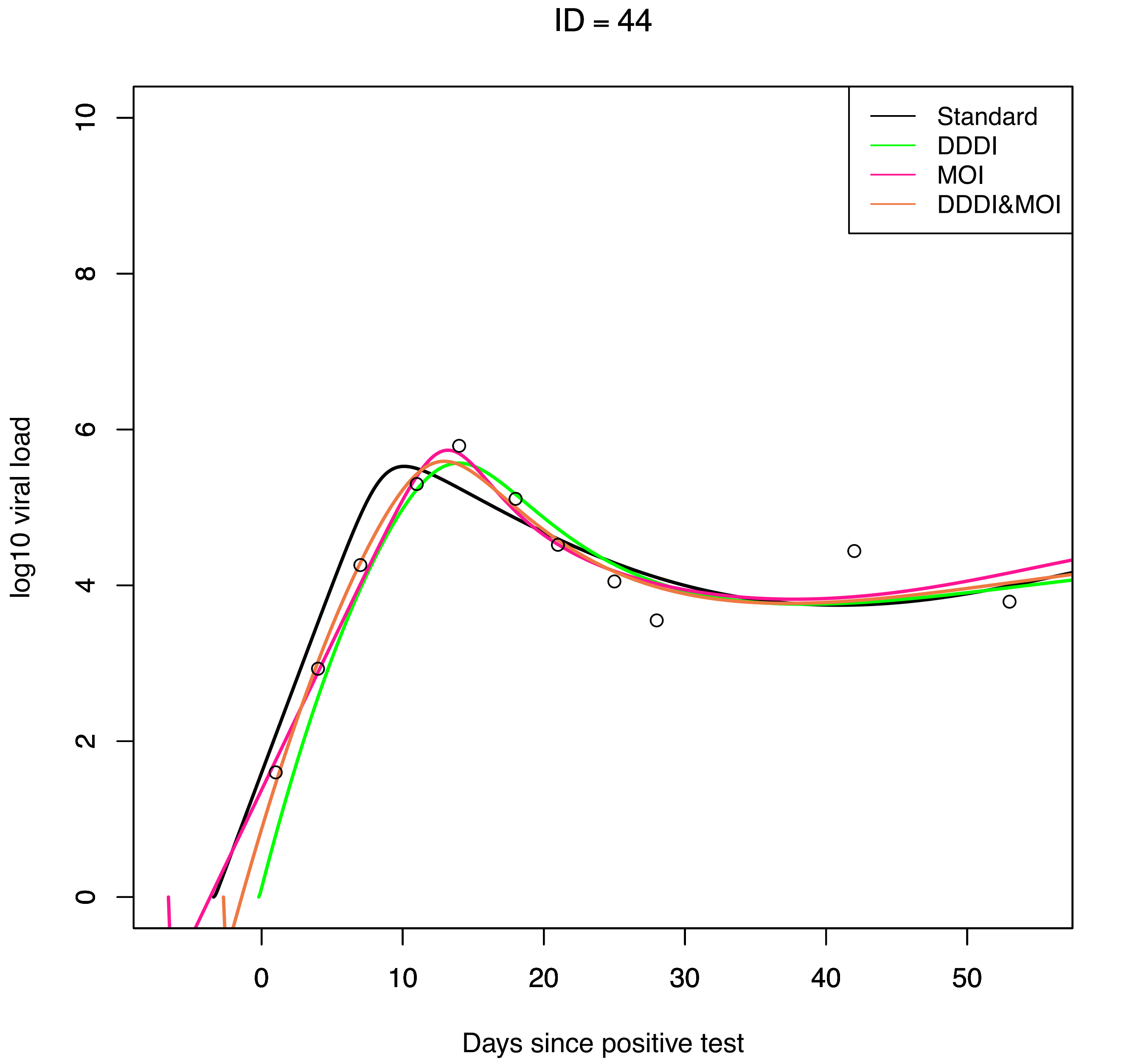

ID = 44
Standard
DDDI
MOI
DDDI&MOI
log10 viral load
Days since positive test
0
2
4
6
8
10
0
10
20
30
40
50

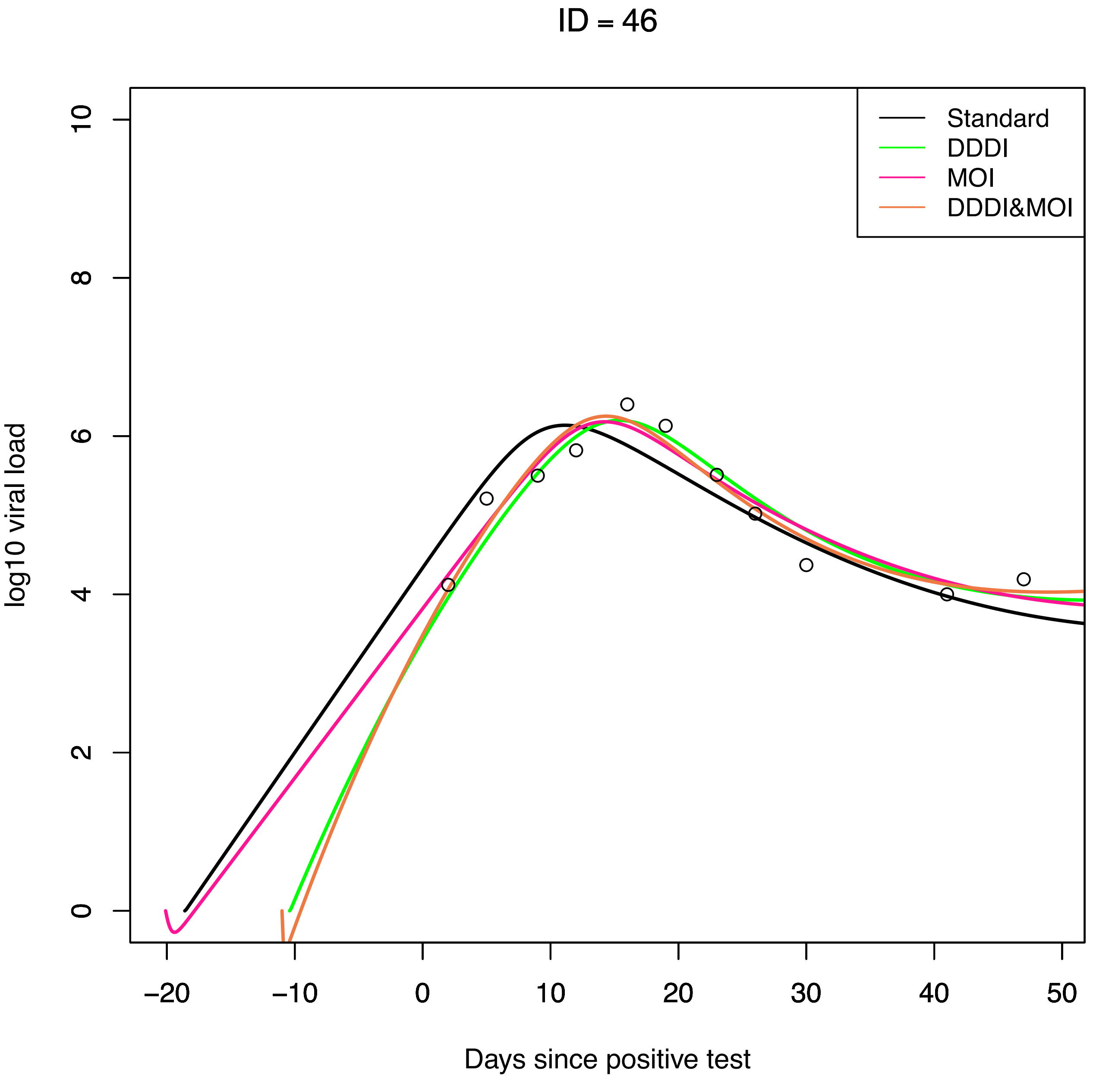
ID = 46
Standard
DDDI
MOI
DDDI&MOI
log10 viral load
0
2
4
6
8
10
−20
−10
0
10
20
30
40
50
Days since positive test

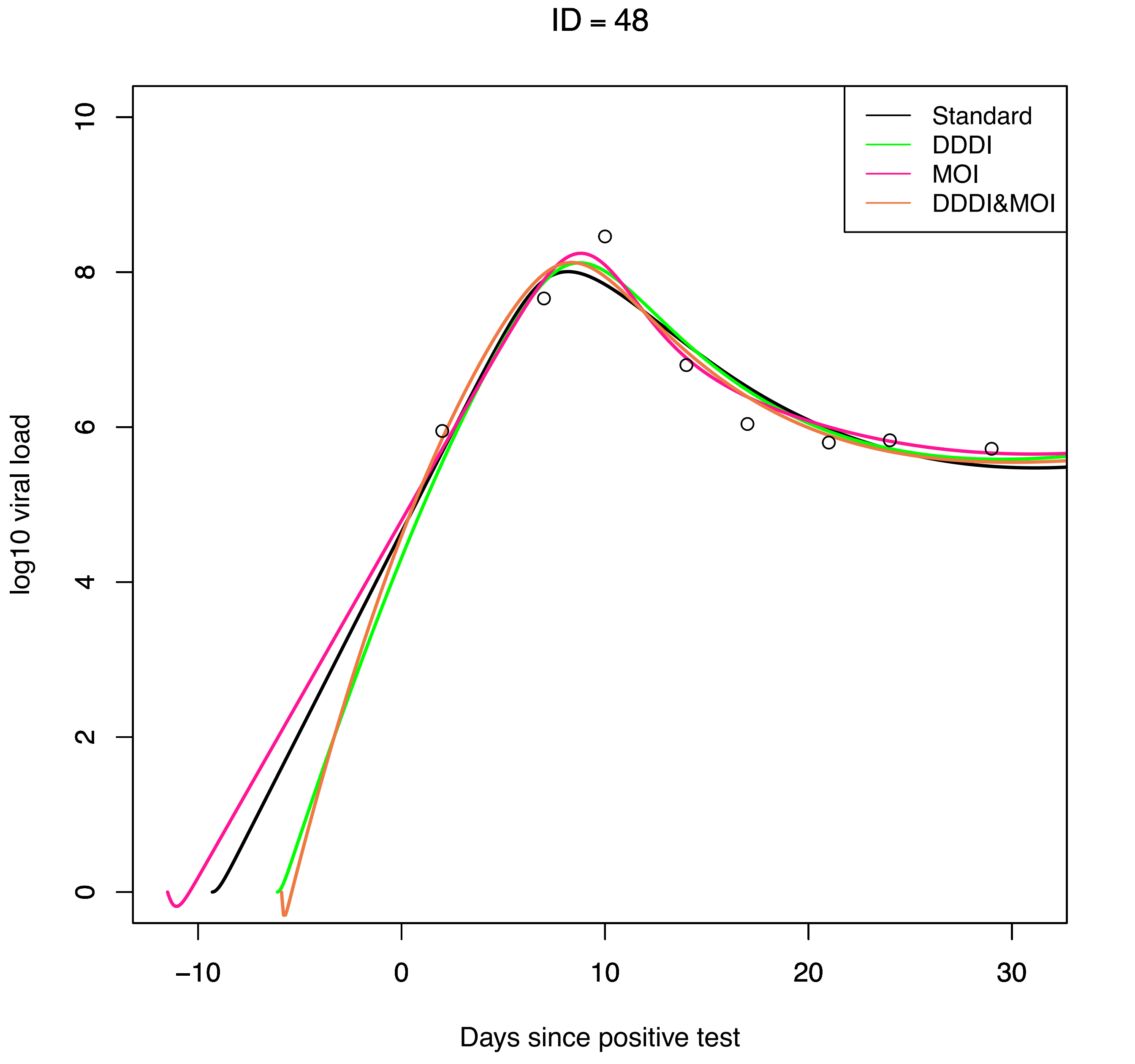

ID = 48
Standard
DDDI
MOI
DDDI&MOI
log10 viral load
Days since positive test
0
2
4
6
8
10
−10
0
10
20
30

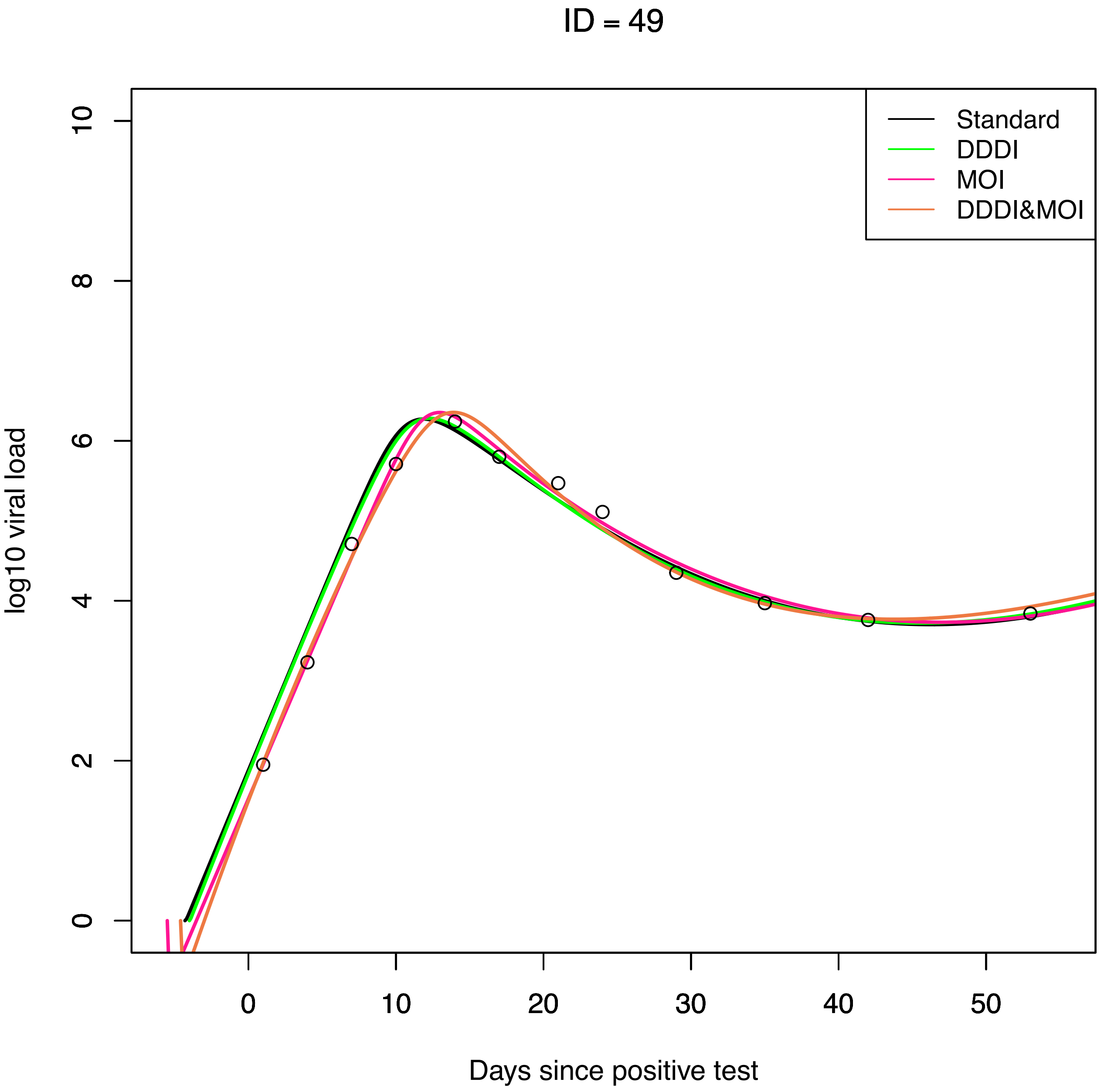

ID = 49
Standard
DDDI
MOI
DDDI&MOI
log10 viral load
0
2
4
6
8
10
0
10
20
30
40
50
Days since positive test

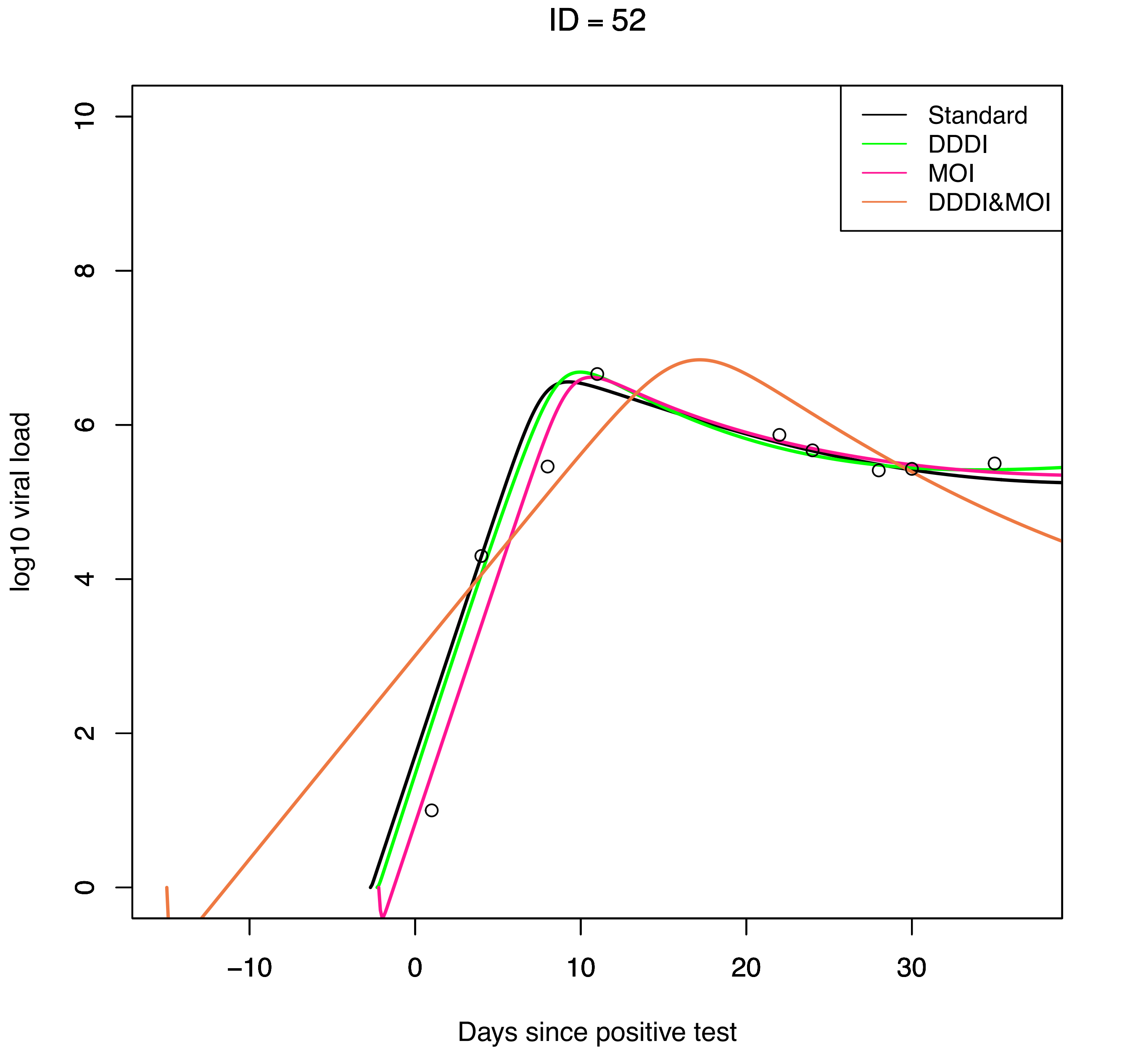

ID = 52
Standard
DDDI
MOI
DDDI&MOI
log10 viral load
0
2
4
6
8
10
−10
0
10
20
30
Days since positive test

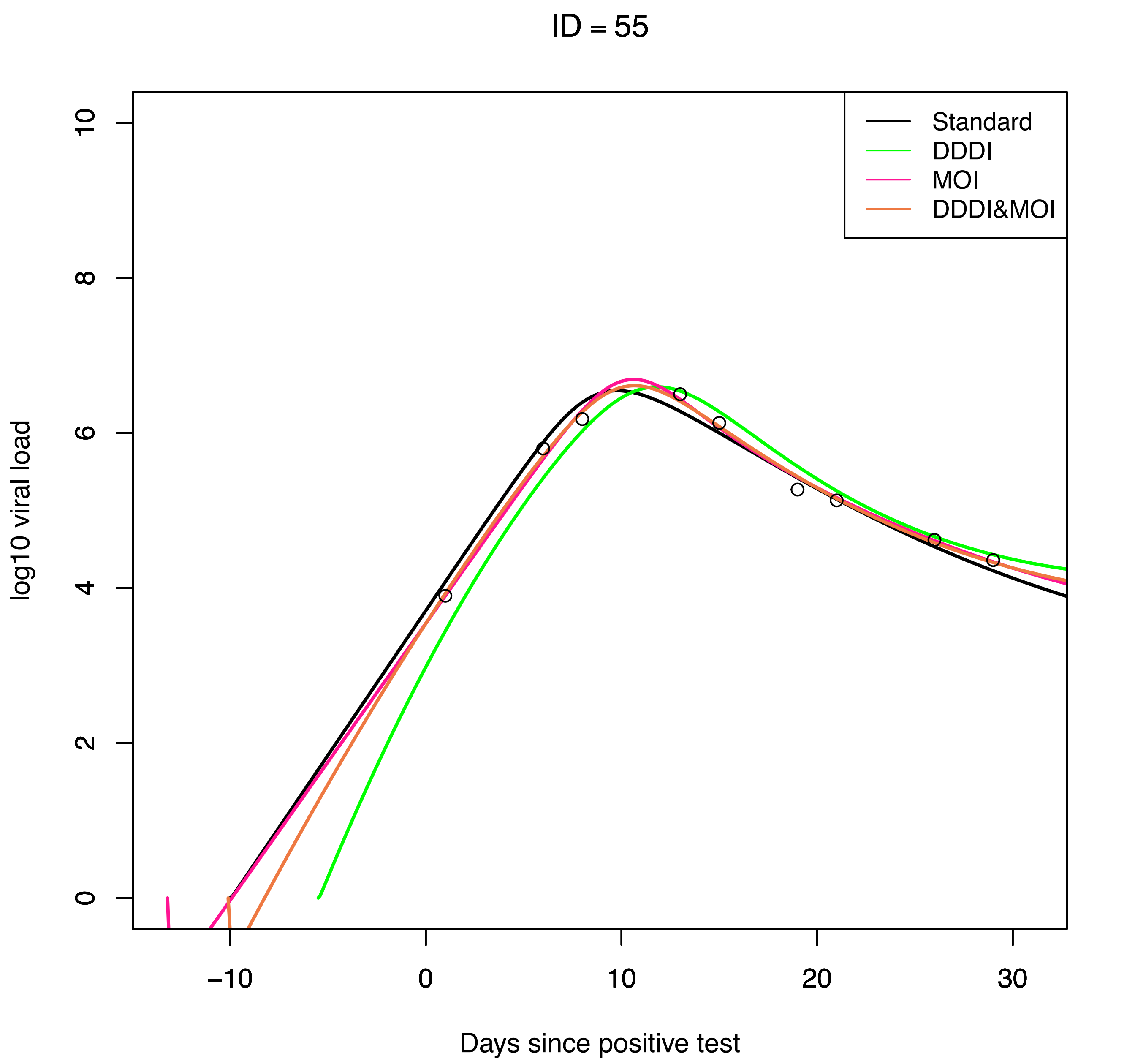

ID = 55
Standard
DDDI
MOI
DDDI&MOI
log10 viral load
0
2
4
6
8
10
−10
0
10
20
30
Days since positive test

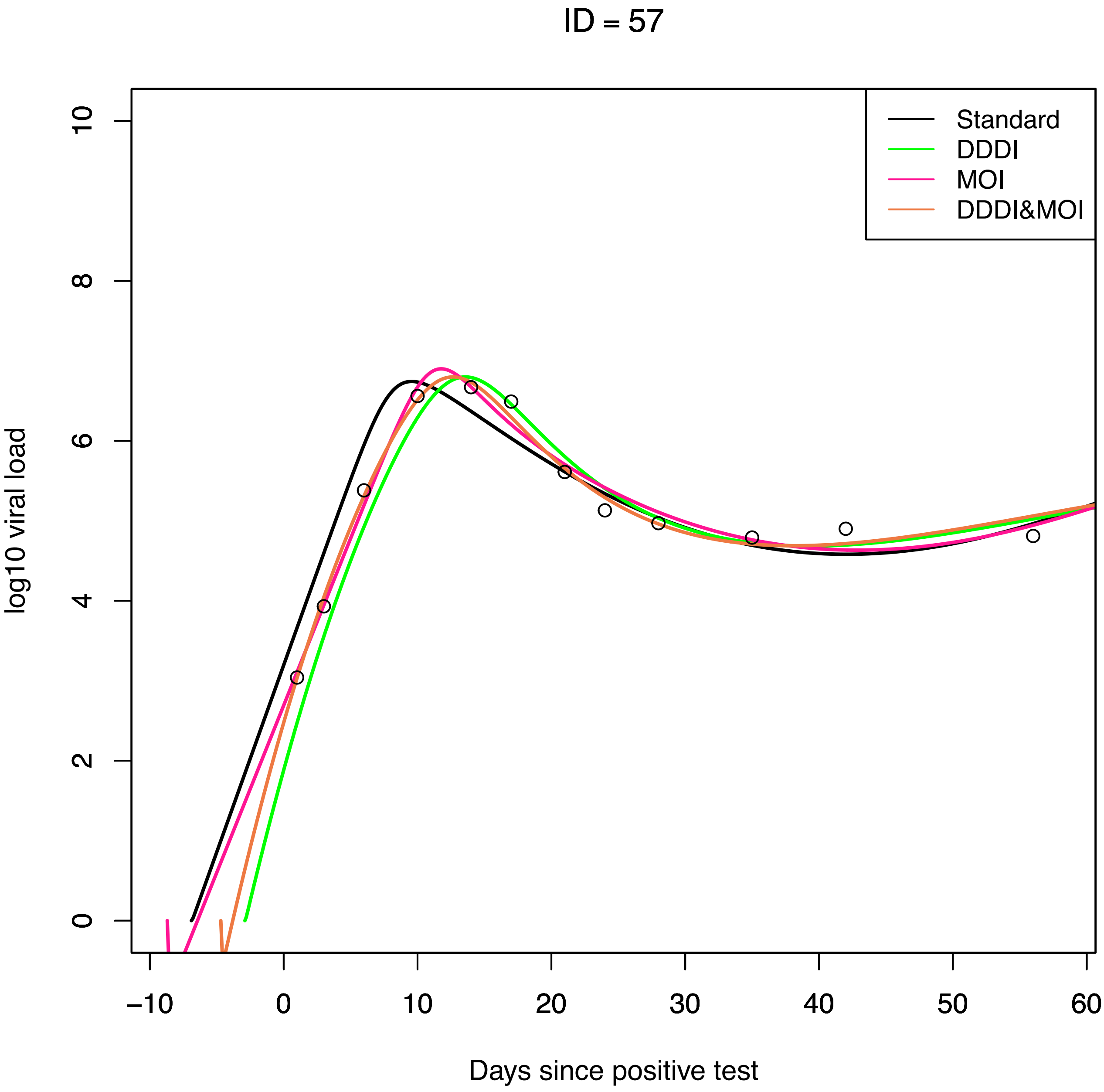
ID = 57
Standard
DDDI
MOI
DDDI&MOI
log10 viral load
0
2
4
6
8
10
−10
0
10
20
30
40
50
60
Days since positive test

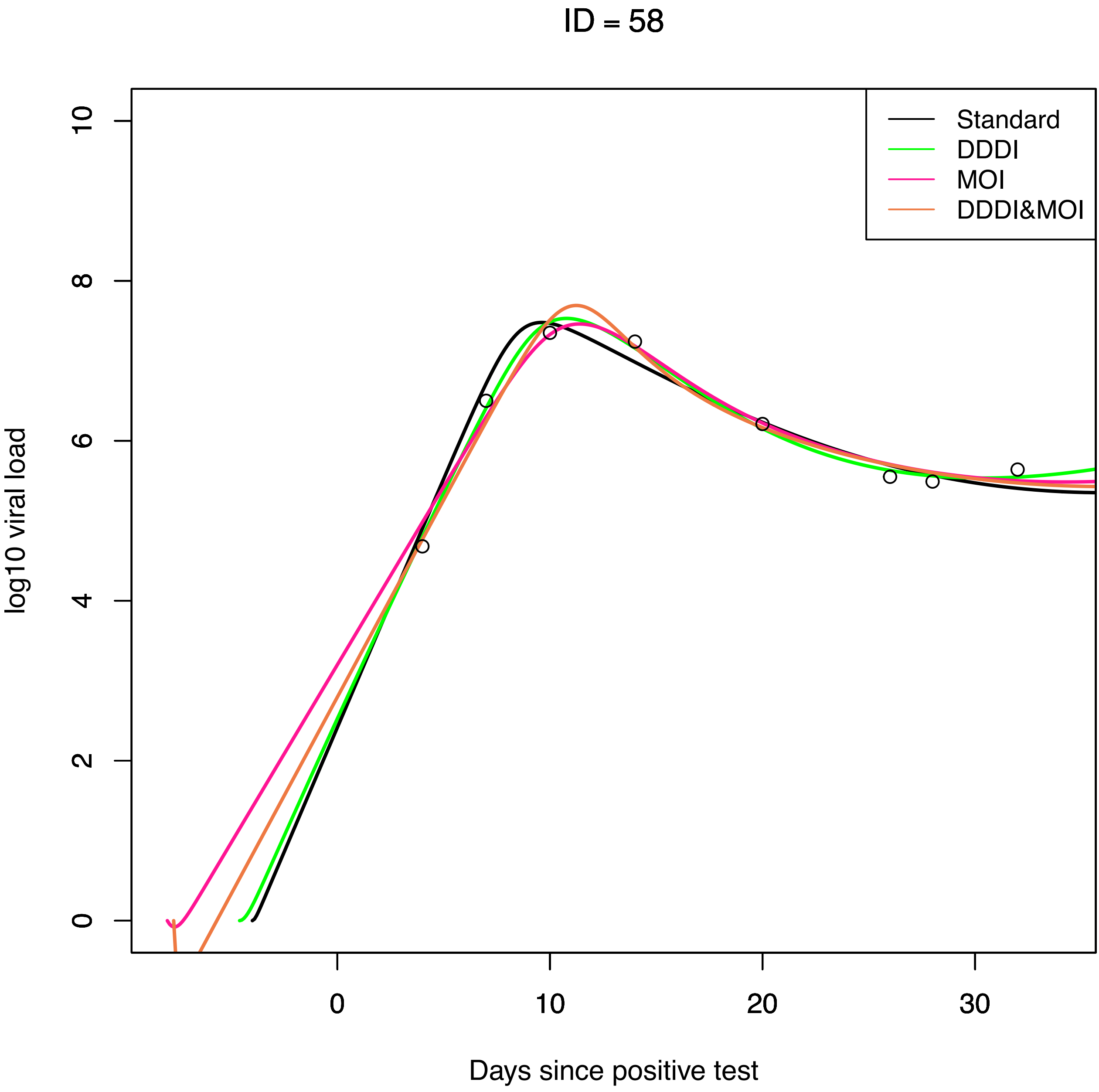

ID = 58
Standard
DDDI
MOI
DDDI&MOI
log10 viral load
0
2
4
6
8
10
0
10
20
30
Days since positive test

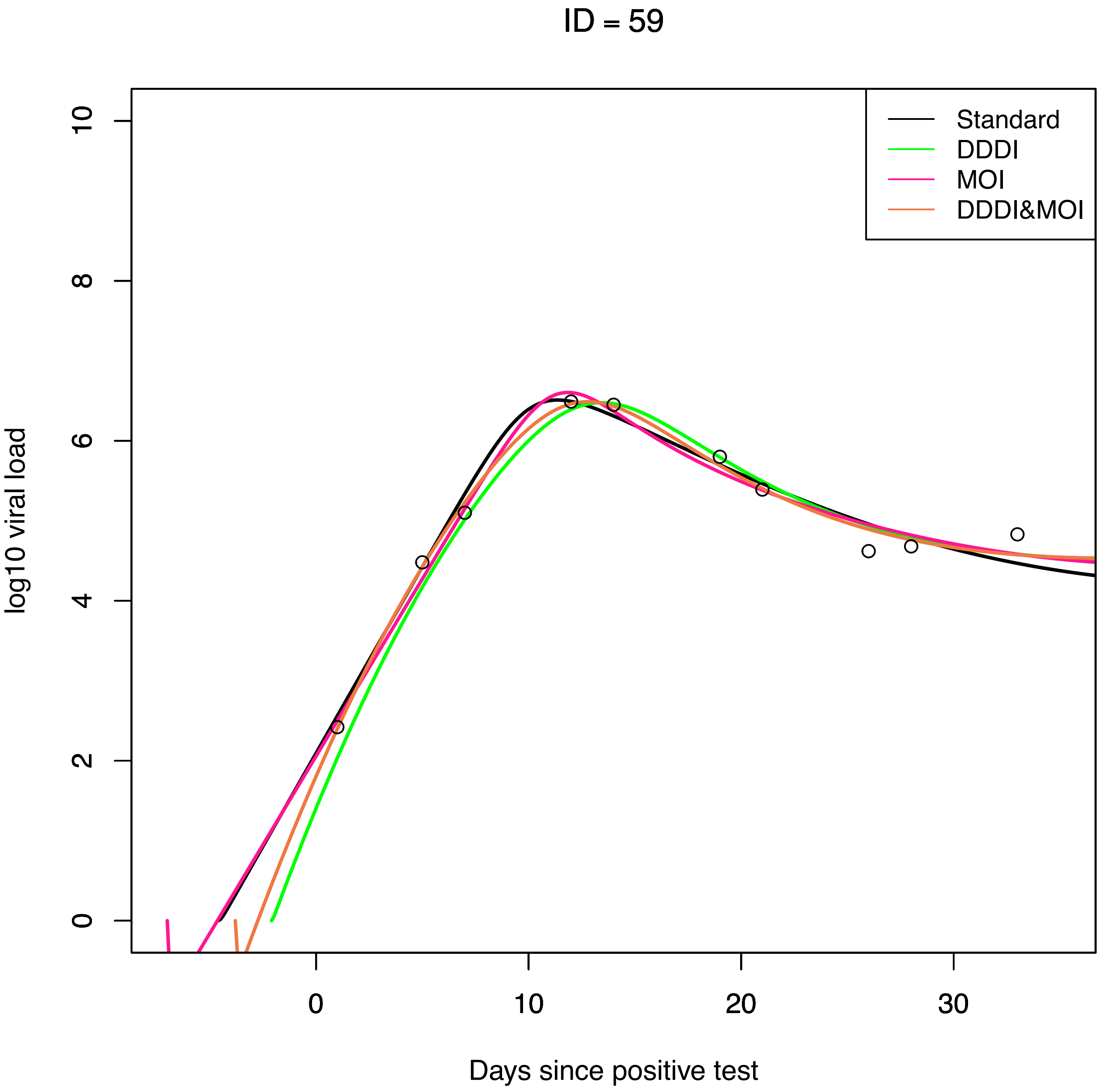
ID = 59
Standard
DDDI
MOI
DDDI&MOI
log10 viral load
0
2
4
6
8
10
0
10
20
30
Days since positive test

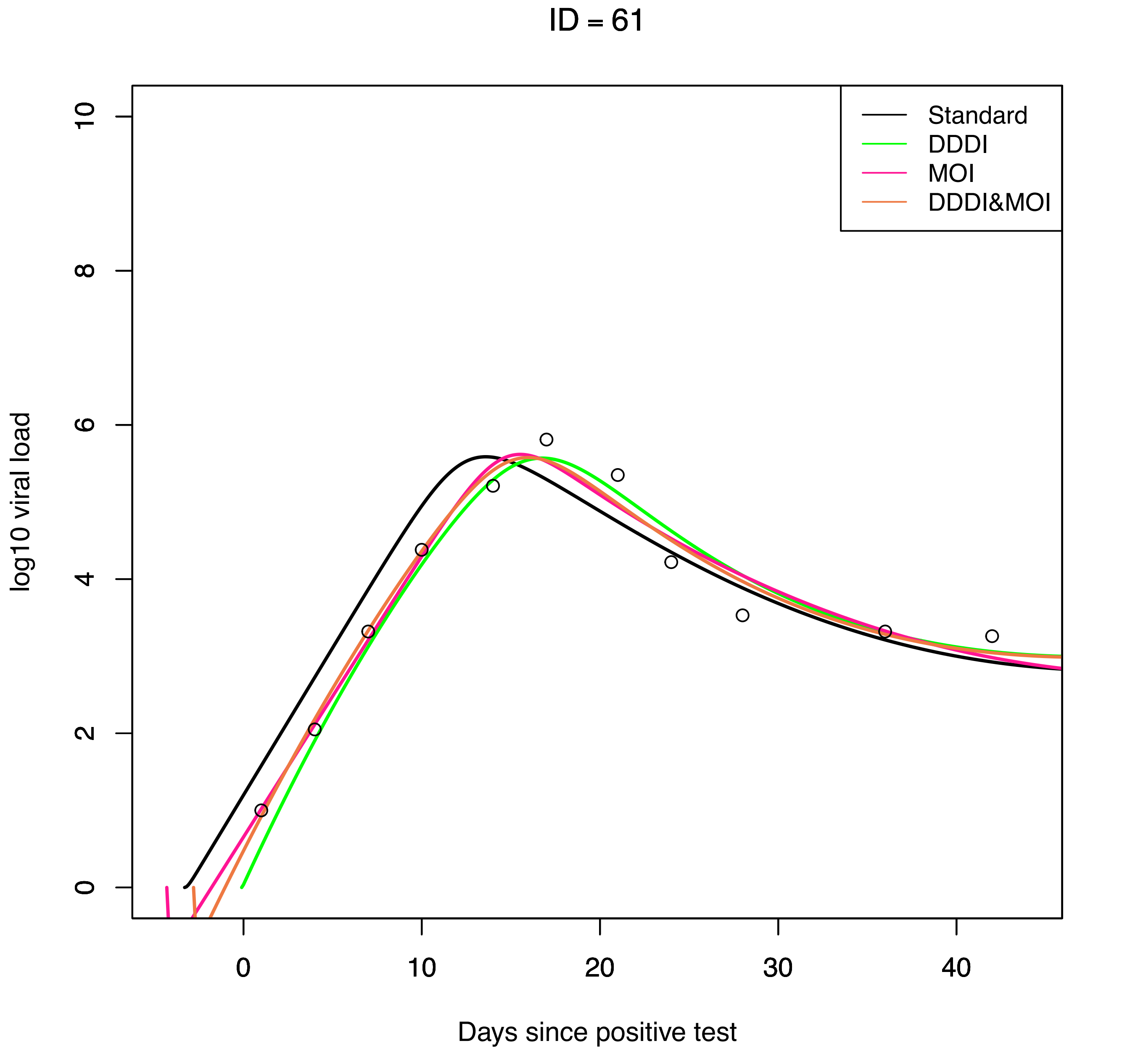

ID = 61
Standard
DDDI
MOI
DDDI&MOI
log10 viral load
Days since positive test
0
2
4
6
8
10
0
10
20
30
40

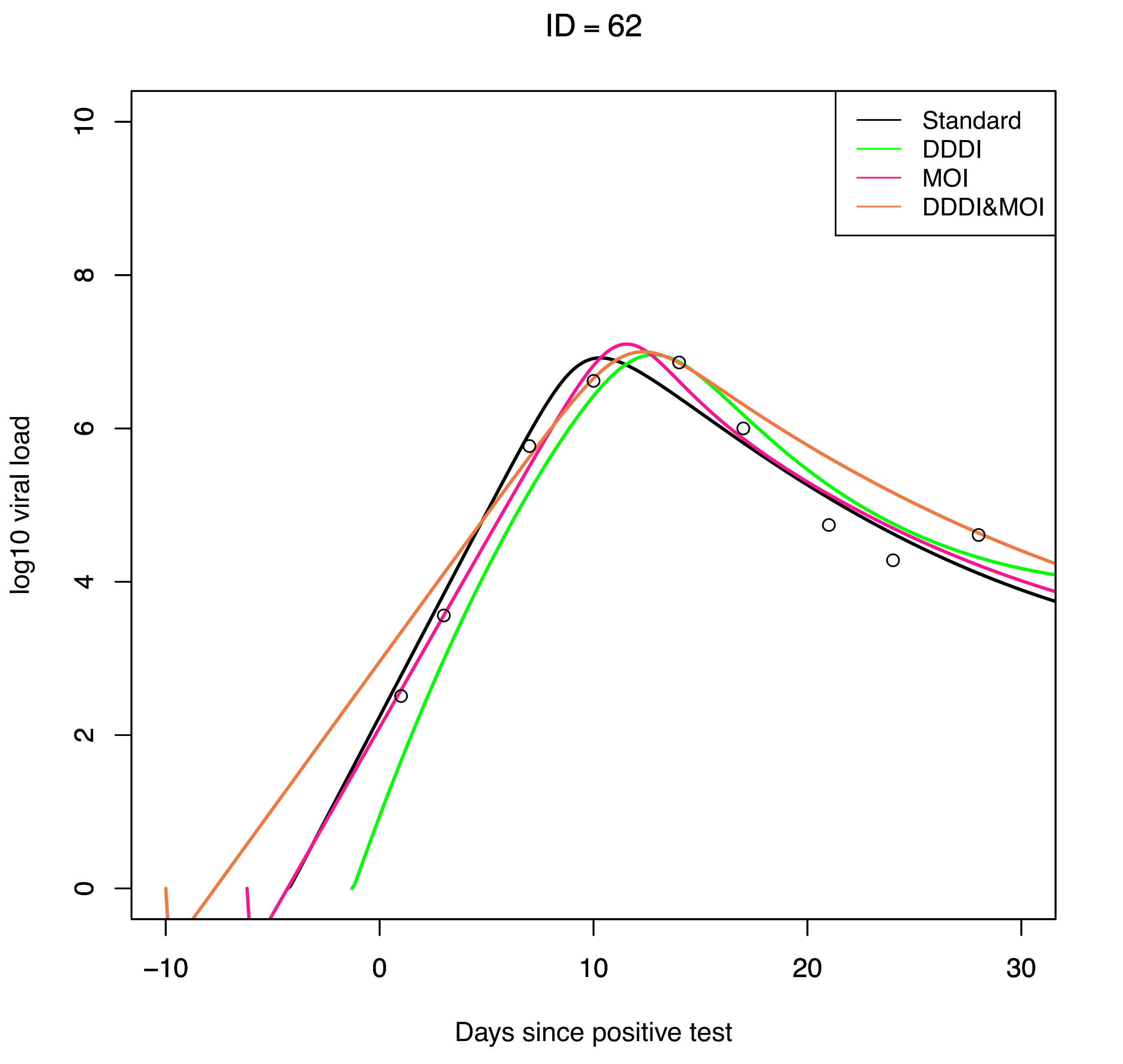

ID = 62
Standard
DDDI
MOI
DDDI&MOI
log10 viral load
0
2
4
6
8
10
−10
0
10
20
30
Days since positive test

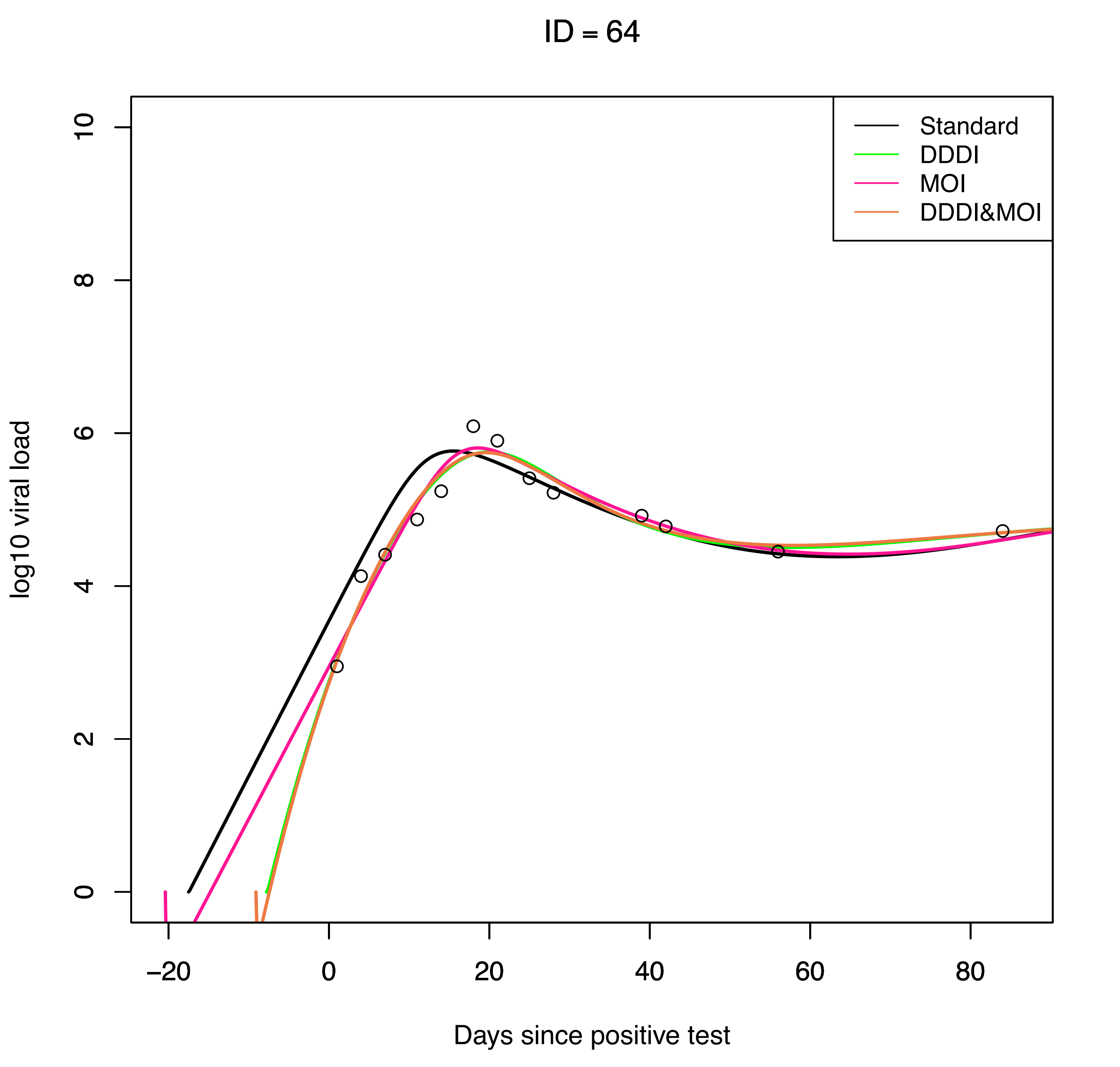

ID = 64
Standard
DDDI
MOI
DDDI&MOI
log10 viral load
0
2
4
6
8
10
−20
0
20
40
60
80
Days since positive test

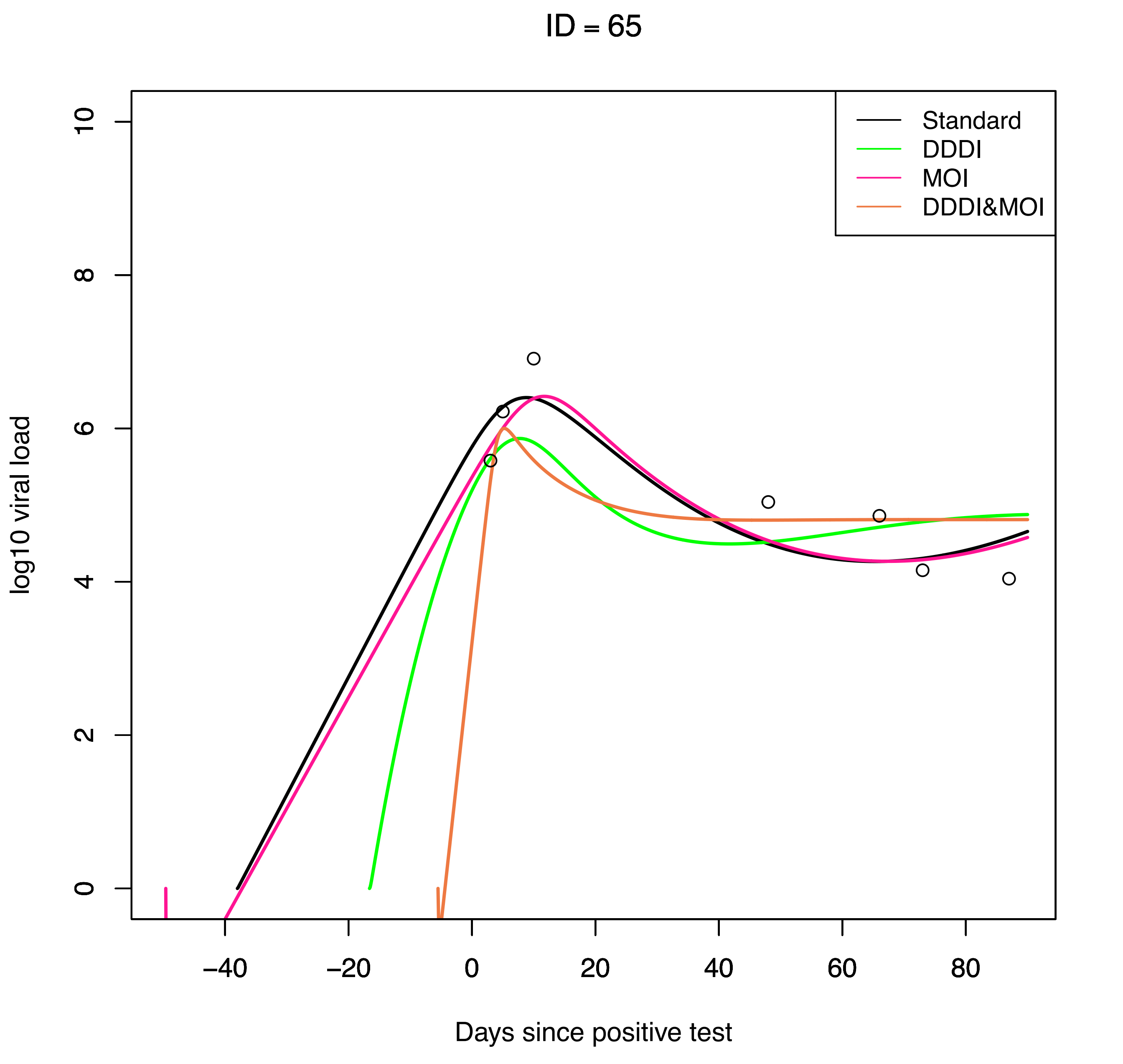
ID = 65
Standard
DDDI
MOI
DDDI&MOI
log10 viral load
0
2
4
6
8
10
−40
−20
0
20
40
60
80
Days since positive test

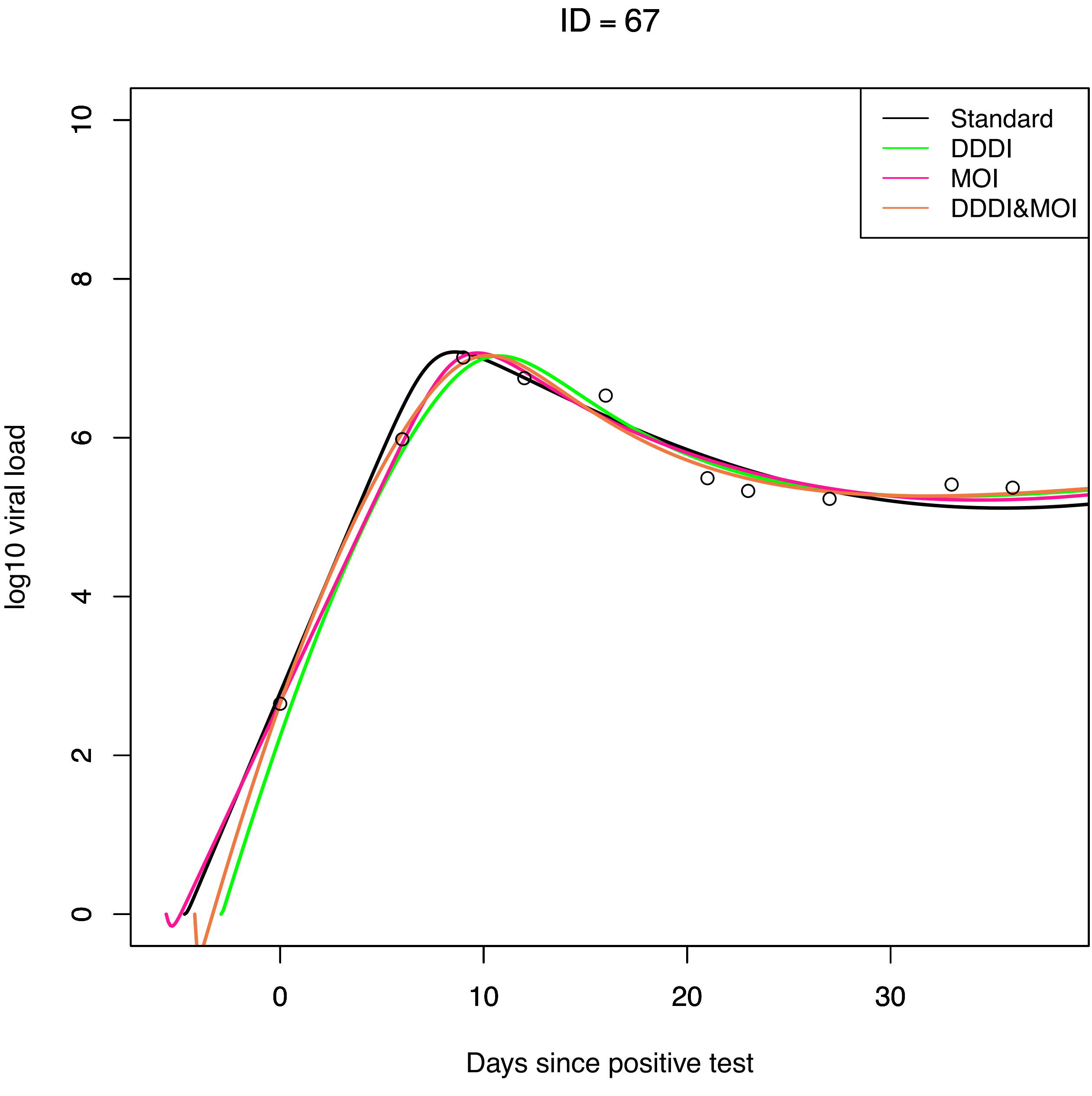
ID = 67
Standard
DDDI
MOI
DDDI&MOI
log10 viral load
0
2
4
6
8
10
0
10
20
30
Days since positive test

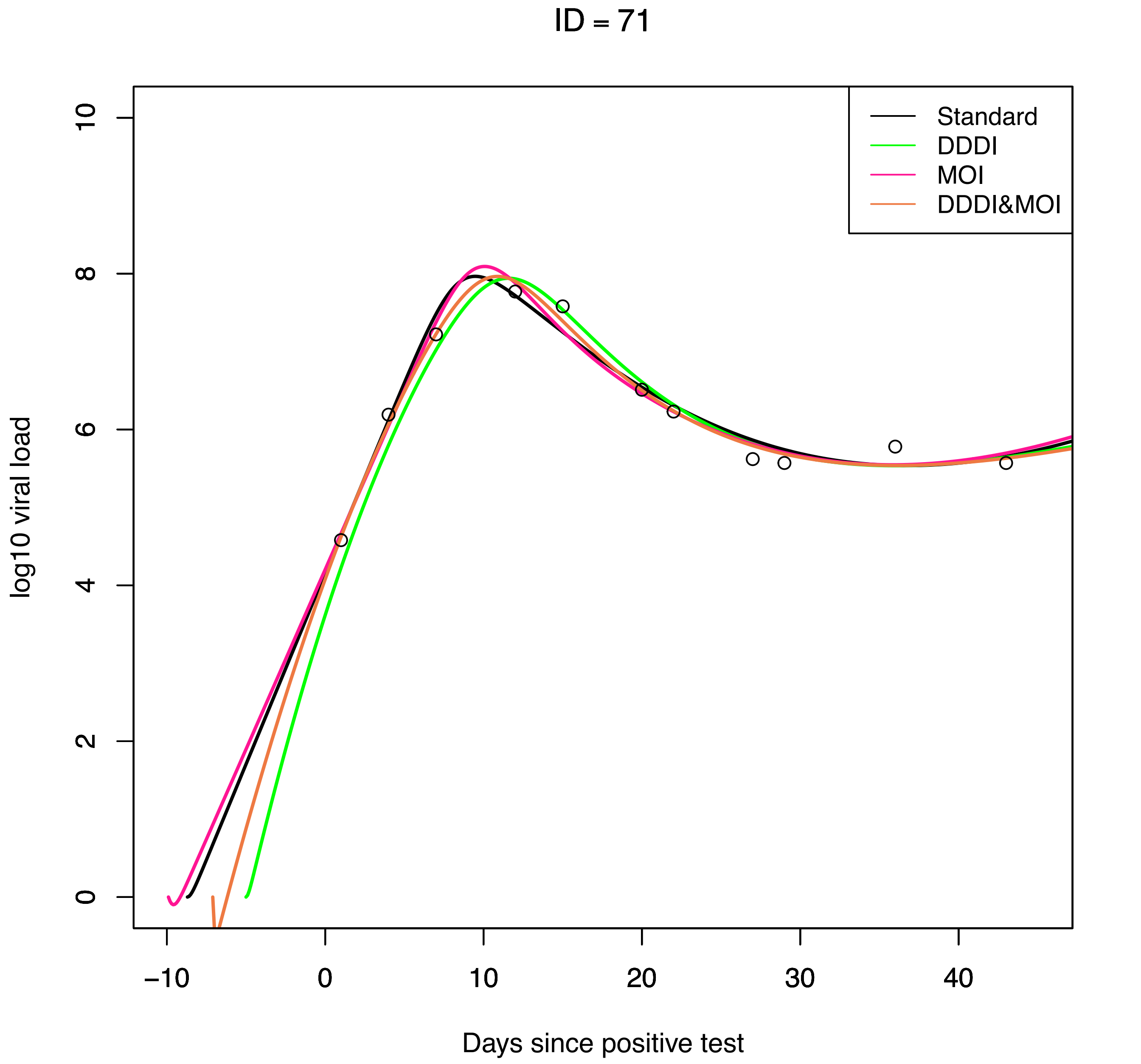
ID = 71
Standard
DDDI
MOI
DDDI&MOI
log10 viral load
0
2
4
6
8
10
−10
0
10
20
30
40
Days since positive test

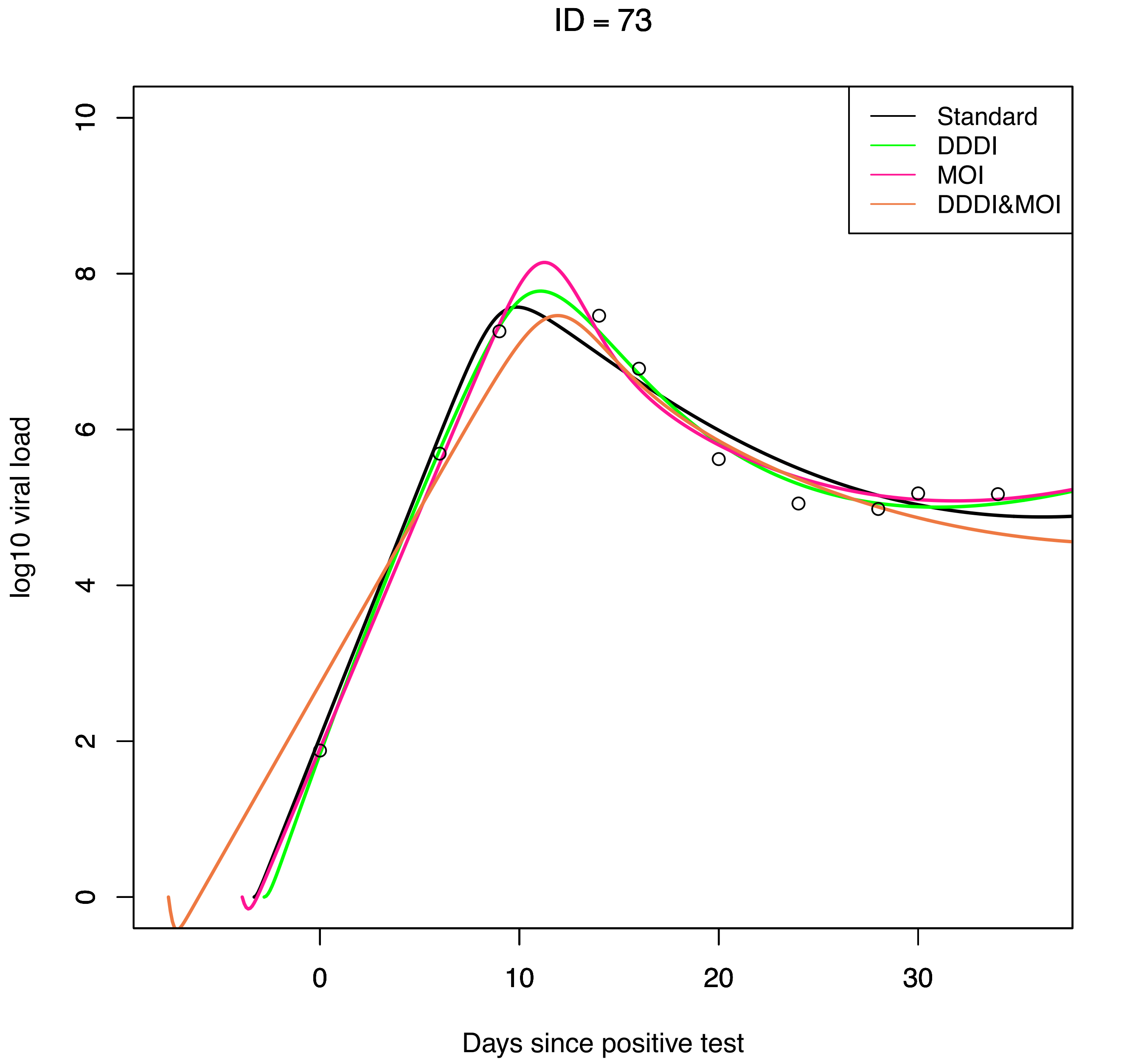

ID = 73
Standard
DDDI
MOI
DDDI&MOI
log10 viral load
0
2
4
6
8
10
0
10
20
30
Days since positive test
